# Optical control of CRISPR-Cas editing with cyclically caged guide RNAs

**DOI:** 10.1101/2022.03.04.482981

**Authors:** Ying-Jie Sun, Ji Liu, Jun-Jin Li, Yu Zhang, Wen-Da Chen, Wei-Qi Cai, Li Liu, Xin-Jing Tang, Jian Hou, Ming Wang, Liang Cheng

## Abstract

The CRISPR/Cas system has been proved as one of the most powerful tools for precise gene editing. However, the approaches for precise control over the genome editing and regulatory events are still desirable. Here, we reported a spatiotemporal and efficient CRISPR/Cas9 and Cpf1-mediated editing with photo-sensitive circular gRNAs. This approach relies on only two or three pre-installed photolabile substituents followed by a simple covalent cyclization, which provides a robust synthesize approach in comparison to heavily modified gRNAs. In established cells stably expressing Cas9, the circular gRNA in coordination with light irradiation could direct a precise cleavage of GFP and VEGFA within a pre-defined cutting region. We have also achieved light-mediated MSTN gene editing in embryos, whereas a new bow-knot-type gRNA showed no background editing in the absence of light irradiation. Together, our work provides a significantly improved method to precisely manipulate where and when genes are edited.

## Introduction

A temporal and spatial control of genome editing holds great potential for genetic disorder treatment and our capacity in investigating genetics variants. CRISPR-Cas, an acquired immune system of prokaryotic organisms in the process of continuous evolution to protect their genomes from interference and destruction by foreign nucleic acids,^1^ has proved a key advance for the easy and precise changing to the targeted genetic sequences. The fast development and wide application of Class II CRISPR systems are largely due to its simplicity by utilization of a single DNA endonuclease protein, like Cas9, Cpf1, and C2c1.^2^ An effective target recognition is propelled by complementarity pairing between the spacer sequence within the guide RNA (gRNA) with the target double-stranded DNA (dsDNA), followed by cleavage of the dsDNA substrates adjacent to the protospacer adjacent motif (PAM). Despite advances in precision, targeting range and applicability of CRISPR to therapeutics in recent years, tools to temporally and spatially control CRISPR editing are lagging behind. The editing proceeds freely in cells until the machinery, *i*.*e*., Cas endonuclease and guide RNA gRNA degrades through cellular processes. Uncontrolled CRISPR systems resulting from prolonged or unrestrained Cas endonuclease activity could have unpredictable off-target effects. To solve this problem, a variety of strategies to manipulate the expression of Cas protein or its activity have been proposed. In some cases, the endonuclease Cas9 production can be modulated by an inducer or be produced in a split or an inactive form, which could later be activated upon stimulation with small molecules or by exposure to biorthogonal triggers such as light.^3-6^

Meanwhile, different approaches have been established to regulate the expression level and duration of functional gRNA, the other indispensable component in the CRISPR machinery.^7-8^ Compared with the optimization of producing functional recombinant inducible Cas proteins, the modification of gRNAs (crRNA or single guide RNA) is simple and cost-effective. The standard phosphoramidite RNA chemistry has enabled the incorporation of versatile modified nucleotide motif in a site-specific manner, which can improve the stability of gRNAs towards ribonucleases and the specificity with Cas endonuclease.^9-16^ Among these approaches, photolabile groups,^17-27^ small-molecule-induced cleavable linkers,^26,28-30^ oligo-binding sites,^31-36^ and thermo-reversible substituents^37^ could be deliberately introduced which, in addition to magnify the duration of oligonucleotides, eliminated the genome editing activity of synthetic gRNAs that can later be switched on by external stimuli hence triggering the editing process. However, a high number of modifying sites, usually one in every 5∽6 nucleotides,^22,29^ were required to hinder the formation of gRNA-DNA heteroduplex during R-loop expansion and destabilize the conformational flexibility of gRNA:DNA:Cas9 complex.^38^ Insufficient caging groups would lead to partially accessable to the target DNA, resulting in undesired background editing before stimuli treatment. Nonetheless, an efficient decaging with high intensities of light exposure, long irradiation time or high concentrations of small molecule triggers was always required for multiply modified gRNAs to achieve an ideal ON/OFF-switch control. An incomplete removal of the caging groups with inferior editing efficiency would be a limitation for its application to spatially and temporally control CRISPR editing *in vitro* or *in vivo*.^39-44^

Recognizing that the gRNA is the most easily programmable component of the CRISPR system, we set out to investigate gRNAs that could be readily constructed with minimal modification with a complete abolishment of its recruitment to the target genomic region and yet speedily activated in the presence of light.^21,45-50^ In this article, we introduce a new controllable CRISPR/Cas editing system with covalently head-to-tail cyclized gRNAs involving a light-sensitive linkage that caged the RNA (Fig. 1A). As opposed to traditional strategy of preparing linear gRNA with multiple photolabile groups, the construction of the photoactivatable circular gRNA requires only two or three pre-installed buckles for each sequence, which provides a robust synthesize approach while still maintaining high stability against degradation by nucleases. Subsequent light irradiation enables a rapid release of the native gRNA and imitate the gene editing process in live cells within only a few minutes. The method is demonstrated on green fluorescent protein (GFP) and vascular endothelial growth factor A (VEGFA), respectively. We also take advantage of this technique to facilitate the successful disruption of the myostatin (MSTN) in mouse embryos by gene editing.

**Fig. 1.**
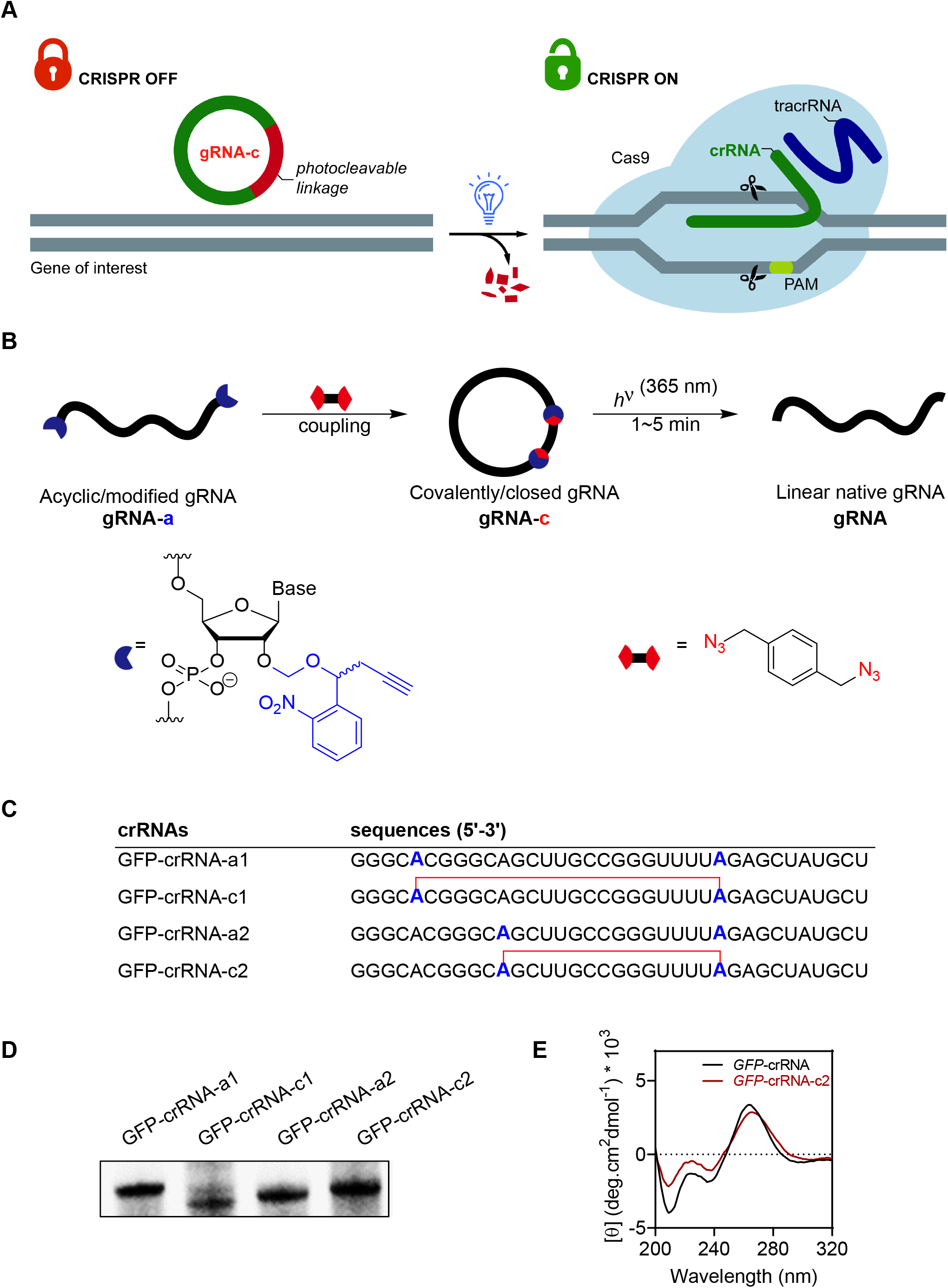
Chemistry and design of the circular crRNAs. **A**. Schematic elucidation of the circular gRNA principle for the spatiotemporal control of CRISPR/Cas editing. **B**. Strategy of constructing the circular gRNA (gRNA-c) through a biorthogonal condensation of di-modified acyclic gRNA (gRNA-a) with bidentate azide component (two red fan-shaped sectors connected by a black linkage), and subsequent photo-dissociation under light irradiation within 1 to 5 minutes to release the linear native gRNA. **C**. Sequences of prepared acyclic NPBOM modified and cyclized crRNAs targeting GFP sequence. Blue bold letters indicate the modification sites substituted with NPBOM, and the red lines indicate the NPBOM derived linkage upon reacting with 1,4-bis(azidomethyl)benzene. **D**. Polyacrylamide gel electrophoresis analysis of the linear GFP-crRNA-a and circular GFP-crRNA-c. **E**. Circular dichroism spectroscopy of native GFP-crRNA (black line) and its circular counterpart GFP-crRNA-c2 (red line).

## Results and Discussion

### Design, synthesis and characterization of photoactivatable circular crRNA-c

The mechanism of photocaged RNA is based on the installation of a photocleavable protecting group onto the nucleotides that renders it inactive, either by halting the formation of ribonucleoprotein (RNP) complex with Cas proteins and/or binding to the target sequence in the genomic DNA. This strategy has already been applied in many examples where the photolabile functionalities were indiscriminately distributed among the spacer and scaffold regions of short crRNAs (36-43 nt).^19,22-25,27^ However, methodology that allows for the preparation of longer gRNAs of high-purity with over 100 nucleotides containing dozens of site-specifically incorporated photocleavable functionalities has been documented as a limitation. We believe that the anchoring of two modified nucleotides at the 5 - and 3 -end respectively, followed by an intramolecular covalent cyclization similar to the head-to tail splice junction of its natural counterpart, the circRNAs,^51^ would form a closed loop with a rigid conformation but still could be fragmented in the presence of light (Fig. 1B). For this, we added the *o*-nitrobenzyloxylmethoxyl group containing a terminal propyne handler (NPBOM), which are sensitive to cleavage when exposed to ultraviolet light, at two arbitrarily selected adenosines located at the spacer and duplex regions on the crRNAs targeting GFP (Fig. 1C, Fig. S1-2). The decorated crRNAs (GFP-crRNA-a1 and GFP-crRNA-a2) with two terminal alkyne residues were then subjected to the copper-catalyzed azide/alkyne cycloaddition reaction (CuAAC) with the bifunctional 1,4-bis(azidomethyl)benzene (Fig. S5-7).^52-54^ It was reported that the cyclization of oligonucleotides often was accompanied by dimerization or cyclodimerization as the ring size varied.^55-57^ Hence, these macrocyclizations usually had to be performed under high dilution conditions. Interestingly, it was found that the covalent condensation proceeded smoothly at 75 μM with almost same rates despite the different ring sizes of these two crRNAs (22 nt and 16 nt, respectively) (Fig. 1D). On the basis of the agarose gel electrophoresis analysis and matrix-assisted laser desorption/ionization-time of flight (MALDI-TOF) mass results, the structures of two circular crRNAs GFP-crRNA-c1 and GFP-crRNA-c2 were confirmed *via* an intramolecular process (Fig. 1D, Fig. S21, S23). The topological change between linear GFP-crRNA and circular GFP-crRNA-c2 was also evaluated directly using circular dichroism (CD) spectroscopy (Fig. 1E).^58^ The ellipticity changes around 210 nm and 225 nm of the CD spectra, due to lone pair (lp)-π* stacking interactions in the nucleobases, was indicative of backbone alterations (cyclization) to the GFP-crRNA-c2.^59^ It also correlated with the slight red shift and the intensity decrease of the peak around 264 nm, which indicated different geometry of those two crRNAs and in consequence different inter- and intra-strand hydrogen bonds and base stacking. All those results proven that the head-to-tail cyclization has originated significant deviations to the secondary and/or tertiary structure.^60-61^

### *In vitro* photomodulation of DNA cleavage with circular crRNAs

To demonstrate that the DNA interference complex in CRISPR-Cas9 systems was dysfunctional with circular crRNA due to the incapacity in recognizing target site or recruiting Cas9 for DNA cleavage, the melting characteristics of the circular GFP-crRNA-c2 and its two linear precursors were investigated (Fig. 2A, Fig. S8). It was shown by comparison that the covalent linkage resulted a significant decrease in melting temperature (T_m_) comparing to the linear crRNAs, indicating its inability to bind with the template DNA. Again, consistent with that, the circular crRNA-c2 was not capable of assembling the ternary dCas9-crRNA-tracrRNA complex when a 3’-FAM dsDNA was used (Fig. 2B), demonstrating that the circular nature of head-to-tail spliced crRNA confers its inertia towards interacting with other biomacromolecules. Indeed, one of the most attractive features of crRNA-c is its extreme stability against endonuclease degradation (Fig. S9). *In vitro* assay of circular GFP-crRNA-c for RNase A stability showed very little degradation after 60 minutes’ incubation. By contrast, prototype GFP-crRNA or GFP-crRNA-a were rapidly degraded within 1 minute (Fig. S9). However, the stable circular GFP-crRNA-c released the linear crRNA within 30 seconds upon exposure with a 365nm UV light of an irradiance at 20 mW·cm^−2^ (Fig. 2C). Consequently, when the mixture with caged crRNA-c was subjected to light irradiation, a considerable amount of dCas9-crRNA-tracrRNA was detected (Fig. 2B). These results correlated with the above photo-inducible release of linear crRNA and the low dose and extremely short exposure to UV radiation for the cleavage of cyclized NPBOM-linkage minimized any potential damage to cells’ DNA and hence provided an excellent response window for the responsive CRISPR editing.

**Fig. 2.**
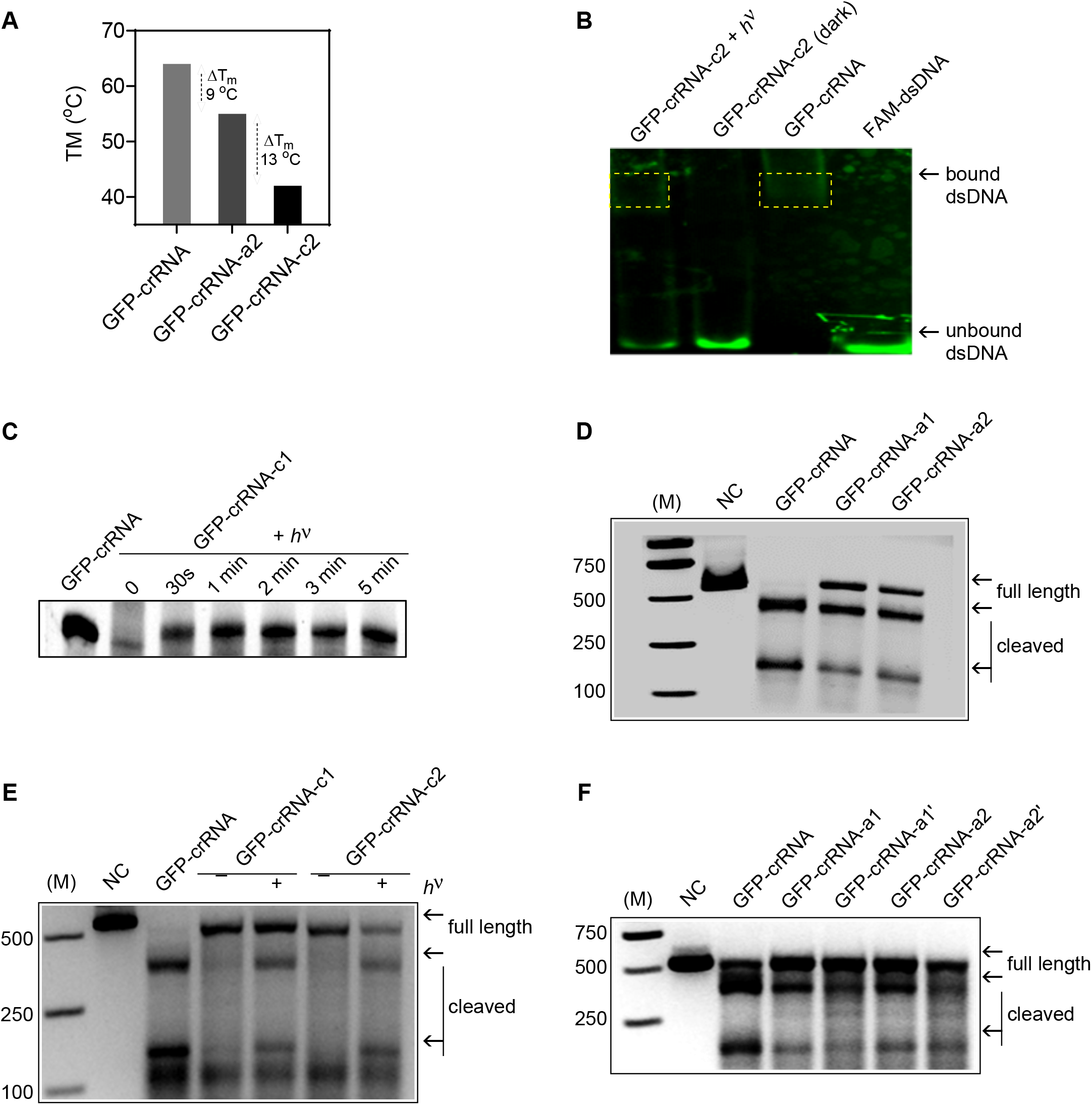
Light-induced CRISPR-Cas9-mediated DNA cleavage of GFP *in vitro*. **A**. Comparison of different melting temperatures among native GFP-crRNA, acyclic GFP-crRNA-a and the circular GFP-crRNA-c as determined by circular dichroism spectroscopy. **B**. Determination of assembling the ternary dCas9-crRNA-tracrRNA complex with native GFP-crRNA or photocaged GFP-crRNA-c2 with and without 365 nm light illumination (5 minutes, 20 mW·cm^−2^). **C**. Representative gel images of cleavage detection assay of the target GFP gene *in vitro* with native GFP-crRNA, acyclic GFP-crRNA-a1/a2. NC = Negative control. **D**. Comparison of native GFP-crRNA and light-mediated uncaging reaction of GFP-crRNA-c2 monitored by gel images at various time points. **E**. Representative gel images of cleavage detection assay of the target GFP gene *in vitro* with native GFP-crRNA, circular GFP-crRNA-c1/c2 with and without light irradiation (5 minutes, 20 mW·cm^−2^). **F**. Representative gel images of cleavage detection assay of the target GFP gene *in vitro* with native GFP-crRNA, acyclic GFP-crRNA-a1/a2 and their analogues GFP-crRNA-a1’/a2’.

We further investigated whether this photo-activable circular crRNA could restore the Cas9 cleavage ability *in vitro*. While the native GFP-crRNA induced Cas9 target DNA cleavage at a given locus, the dialkyne decorated linear GFP-crRNA-a1 and GFP-crRNA-a2 exhibited similar editing ability and resulted in considerable hydrolysis of the target DNA, indicating that the di-modified gRNA could not impede the cleavage (Fig. 2D). On the contrary, when the circular GFP-crRNA-c were used in the absence of light exposure, a complete inhibition of DNA cleavage was observed (Fig. 2E, Fig. S10). Subsequent photo-irradiation towards GFP-crRNA-c1 and GFP-crRNA-c2 both considerably restored the Cas9 nuclease activity to create a double-strand break (Fig. 2E, Fig. S10). Therefore, no leakage editing in the dark and precisely release at targeted locations by external light stimulation made it highly possible for accurate spatial and temporal OFF to ON switch of CRISPR gene editing. In order to assess that the resulting inactivity of editing with circular crRNA-c was due to the closed-loop structure, not from the conformational change simply arising from the two C2’-*O*-substituents within the ribose, the linear GFP-crRNA-a was subjected to react with monofunctional benzylazide under similar CuAAC conditions to afford two linear and di-modified crRNAs: GFP-crRNA-a1’ and GFP-crRNA-a2’, respectively (Fig. 2F, Fig. S11). Notably, while those two crRNAs contained similar triazolmethyl fragments at the same positions as that in GFP-crRNA-**c** were used, no observable inhibition of Cas9 induced cleavage was found, suggesting the irreplaceable role of the cyclic topology in the circular crRNA-**c** for the complete inactivation of crRNA function (Fig. 2F).

### *In vitro* photomodulation of CRISPR-Cpf1 editing with circular crRNAs

Comparing with previous Cas9 involved editing, the CRISPR/Cpf1 system has recently attracted much attention as a fascinating tool for gene editing and genome engineering.^62^ The endonuclease Cpf1 is smaller in size comparing with Cas9. It does not require tracrRNA, but is directed by a single guide RNA (sgRNA) that binds upstream of the PAM and cleaves the DNA at the proximal end far away from the seed region. We next examined whether the circular gRNA strategy could be extended to the long sequence sgRNA (43 nt) and thus hinder the Cpf1 cleavage. To this end, the sgRNA targeting the GFP locus with two adenosines at the spacer segment and the repeat handler were substituted with NPBOM accordingly, followed by similar CuAAC reaction (Fig. 3A, B). MALDI-TOF mass result and agarose gel electrophoresis analysis of GFP-Cpf1-sgRNA-c1 confirmed its structure as a cyclic sgRNA with larger ring size (Fig. S26). As expected, the linear GFP-Cpf1-sgRNA and GFP-Cpf1-sgRNA-a1 exhibited equal Cpf1 cleavage activity (Fig. 3C), meaning the efficiency of Cpf1-sgRNA was not influenced by terminal masking with NPBOM. It was presumed that large cyclic oligonucleotides might adopt balanced conformational flexibility/rigidity for the capability of binding to the target with close affinity and specificity,^63^ and here the circular 43-nt-ring sgRNA may reserve some flat regions that pair with the target DNA sequence or being recognized by Cpf1 protein. To our delight, the inducing activity of circular GFP-Cpf1-sgRNA-c1 was totally extinguished in the absence of light (Fig. 3D), which again proven that the head-to-tail closure may be a generic strategy for the modulation of gRNA function in gene editing.^64-65^ We further investigated whether a rapid exposure with UV light would restore the Cpf1 cleavage.

**Fig. 3.**
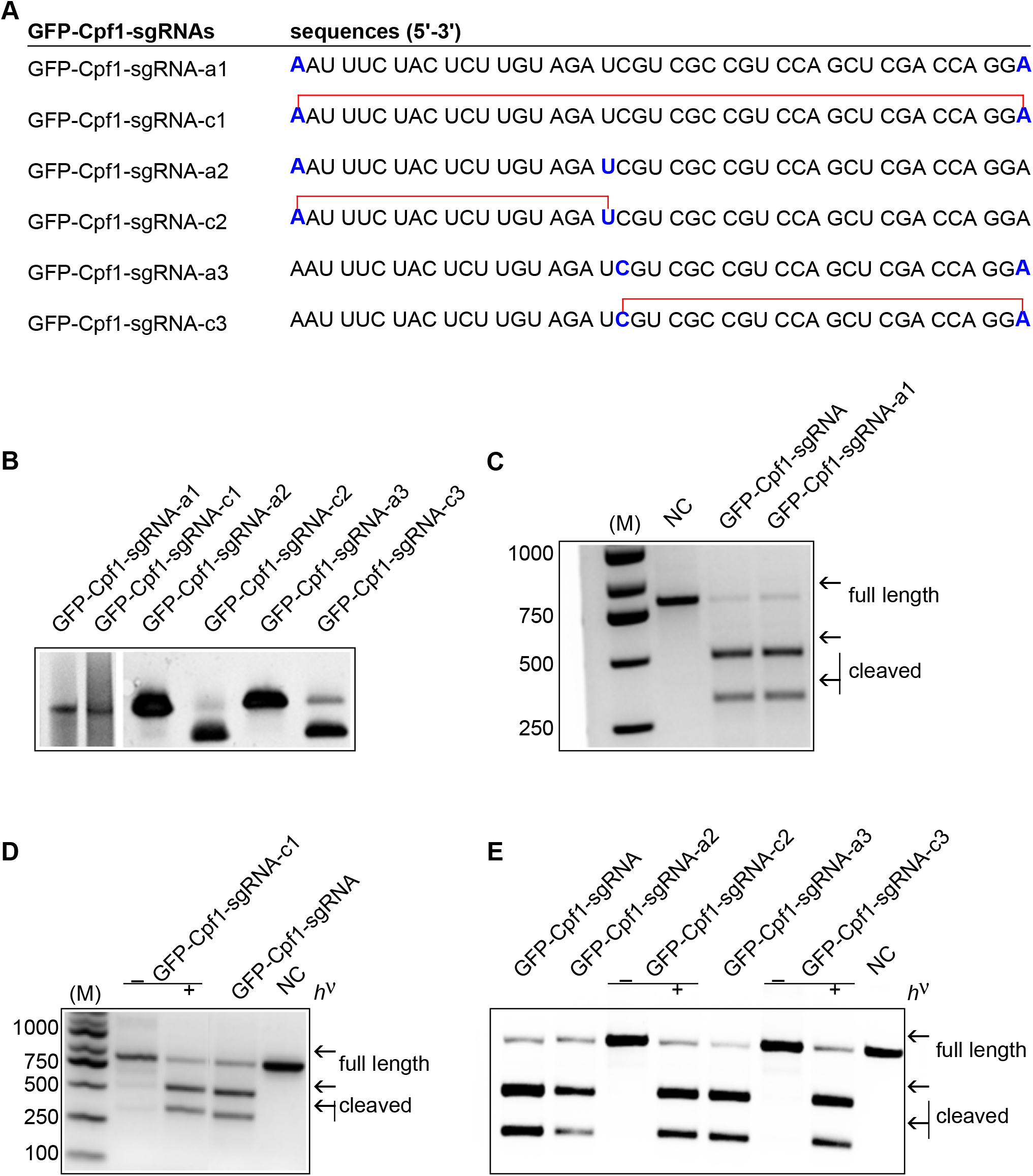
Design, synthesis and evaluation of long sgRNAs and its circular forms and their application in CRISPR/Cpf1 editing. **A**. Sequences of synthetic acyclic NPBOM modified and cyclized sgRNAs targeting GFP gene. Blue bold letters indicate the modification sites substituted with NPBOM, and the red lines indicate the NPBOM derived linkage upon reacting with 1,4-bis(azidomethyl)benzene. **B**. Polyacrylamide gel electrophoresis analysis of the linear GFP-Cpf1-sgRNA-a and circular GFP-Cpf1-sgRNA-c. **C**. Representative gel images of cleavage detection assay of the target GFP gene *in vitro* with native GFP-Cpf1-sgRNA and acyclic GFP-Cpf1-sgRNA-a1. **D**. Representative gel images of cleavage detection assay of the target GFP gene *in vitro* with native GFP-Cpf1-sgRNA and circular GFP-Cpf1-sgRNA-c1 with and without light irradiation (5 minutes, 20 mW·cm^−2^). **E**. Representative gel images of cleavage detection assay of the target GFP gene *in vitro* with native GFP-Cpf1-sgRNA, acyclic GFP-Cpf1-sgRNA-a2/a3, and circular GFP-Cpf1-sgRNA-c2/c3 with and without light irradiation (5 minutes, 20 mW·cm^−2^).

Specifically, a 60 seconds’ irradiation evidently activated the Cpf1 function by fully releasing the parent linear GFP-Cpf1-sgRNA (Fig. 3D). In order to distinguish whether the inactivation of CRISPR/Cpf1 system with circular sgRNA was due to the incomplete recognition of the complementary DNA sequence or recruitment failure of endonuclease, we synthesized two distinct pairs of sgRNAs in which the repeat handler before the 5’-TTTN-3’ PAM (GFP-Cpf1-sgRNA-c2) and the seed segment after PAM (GFP-Cpf1-sgRNA-c3) were annulated, respectively (Fig. 3A, B, Fig. S27-30). Photo-responsive cytidine and uridine building blocks containing NPBOM were used for the covalent cyclization (Fig. S3-4). The photo-modulation of these partly cyclized sgRNAs were explored to elucidate the detailed interaction between the sgRNA with target DNA and Cpf1 protein (Fig. 3E). Interestingly, those two smaller circular sgRNAs also displayed no activities before light irradiation, indicating that the Cpf1 interaction (blocked in GFP-Cpf1-sgRNA-c2) and protospacer pairing with target DNA sequence (blocked in GFP-Cpf1-sgRNA-c3) were both crucial for efficient genome editing in CRISPR/Cpf1 system. Not surprising, the 365 nm light irradiation towards circular sgRNAs fully recovered their Cpf1 cleavage (Fig. 3D, E). This study further suggested that our light-activable circular gRNA strategy is a generic approach for regulating RNA-guided CRISPR editing.

### Photomodulation of CRISPR/Cas9 editing with circular crRNA in live cells

The above results encouraged us to explore the opportunity of this circular photo-responsive crRNA strategy for controllable gene editing in live cells. We rationalized that once the circular crRNA was transfected in cells with tracrRNA and Cas9 mRNA, it would stay inert until expose to the UV light which result in fragmentation of the crRNA and then initiate further CRISPR editing. As a proof-of-concept, we first tested this approach in HEK293T-GFP cells that stably expressed GFP transgene copies (Fig. 4A, Fig. S14). The previously validated crRNA sequence targeting the GFP gene was used to quantify the knockout (KO) efficiency with cyclic gRNAs. The efficiency or GFP KO by GFP-crRNA-c2/tracrRNA/Cas9 mRNA co-transfection at different amount was determined using flow cytometry. The system is not detectably leaky in the dark over a wide range of RNA concentrations (5.4 % *vs*. 9.4 % of indel in GFP gene before and after transfection at 4 pmol/well). However, a maximum level of 56.6 % of the cells had GFP knocked out upon light irradiation (relative to native GFP-crRNA). Therefore, the transfection of chemically synthesized crRNAs with light-sensitive functionalities is a highly effective method for target gene KO.

**Fig. 4.**
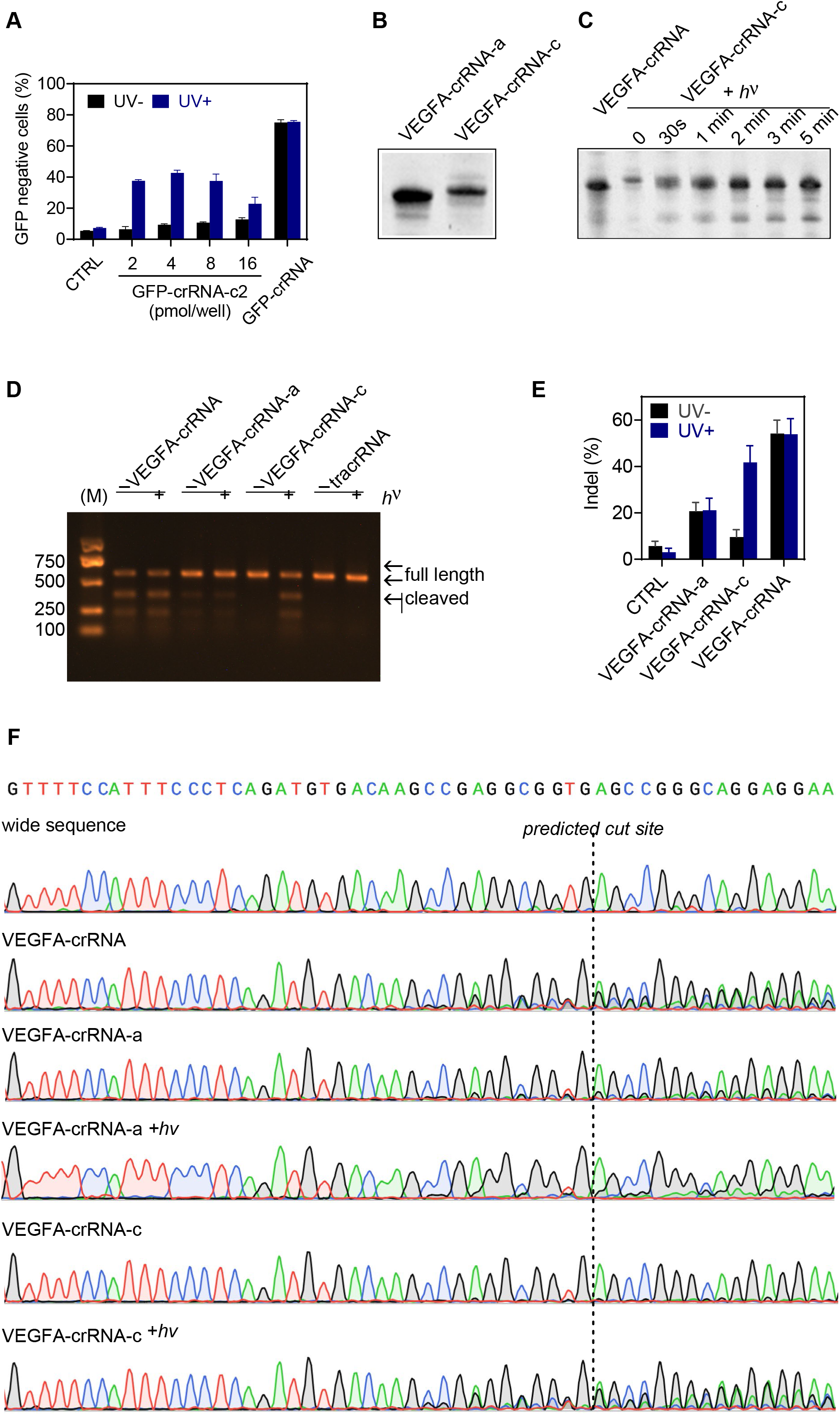
Light-induced CRISPR-Cas9-mediated gene editing in live cells. **A**. Quantification of GFP negative cells deletion by native GFP-crRNA and photocaged GFP-crRNAs at viable concentrations with 365 nm light illumination (5 minutes, 10 mW·cm^−2^). CTRL = nontreated cells. Results are measured *via* flow cytometry and expressed as individual data points overlaid with the mean ± SEM (*N* = 3). **B**. Polyacrylamide gel electrophoresis analysis of the linear VEGFA-crRNA-a and circular VEGFA-crRNA-c. **C**. Comparison of native VEGFA-crRNA and light-mediated uncaging reaction of VEGFA-crRNA-c monitored by gel images at various time points. **D**. Representative gel images of cleavage detection assay of the target VEGFA gene in HEK293T cells with native VEGFA-crRNA, acyclic VEGFA-crRNA-a, circular VEGFA-crRNA-c with and without light irradiation (5 minutes, 10 mW·cm^−2^). tracrRNA alone was used for comparison. **E**. Quantification of target VEGFA gene mutation rates from gel images using ImageJ software. Results are expressed as individual data points overlaid with the mean ± SEM (*N* = 3). **F**. Representative Sanger sequencing chromatograms of mutations in CRISPR/Cas9-treated HEK293T cells. Indels from indicated sample were PCR amplificated and Sanger sequenced. From top to bottom, wild sequence, VEGFA-crRNA, VEGFA-crRNA-a, VEGFA-crRNA-a with light irradiation, VEGFA-crRNA-c and VEGFA-crRNA-c with light irradiation (both in 5 minutes, 10 mW·cm^−2^). The dot line indicated the beginning of predicted cut sites.

Vascular endothelial growth factor A is a potent and specific mitogen for endothelial cells, which is secreted in response to hypoxic and inflammatory stimuli. We intend to employ this photo-responsive CRISPR-Cas9 editing strategy to disrupt the VEGFA gene in human 293T cells. A 36 nt VEGFA-crRNA was selected to target locations in the intron sequence, from which a frameshift mutation would occur and results in truncated and/or nonfunctional gene products (Table S1). The corresponding NPBOM decorated VEGFA-crRNA-a and the circular product VEGFA-crRNA-c were prepared similarly (Fig. 4B), the latter also exhibited a similar light dissociation pattern, releasing the linear VEGFA-crRNA in less than 1 minute upon UV irradiation (Fig. 4C). To validate the possibility of photo-responsive disruption of VEGFA gene in human cells, we later infected 293T-Cas9 cells constitutively expressing Cas9 with native crRNA to target VEGFA. The indel frequency was around 5.7% by using the T7 endonuclease I (T7E1) mismatch detection assay (Fig. 4D, E). The indel formation slightly dropped with light irradiation (3.0%). In consistence to the *in vitro* data, a considerable deletion was observed when the linear VEGFA-crRNA-a was used. Further light treatment did not affect the indel efficiency (21.1% *vs*. 20.7%). This result is explained by the fact that bis-NPBOM decorated crRNA was subsequently degraded by exonucleases upon delivery. Interestingly, circular crRNA-c with a covalently linked ring showed a 9.6% indel efficiency in the dark, and led to a comparable indel efficiency (41.8%) upon 5 minutes’ exposure to UV light. This result was substantially identical from experiments conducted with the native crRNA (53.9%). Sanger sequencing also confirmed that indel formation occurred at the predicted cutting sites, including a mixture of frameshift and in-frame mutations (Fig. 4F). Taking together, all these results provide as proof of concept of the possibility of implementing a photo-responsive gene editing technique with circular gRNAs.

### Photomodulation of CRISPR/Cas9 editing with bow-knot-type gRNA in mouse embryos

Gene editing with CRISPR/Cas9 has empowered the induction of site-specific mutations in mammalian embryos.^66^ Nonetheless, mosaic patterns have always been observed in the genotypes of mutant embryos from zygotes treated with CRISPR/Cas9 editing, the reason of which was predicted to appear due to the timing difference between the gene editing and the first replication of the zygotic genome. A temporal CRISPR/Cas9 system that is functional prior to the first genome replication is highly needed for generating non-mosaic mutants. We hypothesized that if the circular crRNA and other components could be injected into the developing zygotes, followed by light irradiation at specific timing, the problem of controlling editing before replication may be partly solved. To the best of our knowledge, such spatial and temporal control of gene editing with CRISPR/Cas9 system in zygotes has not been realized so far.

In the first stage of testing the above hypothesis, we constructed a circular crRNA targeting the myostatin (Fig. 5A). The 100 nt *MSTN*-crRNA-c1 was smoothly prepared by standard click condensation between the two terminal NPBOM substituted adenine and uridine (Fig. S12). As far as we can tell, this is one of the largest ring-structured nucleic acids, which was still able to be converted to the linear *MSTN*-crRNA in less than 1 minute using UV light (Fig. 5B). However, unsuccessful disruption of the MSTN was observed with the circular crRNA or its linear precursor *MSTN*-crRNA-a1 with Cas9 and tracrRNA (Fig. 5C). The reason for the incomplete halting of editing is the fact that the circular RNA longer than 100 bases may adopt very flexible confirmations in solutions, some of which include flat surface that may partially contribute to the recognition of target DNA (Fig. 5A). This preliminary result prompted us to design a more complex crRNA by placing an additional NPBOM substituent in the middle of the sequence and coupling with a three-arm structure 1,3,5-tris(azidomethyl)benzene (Fig. 5A, Fig. S5, S12). To our delight, the new bow-tie architecture was obtained with enough rigidity to reduce any possible binding upon interaction with the target DNA or Cas 9 protein, which was also confirmed by the thorough halting of CRISPR-Cas-mediated DNA cleavage. In the dark, practically no gene editing could be detected (Fig. 5C). On the other hand, the twist knot, with higher tensions, was readily untied by light irradiation within only 1 minute, which afforded a narrow temporal window for the editing activation (Fig. S13). A substantial restoration of the native sgRNA’s function was achieved (Fig. 5C), again proved the feasibleness of photocaging gRNAs and light-responsive DNA cleavage upon swift exposure to light irradiation *in vitro*.

**Fig. 5.**
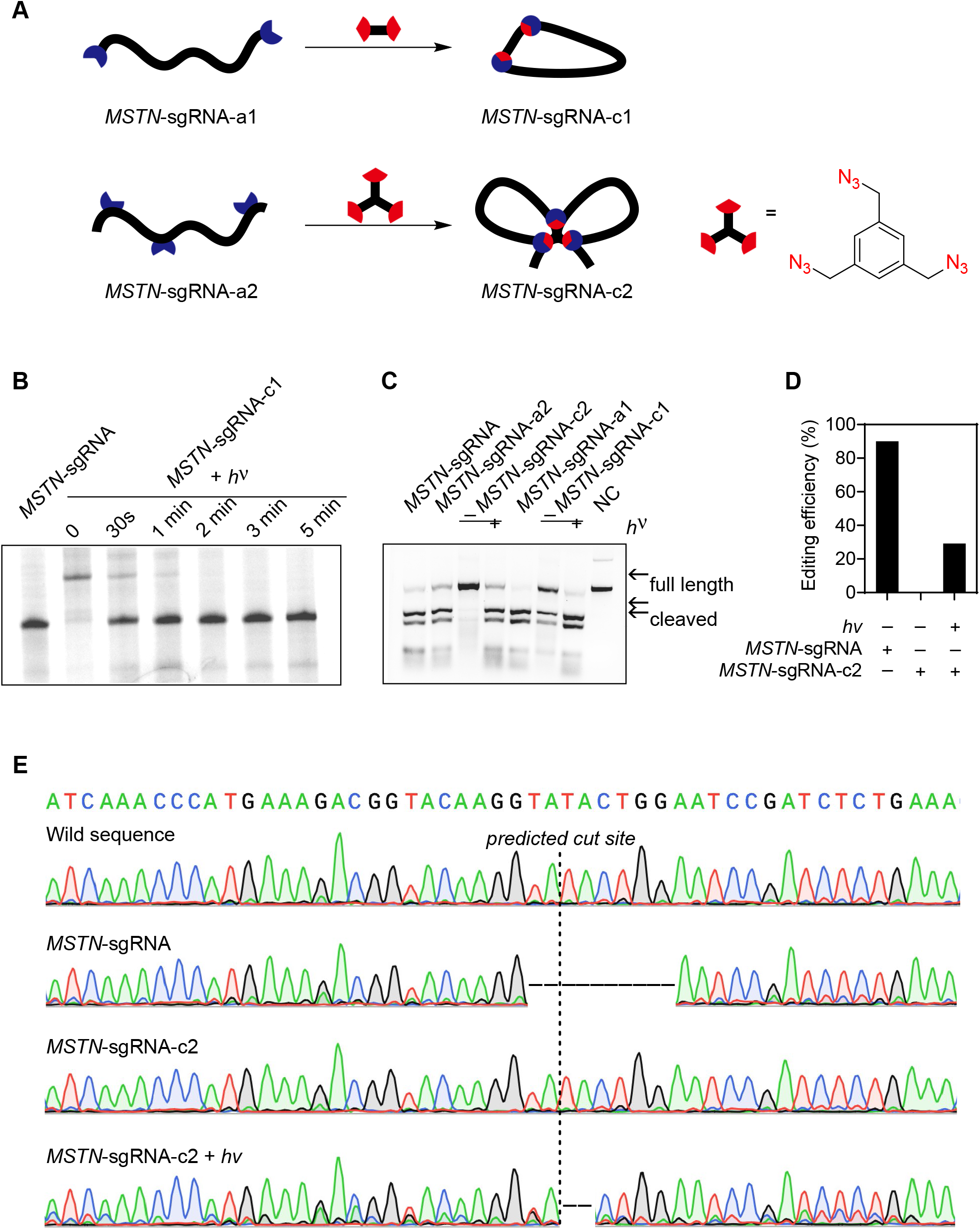
Light-induced CRISPR-Cas9-mediated *MSTN* gene editing in mouse zygotes. **A**. Design and synthesis of two types of circular *MSTN*-sgRNAs. **B**. Comparison of native *MSTN*-sgRNA and light-mediated uncaging reaction of *MSTN*-sgRNA-c1 monitored by gel images at various time points. **C**. Biochemical activity assay of target *MSTN* DNA cleavage with native *MSTN*-sgRNA, acyclic *MSTN*-sgRNA-a1/a2 and circular *MSTN*-sgRNA-c1/c2 with and without light irradiation (5 minutes, 20 mW·cm^−2^). NC = Negative control. **D**. Quantification of mutated zygote percentages after microinjection of *MSTN*-sgRNAs. **E**. Representative Sanger sequencing chromatograms of mutations in CRISPR/Cas9-treated zygotes. Indels from indicated sample were PCR amplificated and Sanger sequenced. From top to bottom, wild sequence, *MSTN*-sgRNA, *MSTN*-sgRNA-c and *MSTN*-sgRNA-c with light irradiation (90 seconds, 10 mW·cm^−2^). The dot line indicated the beginning of predicted cut sites. Horizontal line indicates the missing nucleotides cites.

Finally, we applied this system to spatiotemporally control gene editing in mouse zygotes. The zygotes were injected with MSNT-sgRNA or MSTN-sgRNA-c2 and further cultured to cleavage and blastocyst stages. Sanger sequencing of the sampled embryos revealed high efficiency (90%, 9/10) of mutations at the target sites in MSNT-sgRNA injection groups, confirming the efficacy of the designed sgRNA (Fig. 5D, 5E). Notably, no editing on *MSTN* were found in embryos injected with *MSTN*-crRNA-c2 in the absence of light irradiation, affording an excellent protection from gene editing with this bow-tie sgRNA even in the complex intracellular conditions. However, when zygotes injected with photocaged *MSTN*-crRNA-c2 were exposed to a 90 seconds’ of UV light, around 30% (7/24) of the gene-editing efficiency was reinstated (Fig. 5D), enabling a highly potential for gene editing at the designated locations and/or preferred times. A higher efficiency may be achieved if prolonger irradiation time or optimized procedure are employed on the premise that the embryo is not damaged. The variety of the mutations close to the predicted cleavage site on the *MSTN* gene was also analyzed by sequencing the PCR fragments amplified from the mutated zygotes (Fig. 5E). Missing nucleotides were commonly detected as the dominant mutations (Fig. S18).

## Conclusion

The CRISPR/Cas-based gene editing technology is a revolutionary breakthrough in genetic engineering, which since its discovery has offered a fascinating platform to assessing gene function and precisely manipulating cellular behavior and function due to the simple design of this system and its powerful capacity in editing different loci simultaneously. Nonetheless, the inability of conducting precise gene editing in certain cells/tissues/organisms at a specific time point may lead to adverse consequences like off-target effects and mosaic mutations and therefore limits the clinical translation of this technology. Strategies using genetic regulation and chemical and biophysical approaches to regulate the activity of CRISPR/Cas systems have recently emerged and present promising results in the improvement of spatial and/or temporal controllability. An inspiriting method was to employ a light-activatable gRNA to block its recognition ability of hybridizating with complementary DNA sequences or impede its recruitment of Cas proteins. In most cases, a relatively fast response to the UV light irradiation was achieved. However, these systems usually suffer from substantial leakage of activity in the dark, possibly due to partially functionalization of the modified gRNA or its dissociation products cleaved by intracellular nuclease. Another disadvantage is the relatively high UV dose or irradiation time required to degrade the photoresponsive protectors and to restore gene editing. Herein, in this article, we envisioned that a simpler means by constructing covalently linked circular gRNAs through a head-to-tail biorthogonal condensation. Comparing with its linear counterpart gRNA with multiple photolabile groups, the new circular gRNA is lack of the flexibility. The several conformational changes not only disrupt the base-pairing interactions with the target DNA sequence to avoid any partially access, but also prohibit the inducing of active conformation of Cas for target DNA editing. Upon the light irradiation, the circular gRNA would rapidly release its conformational strain and the molecule resembles more its linear analogue, restoring the CRISPR/Cas editing capacity. Therefore, the photoactivation control with circular gRNA would afford a rapid and non-invasive activation of CRISPR/Cas activity in a generic manner towards any specified sequences with minimal background editing. We have demonstrated this advanced strategy in the cleavage of GFP and VEGFA locus *in vitro* and in live cells. Another significant advantage of this approach is its capacity to incorporate minimal nucleotide sites with photoliable functionalities in longer gRNAs by geometrically compressing into complex architectures, such as bow-tie styles. The above method combine external light stimuli has realized the spatiotemporal control of CRISPR/Cas-based gene editing of *MSTN* in zygotes, which established that this biorthogonally activatable circular guide RNA was applicable to control the CRISPR systems. We predict that these features will be used to investigate spatiotemporally complex physiological events since it will help to better understand the dynamics of gene regulation.

## Methods

Methods and additional data are available in the Supplementary Information.

### Circularization of NPBOM incorporated oligonucleotides

Circular gRNAs were prepared *via* copper-catalyzed azide alkyne cycloaddition reactions (CuAAC) with NPBOM incorporated oligonucleotides and different azido-linkers. Generally, to an aqueous solution of linear modified gRNAs (1.5 nmol) in a reaction tube was added the following reagents: triethylammonium acetate buffer, bis-azido-linker (1.5 nmol), magnesium chloride (0.1 μmol), fresh ascorbic acid (2.5 μmol), DMSO and copper sulfate-tris(3-hydroxypropyltriazolylmethyl)amine (2.5 μmol). Then prepared circular oligonucleotides were purified by HPLC and analyzed by denatured PAGE gel and MALDI-TOF.

### Live-cell editing of VEGFA gene with photo-responsive gRNAs

293T-Cas9 cells stably expressing Cas9 protein (UBIGENE) were cultured in DMEM with 10% FBS overnight. Pre-transfection experiments were performed using different doses of unmodified crRNA/tracrRNA duplexes and Lipofectamine 3000 in Opti-MEM (Gibco) for 15 minutes at room temperature before mixing with 293T-Cas9 cells. The medium was changed to complete DMEM after an incubation of 6 hours and the cells were further cultured another 72 hours at 37 °C. Genomic DNAs were extracted using TIANamp Genomic DNA Kit for polymerase chain reaction (Vazyme) to amplify the target DNA fragments. The T7E1 assays of the PCR products were performed according to manufacturer’s protocol. Products were analyzed by 2 % agarose gel electrophoresis. For light-activated gene editing, 293T-Cas9 cells were transfected with crRNA/tracrRNA duplexes as described above. After the transfection mixture was replaced with fresh cell culture medium, the cells were subjected to 5 minutes of 365 nm irradiation by a UV lamp (365 nm, 10 mW·cm^−2^). The PCR amplicons were also sequenced by Sanger sequencing.

### Light-activated MSTN gene editing in zygotes

All the studies with mouse zygotes were conducted in accordance with the Guide for the Care and Use of Laboratory Animals (Ministry of Science and Technology of China, 2006), and were approved by the animal ethics committee of China Agricultural University. The female mice (6-week old ICR mice from Beijing Vital River Experimental Animals Centre) were injected intraperitoneally with 10 IU of pregnant mare serum gonadotropin (PMSG) (Ningbo Sansheng Biological Technology, Ningbo, China), followed by injection with 10 IU of human chorionic gonadotropin (hCG) (Ningbo Sansheng Biological Technology) 46-48 hours later. The female mice were immediately caged with mature male mice and zygotes were harvested from oviducts ∽20 hours after hCG injection. After several washes, the zygotes were placed into drops of M2 medium. The zygotes were microinjected with ∽10 pL of the RNPs solution (Cas9 protein and different sgRNAs in TE buffer) using a microinjection pump (Eppendorf Femtojet) under an inverted fluorescence microscope (Nikon TE300, Japan). The *hv*+ group was followed by irradiation with ultraviolet light (excitation wavelengths of 365 nm, 10 mW·cm^−2^) for 90 seconds and others were incubated in dark. After the injections were completed, the zygotes were transferred to 50 μL-drop of KSOM medium and cultured to blastocyst stage, in humidified 5% CO_2_ at 37 °C. The embryos were harvested and placed into PCR reaction solution, and the target genes were amplified for sequencing analysis. Two rounds of continuous PCR were used for amplification.

## Acknowledgments

This work was supported by the National Key R&D Program of China (2017YFA0208100 and 2020YFA0707901), National Natural Science Foundation of China (22022704 and 21977097) and Chinese Academy of Sciences. We thank Prof. Yuan-Chen Dong at CAS Laboratory of Colloid, Interface and Chemical Thermodynamics, ICCAS for assistance of MALDI-TOF measurement of synthetic gRNAs. We also thank Prof. Wei Li, Prof. Hao-Yi Wang at Institute of Zoology, Chinese Academy of Sciences and Prof. Liang Xu at Sun Yat-sen University for insightful discussions.

## Author contributions

Y.J.S, J.L, J.J.L, Y.Z, W.D.C. and W.Q.C. performed the experiments. Y.J.S, X.J.T, J.H, M.W. and L.C. designed the experiments and analyzed the data. M.W. and L.C. conceived the experiments. All authors discussed the results and contributed to the preparation and editing of the manuscript.

## Notes

Institute of Chemistry, Chinese Academy of Sciences (ICCAS) has filed a patent on the discovery and development of this method that are described in the paper at the National Intellectual Property Administration.

