## Supplementary material for "Optical control of CRISPR-Cas editing with cyclically caged guide RNAs": SI

### Optical control of CRISPR-Cas editing with cyclic caged gRNAs

*Beijing National Laboratory for Molecular Sciences (BNLMS), CAS Key Laboratory of Molecular Recognition and Function, CAS Research/Education Center for Excellence in Molecular Sciences, Institute of Chemistry, Chinese Academy of Sciences, Beijing 100190, China; BNLMS, CAS Key Laboratory of Analytical Chemistry for Living Biosystems, Institute of Chemistry, Chinese Academy of Sciences, Beijing 100190, China; State Key Laboratory of Agrobiotechnology and College of Biological Science, China Agricultural University, Beijing, 100193, China; State Key Laboratory of Natural and Biomimetic Drugs, School of Pharmaceutical Sciences, Peking University, Beijing 100191, China; Hangzhou Institute for Advanced Study, University of Chinese Academy of Sciences, Chinese Academy of Sciences, Hangzhou 310024, China and University of Chinese Academy of Sciences, Beijing 100049, China.*

 (L.C.)

**Table of contents**


| | | |
|---|---|---|
| 1 | General information | S3 |
| 2 | Chemical synthesis | S3 |
| 3 | Preparation of oligonucleotides | S10 |
| 4 | General procedure for oligonucleotide circularization | S10 |
| 5 | Decaging assay of circular crRNAs | S11 |
| 6 | CrRNA stability assay (RNase A) | S11 |
| 7 | Expression and purification of Cas9/Cpf1 protein | S12 |
| 8 | *In vitro* DNA cleavage assay | S12 |
| 9 | Binding assay of circular gRNA | S13 |
| 10 | Light-activated exogenous GFP gene editing in cells | S13 |
| 11 | Light-activated endogenous VEGFA gene editing in cells | S13 |

12 Light-activated *MSTN* gene editing in zygotes S14

13 Supplementary pictures S15

14 NMR and mass spectra of new compounds S37

15 References S49

#### 1. General information

All chemical reagents were purchased from commercial suppliers and were used without further purification. The $^{1}H$ NMR, $^{13}C$ NMR and $^{31}P$ NMR spectra were recorded on Bruker AVANCE 300 MHz, AVANCE 400 MHz or AVANCE III HD 500 MHz spectrometers. Chemical shifts δ are given in ppm relative to the residual proton signals of the deuterated solvent for $^{1}H$ and $^{13}C$ NMR. Mass spectra were collected by Thermo Fisher Scientific (Exactive). HPLC analysis was performed on an Agilent 1260 system using the PLRP-S column (250 mm × 4.6 mm, 100 Å, 8 μm). crRNAs were purchased from Biosyntech (Suzhou). Alt-R® tracrRNA, Alt-R® Genome Editing Detection Kit (T7E1) and dCas9 protein were purchased from Integrated DNA Technologies (IDT). DPBS buffer, DMEM, Opti-MEM and Penicillin/Streptomycin were purchased from Gibco. FBS was purchased from Biological Industries (BI). Lipofectamine 3000 was purchased from Thermo Fisher Scientific. NEB buffer 3.1 and Gel loading Dye, Blue (6x) were purchased from New England Biolabs (NEB). 40% Page pre-solution, 10 x TBE, 50 x TAE and the nucleic acid stains 10000 x Solar Red were purchased from Solarbio. Fastpure Gel DNA Extraction mini Kit and 2 × Taq Master Mix (Dye Plus) were purchased from Vazyme. The concentration of RNA was quantified by Nanodrop One (Thermo Fisher Scientific). Gel Imaging was performed with a Tanon-5200 Multi Fluorescence Imager.

#### 2. Chemical synthesis

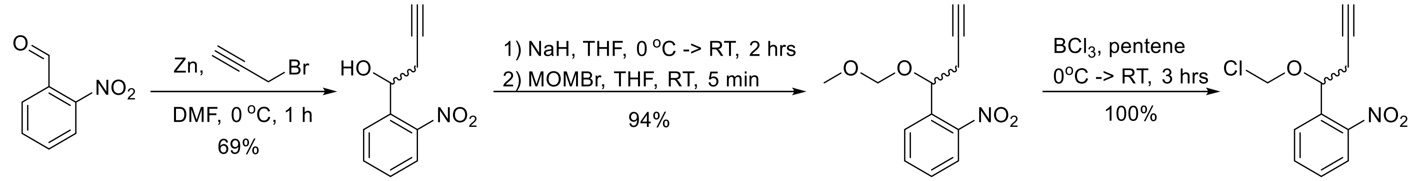


1 2 3 4

**Figure S1.** Synthesis of photo-responsive alkyne linkage **4**

To a suspension of activated zinc powder (8.3 g, 127 mmol) in dry dimethylformamide (100 mL) was added propargyl bromide (8.1 mL, 75.8 mmol) dropwise at 0 °C. The mixture was stirred at 0 °C for 1 hour and then 2-nitrobenzaldehyde **1** (7.6 g, 50 mmol) was added. The reaction was stirred for another 15 minutes and was then quenched by adding saturated ammonium chloride solution. The mixture was extracted three times with ethyl acetate. The combined organic layers were dried over sodium sulfate, filtrated and then evaporated to dryness under reduced pressure. The residue was purified by column chromatography (petroleum ether/ethyl acetate 5/1) to afford the intermediate **2** as a brown oil (6.6 g, yield 69%). $^{1}H$ NMR (300 MHz, $CDCl_3$): δ 7.97-7.94 (d, *J* = 9.0 Hz, 1H), 7.90-7.87 (d, *J* = 8.1 Hz, 1H), 7.70-7.65 (t, *J* = 7.4 Hz, 1H), 7.49-7.43 (d, *J* = 7.7 Hz, 1H), 5.47-5.45 (m, 1H), 2.95-2.87 (m, 2H), 2.71-2.63 (m, 1H), 2.12-

2.10 (t, $J$ = 2.6 Hz, 1H). $^{13}$C NMR (75 MHz, DMSO-$d_6$): δ 147.7, 138.7, 133.1, 128.6, 128.5, 123.7, 81.0, 72.9, 66.3, 28.1.[1]

To a stirring suspension of sodium hydride (0.8 g, 20 mmol, 60% dispersion in mineral oil) in dry tetrahydrofuran (90 mL) under nitrogen was added the solution of **2** (1.9 g, 10 mmol) in tetrahydrofuran (10 mL) dropwise at 0 °C. The mixture was stirred for another 15 minutes and then another 2 hours at room temperature. Bromomethyl methyl ether (0.98 mL, 12 mmol) was then added dropwise. After 5 minutes, the mixture was filtered through celite and the solvent was evaporated to dryness. The residue was purified by column chromatography (petroleum ether/ethyl acetate 15/1) to afford the intermediate **3** as a brown oil (2.2 g, yield 94%). $^1$H NMR (500 MHz, $CDCl_3$): δ 7.97-7.96 (d, $J$ = 8.2 Hz, 1H), 7.86-7.84 (d, $J$ = 7.9 Hz, 1H), 7.68-7.65 (t, $J$ = 7.6 Hz, 1H), 7.48-7.45 (t, $J$ = 7.8 Hz, 1H), 5.44-5.42 (m, 1H), 4.68-4.67 (dd, $J$ = 2.9 Hz, 6.9 Hz, 1H), 4.55-4.53 (dd, $J$ = 3.2 Hz, 6.9 Hz, 1H), 3.37 (s, 1H), 2.89-2.85 (m, 1H), 2.78-2.73 (m, 1H), 2.034-2.028 (d, $J$ = 2.9 Hz, 1H). $^{13}$C NMR (125 MHz, $CDCl_3$): δ 148.3, 136.4, 133.3, 128.95, 128.67, 124.5, 95.3, 80.0, 71.4, 70.7, 56.0, 27.3. HRMS (ESI): $m/z$ $[M + H]^+$ calcd for $C_{12}H_{14}NO_4$: 236.0917; found: 236.0917.

To a suspension of intermediate **3** (540 mg, 2.3 mmol) dry pentane (15 mL) was added a solution of boron chloride (1 M, 1.1 mL, in hexane) dropwise. The mixture was stirred at 0 °C for 5 minutes, and was then warmed to room temperature and allowed to stir for another 2 hours. When the reaction was completed as indicated by $^1$H NMR analysis, the mixture was then concentrated under reduced pressure, and the crude product **4** was used in the next step without purification. $^1$H NMR (500 MHz, $CDCl_3$): δ 8.02-8.01 (d, $J$ = 8.2 Hz, 1H), 7.79-7.77 (d, $J$ = 7.9 Hz, 1H), 7.70-7.67 (t, $J$ = 7.4 Hz, 1H), 7.52-7.49 (t, $J$ = 7.4 Hz, 1H), 5.61-5.59 (m, 2H), 5.33-5.32 (d, $J$ = 6.1 Hz, 1H), 2.93-2.78 (m, 2H), 2.05-2.04 (t, $J$ = 2.5 Hz, 1H). $^{13}$C NMR (125 MHz, $CDCl_3$): δ 148.2, 134.7, 133.5, 129.2, 129.0, 124.7, 80.5, 78.9, 74.2, 71.4, 26.9.

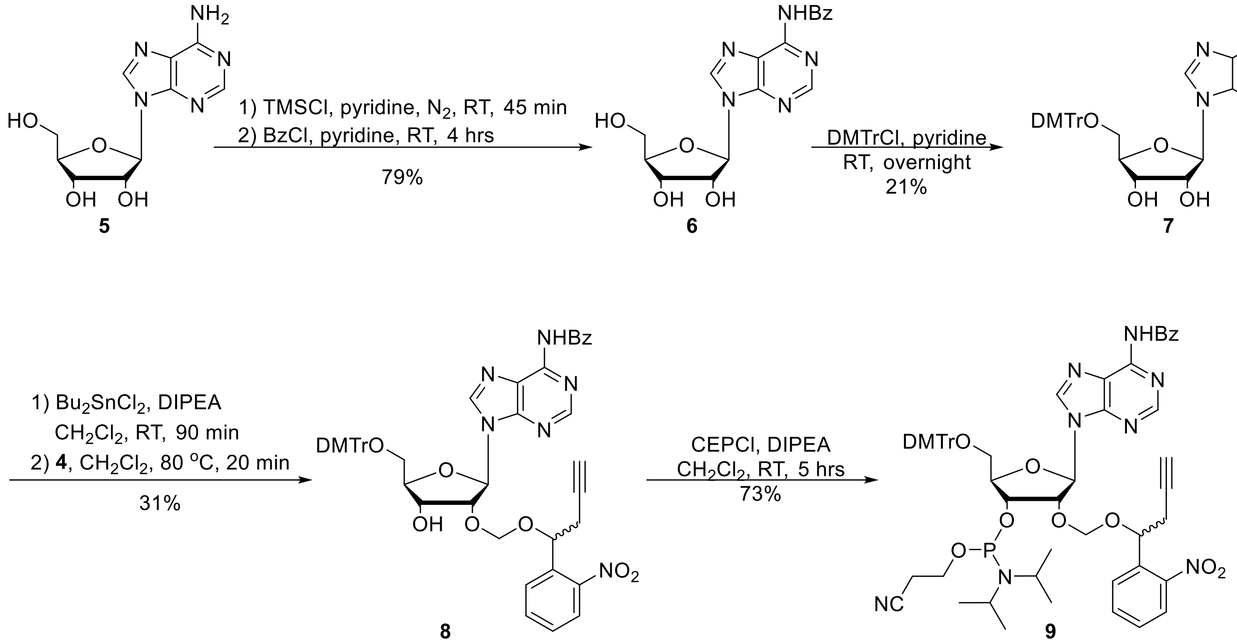


**Figure S2**. Synthesis of photo-responsive adenosine phosphoramidite

To a solution of adenosine **5** (10.7 g, 40 mmol) dry pyridine (100 mL) was added trimethylsilyl chloride (38 mL, 300 mmol) dropwise. The mixture was stirred at room temperature for 45 minutes and was then added benzoyl chloride (23 mL, 200 mmol). The reaction was allowed to stir for another 4 hours and then quenched with water at 0 °C. After 20 minutes the reaction was quenched with ammonium hydroxide (25%, 20 mL) at room temperature, stirred for 45 minutes and then concentrated under reduced pressure. The residue was recrystallized with methanol and water. **6** precipitated as a white solid and was collected by filtration and dried under air (11.7 g, yield 79%), whose $^{1}$H NMR and $^{13}$C NMR spectra were identical to that reported in literature.[2]

To a solution of **6** (5.6 g, 15.1 mmol) in dry pyridine (100 mL) was added 4,4'-dimethoxytrityl chloride (7.7 g, 22.7 mmol) under nitrogen. The mixture was stirred overnight at room temperature, evaporated to dryness and purified by column chromatography (dichloromethane/methanol 50/1-30/1). The crude intermediate **7** was obtained as a white foam (2.1 g, yield 21%), whose $^{1}$H NMR and $^{13}$C NMR spectra were identical to that reported in literature.[3]

To a solution of intermediate **7** (1.0 g, 1.5 mmol) in dry dichloromethane (25 mL) was added *N*,*N*-diisopropylethylamine (1.2 mL, 7.5 mmol) and dibutyltin dichloride (559 mg, 1.84 mmol), respectively. The mixture was stirred at room temperature for 2 hours and then a solution of **4** (crude product, 2.3 mmol) in dichloromethane (5 mL) was added dropwise. The reaction mixture was heated and stirred at 80 °C for 20 minutes, until TLC indicated the consumption of **7**.

The reaction was quenched with aqueous sodium carbonate. The aqueous layer was extracted three times with dichloromethane. The organic phases were combined, dried over sodium sulfate and filtrated. The filtrate was evaporated to dryness and purified by column chromatography (dichloromethane/ethyl acetate 3/1) to afford the product **8** as a yellow foam (800 mg, yield 31%). $^{1}H$ NMR (300 MHz, $CDCl_3$): δ 9.05-9.03 (m, 1H), 8.74-8.67 (m, 1H), 8.23-8.11 (m, 1H), 8.04-7.90 (m, 3H), 7.72-7.19 (m, 14H), 6.82-6.79 (m, 4H), 6.28-6.11 (m, 1H), 5.46-5.29 (m, 1H), 5.06-4.91 (m, 2H), 4.79-4.62 (m, 1H), 4.40-4.35 (m, 1H), 4.26-4.19 (m, 1H), 3.78 (s, 6H), 3.51-3.36 (m, 2H), 2.83-2.55 (m, 3H). $^{13}C$ NMR (125 MHz, DMSO-$d_6$): δ 158.5, 152.3, 152.2, 151.0, 148.5, 145.3, 143.9, 136.0, 135.9, 135.3, 133.8, 132.9, 130.1, 129.6, 129.3, 129.0, 128.9, 128.2, 128.1, 127.1, 124.6, 113.6, 85.9, 80.1, 73.4, 71.8, 29.5, 27.0, 26.7, 22.6, 14.4. HRMS (ESI): *m/z* $[M + H]^+$ calcd for $C_{49}H_{45}N_6O_{10}$: 877.3192; found: 877.3200.

To a solution of intermediate **8** (450 mg, 0.5 mmol) in dry dichloromethane (10 mL) under nitrogen was added *N*,*N*-diisopropylethylamine (DIPEA, 424 μL, 2.6 mmol). The mixture was stirred for 5 minutes, followed by the addition of 2-cyanoethoxy-*N*,*N*-diisopropylaminochlorophosphine (CEPCl, 228 μL, 1.0 mmol). The reaction was quenched after 2 hours by adding aqueous sodium bicarbonate (15 mL). The aqueous layer was extracted three times with dichloromethane. The organic phases were combined, dried over sodium sulfate and filtrated. The filtrate was evaporated to dryness under reduced pressure and then purified by column chromatography (dichloromethane/methanol 150/1, with 0.5% triethylamine) to afford the product **9** as yellow foam (400 mg, yield 73%). $^{1}H$ NMR (400 MHz, $CDCl_3$): δ 9.04-8.98 (m, 1H), 8.77-8.62 (m, 1H), 8.30-7.83 (m, 5H), 7.61-7.26 (m, 12H), 6.82-6.79 (m, 5H), 6.28-5.89 (m, 1H), 5.36-5.20 (m, 2H), 4.94-4.56 (m, 3H), 4.38-4.31 (m, 1H), 3.78 (s, 6H), 3.67-3.32 (m, 6H), 2.70-2.54 (m, 5H), 1.32-1.01 (m, 12H). $^{13}C$ NMR (125 MHz, $CDCl_3$): δ 158.9, 153.2, 151.8, 136.5, 135.8, 134.0, 133.1, 130.2, 130.1, 129.0, 128.9, 128.28, 128.25, 128.2, 127.9, 127.8, 113.2, 92.2, 91.5, 71.7, 71.2, 43.4, 42.0, 29.8, 29.7, 29.6, 29.5, 29.3, 29.2, 27.2, 27.1, 27.0, 24.62, 24.58, 24.5. $^{31}P$ NMR: δ 150.66, 150.51. HRMS (ESI): *m/z* $[M + H]^+$ calcd for $C_{58}H_{62}N_8O_{11}P$: 1077.4270; found: 1077.4270.

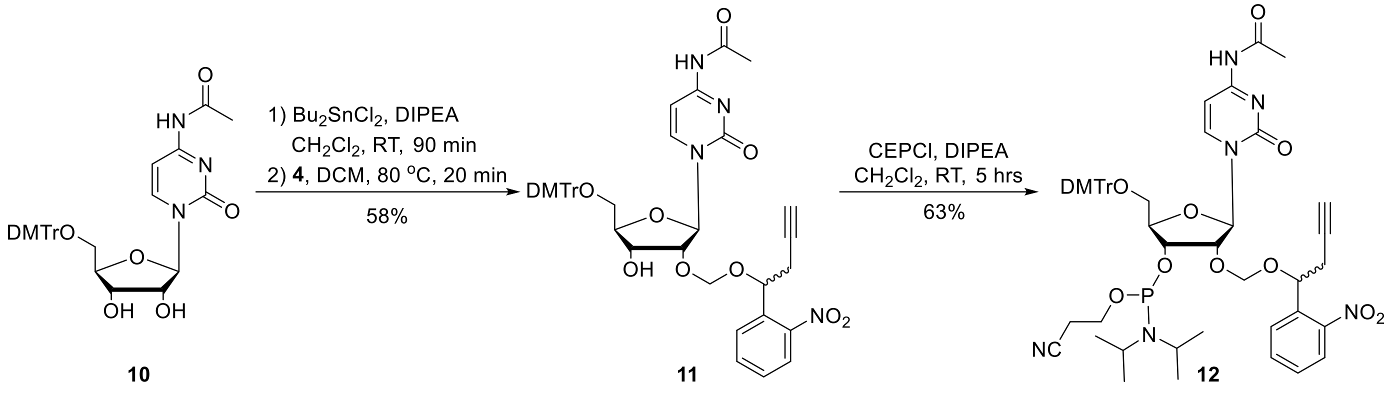


**Scheme S3**. Synthesis of photo-responsive cytidine phosphoramidite

To a solution of **10** (881 mg, 1.5 mmol, prepared according to literature[4]) in dry dichloromethane (25 mL) was added *N*,*N*-diisopropylethylamine (1.2 mL, 7.5 mmol) and dibutyltin dichloride (559 mg, 1.8 mmol), respectively. The mixture was stirred at room temperature for 2 hours and then **4** (crude product, 2.3 mmol) in dichloromethane (5 mL) was added dropwise. The reaction mixture was heated and stirred at 80 °C for 20 minutes, until TLC indicated the consumption of **10**. The reaction was quenched with aqueous sodium carbonate. The aqueous layer was extracted three times with dichloromethane. The organic phases were combined, dried over sodium sulfate and filtrated. The filtrate was evaporated to dryness and purified by column chromatography (dichloromethane/methanol 60/1) to afford the product **11** as a yellow foam (865 mg, yield 58%). $^{1}H$ NMR (400 MHz, DMSO-*d*$_6$): δ 10.93 (s, 1H), 8.27-8.21 (m, 1H), 7.99-7.95 (m, 1H), 7.81-7.70 (m, 2H), 7.58-7.53 (m, 1H), 7.41-7.25 (m, 9H), 6.91-6.89 (m, 4H), 5.76-5.73 (m, 1H), 5.52-5.36 (m, 2H), 5.12-4.97 (m, 1H), 4.79-4.68 (m, 1H), 4.33-4.16 (m, 2H), 4.02-4.00 (m, 1H), 3.76 (s, 6H), 3.33-3.31 (m, 2H), 2.86-2.73 (m, 3H), 2.11 (s, 3H). $^{13}C$ NMR (100 MHz, DMSO-*d*$_6$): δ 171.4, 162.9, 158.6, 154.7, 148.6, 148.5, 144.9, 135.9, 135.62, 135.57, 133.8, 133.7, 130.24, 130.17, 129.7, 129.5, 128.4, 128.2, 127.3, 124.6, 113.7, 95.9, 92.6, 89.7, 86.4, 82.3, 80.5, 80.4, 79.1, 78.9, 74.0, 73.9, 71.4, 71.3, 68.3, 67.9, 62.1, 61.9, 26.8, 26.4, 24.8. HRMS (ESI): *m/z* $[M + H]^+$ calcd for $C_{43}H_{43}N_4O_{11}$: 791.2923; found: 791.2909.

To a solution of intermediate **11** (406 mg, 0.5 mmol) in dry dichloromethane (10 mL) under nitrogen was added *N*,*N*-diisopropylethylamine (424 μL, 2.6 mmol). The mixture was stirred for 5 minutes, followed by the addition of 2-cyanoethoxy-*N*,*N*-diisopropylaminochlorophosphine (228 μL, 1.0 mmol). The reaction was quenched after 2 hours by adding aqueous sodium bicarbonate (15 mL). The aqueous layer was extracted three times with dichloromethane. The organic phases were combined, dried over sodium sulfate and filtrated. The filtrate was evaporated to dryness under reduced pressure and then purified by column chromatography (dichloromethane/ethyl acetate 1/1, with 0.5%

triethylamine) to afford the product **12** as yellow foam (319 mg, yield 63%). $^1$H NMR (500 MHz, $CDCl_3$): δ 9.16-9.12 (m, 1H), 8.45-8.23 (m, 1H), 7.88-7.80 (m, 2H), 7.57-7.53 (m, 1H), 7.36-7.19 (m, 9H), 6.82-6.77 (m, 4H), 6.03-5.87 (m, 1H), 5.59-5.50 (m, 1H), 5.14-5.04 (m, 1H), 4.79-4.62 (m, 1H), 4.42-4.07 (m, 3H), 3.75-3.74 (d, *J* = 3.5 Hz, 6H), 3.50-3.36 (m, 6H), 2.83-2.53 (m, 5H), 2.17-2.16 (m, 3H), 1.22-1.03 (m, 12H). $^{13}$C NMR (125 MHz, $CDCl_3$): δ 170.2, 162.5, 158.8, 155.3, 148.8, 148.6, 144.9, 144.2, 135.9, 135.6, 135.4, 135.3, 135.2, 133.4, 133.1, 130.4, 130.2, 129.6, 129.5, 128.6, 128.4, 128.0, 127.3, 124.3, 117.7, 113.4, 113.3, 96.5, 93.3, 92.8, 90.1, 89.6, 87.1, 87.0, 79.9, 79.7, 79.5, 78.9, 72.2, 71.6, 71.3, 71.23, 71.16, 60.9, 60.5, 58.4, 58.2, 47.2, 45.4, 45.3, 43.3, 27.1, 26.9, 25.0, 24.6, 24.5, 24.4, 23.0, 22.9, 20.2, 20.1. $^{31}$P NMR: δ 150.81. HRMS (ESI): *m/z* $[M + H]^+$ calcd for $C_{52}H_{59}N_6O_{12}PNa$: 1013.3821; found: 1013.3824.

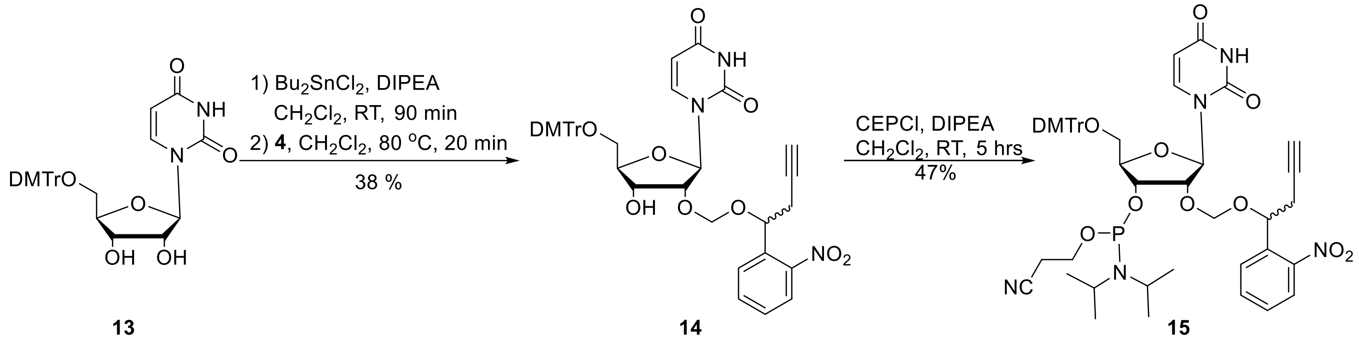


**Figure S4**. Synthesis of photo-responsive uridine phosphoramidite **15**

To a solution of **13** (819 mg, 1.5 mmol, prepared according to literature[5]) in dry dichloromethane (25 mL) was added *N*,*N*-diisopropylethylamine (1.2 mL, 7.5 mmol) and dibutyltin dichloride (559 mg, 1.8 mmol), respectively. The mixture was stirred at room temperature for 2 hours and then **4** (crude product, 2.3 mmol) in dichloromethane (5 mL) was added dropwise. The reaction mixture was heated and stirred at 80 °C for 20 minutes, until TLC indicated the consumption of **13**. The reaction was quenched with aqueous sodium carbonate. The aqueous layer was extracted three times with dichloromethane. The organic phases were combined, dried over sodium sulfate and filtrated. The filtrate was evaporated to dryness and purified by column chromatography (dichloromethane/ethyl acetate 3/1) to afford the product **14** as a yellow foam (423 mg, yield 38%). $^1$H NMR (400 MHz, DMSO-$d_6$): δ 11.4-11.3 (m, 1H), 7.99-7.93 (m, 1H), 7.78-7.52 (m, 4H), 7.39-7.30 (m, 4H), 7.26-7.24 (m, 5H), 6.91-6.89 (m, 4H), 5.84-5.68 (m, 1H), 5.42-5.34 (m, 2H), 5.25-5.20 (m, 1H), 4.94-4.87 (m, 1H), 4.73-4.60 (m, 1H), 4.28-4.13 (m, 2H), 3.93-3.92 (m, 1H), 3.74 (s, 1H), 3.25-3.22 (m, 2H), 2.77-2.73 (m, 2H), 2.71-2.69 (m, 1H). $^{13}$C NMR (125 MHz, DMSO-$d_6$): δ 163.4, 158.6, 150.7, 148.6, 145.2, 135.8, 135.6, 133.8, 130.3, 129.6, 128.4, 128.2, 124.7, 113.7, 102.3, 87.9, 86.4, 83.2, 80.3, 78.0, 74.0, 71.3, 69.0, 65.5, 63.3, 60.4, 27.0, 26.5, 14.6. HRMS (ESI): *m/z* $[M - H]^-$ calcd for $C_{41}H_{38}N_3O_{11}$: 748.2512; found: 748.2501.

To a solution of intermediate **14** (450 mg, 0.6 mmol) in dry dichloromethane (10 mL) under nitrogen was added *N*,*N*-diisopropylethylamine (496 μL, 3.0 mmol). The mixture was stirred for 5 minutes, followed by the addition of 2-cyanoethoxy-*N*,*N*-diisopropylaminochlorophosphine (268 μL, 1.2 mmol). The reaction was quenched after 2 hours by adding aqueous sodium bicarbonate (15 mL). The aqueous layer was extracted three times with dichloromethane. The organic phases were combined, dried over sodium sulfate and filtrated. The filtrate was evaporated to dryness under reduced pressure and then purified by column chromatography (dichloromethane/methanol 150/1, with 0.5% triethylamine) to afford the product **15** as yellow foam (270 mg, yield 47%). $^1$H NMR (500 MHz, $CDCl_3$): δ 7.90-7.84 (m, 2H), 7.78-7.75 (m, 1H), 7.57-7.51 (m, 1H), 7.40-7.30 (m, 3H), 7.24-7.19 (m, 7H), 6.79-6.77 (m, 4H), 6.02-5.85 (m, 1H), 5.47-5.45 (m, 1H), 5.27-5.09 (m, 1H), 4.97-4.87 (m, 1H), 4.72-4.55 (m, 1H), 4.48-4.31 (m, 2H), 4.17-4.10 (m, 1H), 3.73 (s, 6H), 3.55-3.30 (m, 6H), 2.80-2.30 (m, 5H), 1.11-1.03 (m, 12H). $^{13}$C NMR (125 MHz, $CDCl_3$): δ 163.0, 158.8, 150.09, 150.06, 150.0, 148.7, 148.5, 148.2, 148.0, 144.30, 144.27, 144.2, 140.1, 140.0, 136.0, 133.2, 130.30, 130.27, 129.3, 129.2, 129.1, 128.81, 128.77, 128.7, 128.3, 128.03, 128.01, 127.2, 124.5, 124.4, 117.8, 117.5, 117.4, 113.3, 102.33, 102.28, 93.4, 92.9, 92.6, 88.1, 88.0, 87.7, 87.2, 87.1, 82.9, 82.7, 82.6, 79.7, 79.6, 78.3, 71.5, 71.4, 71.3, 71.2, 71.1, 61.7, 61.5, 60.4, 58.6, 58.5, 58.0, 57.9, 49.4, 43.4, 43.34, 43.28, 43.2, 43.1, 30.7, 29.7, 29.6, 27.1, 26.9, 24.71, 24.65, 24.6, 24.51, 24.46, 21.1, 20.44, 20.38, 20.23, 20.17, 17.7, 14.2. $^{31}$P NMR: δ 150.76, 150.19. HRMS (ESI): *m/z* $[M + Na]^+$ calcd for $C_{50}H_{56}N_5O_{12}PNa$: 972.3555; found: 972.3552.

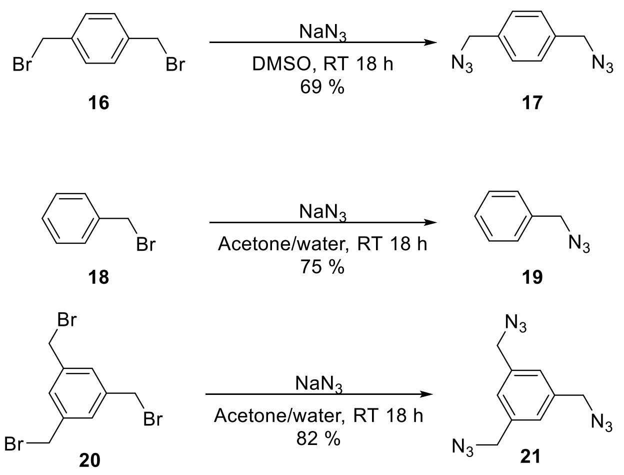


**Figure S5**. Synthesis of bis-azido-linker **17**, mono-azido linker **19** and tris-azido linker **21**

To a solution of sodium azide (163 mg, 2.5 mmol) in dimethyl sulfoxide (4 mL) was added 1,4-bis(bromomethyl) benzene **16** (264 mg, 1 mmol). The mixture was stirred until all the starting materials have been consumed. The reaction was quenched with water, and then extracted with diethyl ether. The organic phases were combined, washed with water (five times) to remove dimethyl sulfoxide, dried over sodium sulfate and then filtrated. The filtrate was evaporated under reduced pressure to afford the pure 1,4-bis(azidomethyl)benzene **17**.[6]

To a solution of bromomethyl benzene **18** (119 μL, 1 mmol) in a mixture of water/acetone (10 mL, v/v 1/4) was added sodium azide (97.5 mg, 1.5 mmol). The mixture was stirred until all the starting materials have been consumed. The mixture was extracted with dichloromethane, and the organic phases were combined, washed three times with water, dried over sodium sulfate and then filtrated. The filtrate was evaporated under reduced pressure to afford the pure azidomethyl benzene **19**.[7]

To a solution of sodium azide (195 mg, 6 mmol) in a mixture of water/acetone (10 mL, v/v 1/4) was added 1,3,5-tris(bromomethyl) benzene **20** (178.45 mg, 1 mmol). The mixture was stirred until all the starting materials have been consumed. The reaction was quenched with water, and then extracted with dichloromethane. The organic phases were combined, dried over sodium sulfate and then filtrated. The filtrate was evaporated under reduced pressure to afford the pure 1,3,5-tris(azidomethyl)benzene **21**.[8]

**3. Preparation of oligonucleotides**

CrRNAs and tracrRNA were optimized for guide RNA according to literature.[9] Unmodified crRNAs were purchased from Biosyntech (Suzhou) and tracrRNA (Alt-R® tracrRNA) was purchased from Integrated DNA Technologies (IDT, Coralville, IA). Modified crRNAs were chemically synthesized by a Bioautomation MerMade 12 DNA/RNA Synthesizer according to the standard solid phase synthesis protocol. The synthesized RNA oligonucleotides were purified by HPLC (PLRP-S column, 250 mm × 4.6 mm, 100 Å, 8 μm; elute A = triethylammonium acetate (0.1 M, pH 7), B = methanol, gradient elution: 15% to 85% methanol in 25 minutes, flow 1 mL·min$^{-1}$). Desalting purification was performed by Amicon® Ultra Centrifugal Filters (Millipore) according to the manufacturer’s instructions. Usually, 3-5 washes were required. Finally, the RNA oligonucleotides solutions were collected, lyophilized and stored at −20 °C or below. Their molecular weights were confirmed by ESI-MS (Sangon Biotech).

**4. General procedure for oligonucleotide circularization**

To an aqueous solution of modified crRNAs (1.5 nmol) in a reaction tube were added in following order: triethylammonium acetate buffer (2 M, 2 μL, pH 7), bis-azido-linker (0.75 mM in water/dimethyl sulfoxide/*tert*-butyl alcohol 4/3/1, 2 μL), dimethyl sulfoxide (55% of the final solution, v/v), magnesium chloride (100 mM, 1 μL) and fresh ascorbic acid (125 mM, 2 μL). After the oligonucleotide solution was degassed with nitrogen, a solution of copper sulfate-tris(3-hydroxypropyltriazolylmethyl)amine (250 mM in dimethyl sulfoxide/water, 1 μL, 55% v/v) was added and the mixture (final volume = 20 μL) was left for 4 hours at room temperature. The solution was then diluted with RNase-free water and desalted using an Amicon Ultra Centrifugal Filter (Millipore). Usually, 6 washes were required. Then oligonucleotides were purified by HPLC (PLRP-S column, 250 mm × 4.6 mm, 100 Å, 8 μm; elute A = triethylammonium acetate (0.1 M, pH 7), B = methanol, gradient elution: 15% to 85% methanol in 25 minutes, flow 1 mL·min$^{-1}$) and analyzed by 15% denatured PAGE gel. The procedures of click reaction with mono-azido linker and tris-azido linker were similar, except that the amount of mono-azido linker was needed twice that of the modified crRNAs.

**5. Decaging assay of circular crRNAs**

A sample of the circular RNA (30 μL, 3 μg) was irradiated with UV light (365 nm, 20 mW·cm$^{-2}$) for different durations (0 second, 30 seconds, 1 minute, 2 minutes, 3 minutes and 5 minutes, respectively) and then analyzed by 15% denatured PAGE gel (170 V, 45 minutes).

**6. crRNA/sgRNA stability assay**

This assay was performed in 1× NEB 3.1 buffer in a reaction volume of 100 μL. The unmodified gRNAs (crRNA or sgRNA)/linear modified gRNAs (crRNA-**a** or sgRNA-**a**)/circular gRNAs (crRNA-**c** or sgRNA-**c**) (10 μM, 10 μL) was incubated in the presence of RNase A (SIGMA, R4642) in different concentrations (0.01 mg·L$^{-1}$ or 0.1 mg·L$^{-1}$) at 37 °C. Each 15 μL of the reaction solution was extracted after incubation at 1, 2.5, 5, 10, 30 and 60 minutes, respectively. 5 μL of ethylenediaminetetraacetic acid (0.5 M) was added to quench each solution, and the mixture was quickly immersed in dry ice. All RNA samples were analyzed by 15% native polyacrylamide gel electrophoresis (PAGE) gel (250 V, 30 minutes).

**7. Expression and purification of Cas9/Cpf1 protein**

The plasmid of the coding sequence for SpyCas9 (Plasmid #62933) and AsCpf1 (Plasmid #79007) were gifts from Prof. Wang's group. Each protein was expressed in the *E.coli* BL21 (DE3) cells. Briefly, cells were grown at 37 °C to $OD_{600}$ > 0.6. IPTG (Sigma, 0.2 mM) was then added and the culture temperature was lowered to 20 °C. Cells were grown overnight and harvested by centrifugation at 6000 rpm at 4 °C and lysed by sonication in a buffer containing 20 mM Tris-HCl, pH 7.5 and 500 mM sodium chloride. The protein was then purified *via* Ni-NTA, collected and concentrated using

100 K Amicon Ultra column (Millipore). Purity of protein was analyzed by SDS-polyacrylamide gel electrophoresis.

**8. *In vitro* DNA cleavage assay**

Target DNA were amplified by polymerase chain reaction from pEGFP-N1 vector using the following primer pairs: EGFP-FW1: 5'-TGGTGAGCAAGGGCGAGGAG-3', EGFP-RV1: 5'-GTCCTCGATGTTGTGGCGGAT-3' and then purified using Cycle Pure Kit (Gene-protein Link Biotech). The prepared crRNA/tracrRNA duplex (each 10 pmol) were incubated with 10 pmol Cas9 and 100 ng target DNA with 1 × NEB buffer 3.1 (NEB) in RNase-free water for 90 minutes at 37 °C. Cas9 protein was degraded by incubating with proteinase K at 37 °C for 15 minutes, and protease K (TransGen Biotech) was inactivated by heating at 95 °C for 5 minutes finally. DNA cleavage product was analyzed by 2% agarose gel electrophoresis. For light activation assay, the circular crRNA/tracrRNA duplex was irradiated with UV light (365 nm, 20 mW·cm$^{-1}$) for 5 minutes before incubated with Cas9.

The steps of related vitro DNA cleavage assay for single gRNAs (*GFP*-Cpf1-sgRNA or *MSTN*-sgRNA) were similar. The corresponding target DNAs were obtained by polymerase chain reaction from pEGFP-N1 vector or MSTN-pMDTM18-T vector using the following primer pairs: EGFP-FW2: 5'-CGGTTTGACTCACGGGGATT-3', EGFP-RV2: 5'-CTCGATGTTGTGGCGGATCT-3'; MSTN-FW1: 5'-TCATTTTTCATAAAAATGATCT-3', MSTN-RV1: 5'-ATAAGCACAGGAAACTGGTAGT-3'. The sgRNAs or modified sgRNAs (each 5 pmol) were incubated with 5 pmol of Cpf1/Cas9 and 100 ng of target DNA with 1 × NEB buffer 3.1 (NEB) in RNase-free water for 30 minutes at 37°C. Cpf1/Cas9 proteins were degraded by incubating with proteinase K at 37 °C for 15 minutes, and protease K was inactivated by heating at 95 °C for 5 minutes finally. DNA cleavage product was analyzed by 2 % agarose gel electrophoresis.

**9. Binding assay of circular gRNAs**

The unmodified or circular crRNA and tracrRNA (each 10 pmol) were mixed with dCas9 protein (8 pmol) and FAM-dsDNA (1 pmol) in the 1 x NEB buffer 3.1. The mixture was incubated at 37 °C for 15 minutes, and then analyzed by electrophoresis in 5% natured polyacrylamide gel. For light activation assay, the circular crRNA/tracrRNA duplex was irradiated with UV light (365 nm, 20 mW·cm$^{-1}$) for 5 minutes before incubated with dCas9.

To measure the thermostability of circular crRNA/DNA hybrid, modified and unmodified crRNA/complementary DNA were mixed in 1 × PBS to the final concentration of 5 μM. Oligonucleotide solutions were hybridized by first heating at 65 °C for 15 minutes and then slowly cooled down to room temperature. The maximum absorbance of CD spectrum was recorded by Jasco J-815 along with the increasing temperature from 25 to 90 °C. The heating rate was 1 °C·min$^{-1}$. Melting temperatures were calculated by measuring the CD(t)/CD(25 °C) at each temperature with GraphPad Prism 8.

**10. Light-activated exogenous GFP gene editing in cells**

GFP-stably expressed HEK cells (HEK-GFP cells) were cultured at 37 °C, 5 % $CO_2$, and 95 % humidity in DMEM culture medium (10 % FBS, BI; 1 × Penicillin/Streptomycin, Gibco) and were seeded in 48-well plate one day overnight before experiment (25 k cells per well). 100 ng of Cas9 mRNA was mixed with different doses of GFP-targeting gRNAs and 0.8 μL of Lipofectamine 3000 (LPF 3k) in 50 μL of free DMEM, followed by 15 minutes of incubation at room temperature. The LPF 3k/Cas9 mRNA/gRNAs complexes were then added to cells and incubated for 10 hours before changing fresh cell culture medium. The GFP expression profile was quantified using flow cytometry analysis (CytoFLEX S) at 48 hours post Cas9 mRNA delivery, and normalized to cells without Cas9 mRNA/gRNA delivery to determine genome editing efficiency. For light-activated gene editing, HEK-GFP cells were transfected with Cas9 mRNA and modified gRNAs complexes as described above. After the transfection mixture was replaced with fresh cell culture medium, the cells were subjected to 5 minutes of 365 nm irradiation by a UV lamp (10 mW·cm$^{-2}$). For the spatial control experiments, UV irradiations were performed through a tin foil mask to only expose a subset of cells to 365 nm light for 5 minutes. The GFP expression profile was quantified using flow cytometry analysis at 48 hours post Cas9 mRNA delivery, and normalized to cells without Cas9 mRNA/gRNA delivery to determine genome editing efficiency.

**11. Light-activated endogenous VEGFA gene editing in cells**

293T-Cas9 cells stably expressing Cas9 protein (UBIGENE) were cultured in 12-well plate (50 k cells per well) in DMEM with 10% FBS overnight before transfection and washed with DPBS buffer. Unmodified crRNA, linear crRNA-a or circular crRNA-c were 1:1 premixed with tracrRNA in Nuclease Free Duplex Buffer (Integrated DNA Technologies) respectively. Pre-transfection experiments were performed using different doses (10/20/30/40 pmol per well respectively) of unmodified crRNA/tracrRNA duplexes and Lipofectamine 3000 (1/2/3/4 μL per well respectively) in 100 μL Opti-MEM (Gibco) for 15 minutes at room temperature before mixing with 900 μL of 293T-Cas9 cells. The medium was changed to complete DMEM after an incubation of 6 hours and the cells were further cultured another 72 hours at 37 °C. Genomic DNAs were extracted using TIANamp Genomic DNA Kit for polymerase chain reaction (Vazyme) to amplify the target DNA fragments. The primer pairs were as follows: VEGFA-FW: 5'-CAGGGCTGGGGCTGTTCTCATAC-3'; VEGFA-RV: 5'-GTATGTGGGTGGGTGTGTCTACAG-3'. The purity of the PCR products were characterized by 2 % agarose gel electrophoresis. The T7E1 assays of the PCR products were performed according to manufacturer's protocol. Products were analyzed by 2 % agarose gel electrophoresis. After determining 10 pmol·mL$^{-1}$ as the optimal transfection concentration, it was then used as the transfection concentration of the further experiments. For light-activated gene editing, 293T-Cas9 cells were transfected with crRNA/tracrRNA duplexes as described above. After the transfection mixture was replaced with fresh cell culture medium, the cells were subjected to 5 minutes of 365 nm irradiation by a UV lamp (365 nm, 10 mW·cm$^{-2}$). The PCR amplicons were also sequenced by Sanger sequencing using primer 5'-CAGGGCTGGGGCTGTTCTCATAC-3'.

**12. Light-activated MSTN gene editing in zygotes**

The female mice (6-week old ICR mice from Beijing Vital River Experimental Animals Centre) were injected intraperitoneally with 10 International Units (IU) of pregnant mare serum gonadotropin (PMSG) (Ningbo Sansheng Biological Technology, Ningbo, China), followed by injection with 10 IU of human chorionic gonadotropin (hCG) (Ningbo Sansheng Biological Technology) 46-48 hours later. The female mice were immediately caged with mature male mice and zygotes were harvested from oviducts ~20 hours after hCG injection. After several washes, the zygotes were placed into drops of M2 medium (Sigma-Aldrich). RNPs were prepared to contain 2 μg of Cas9 protein and 1 μg of different sgRNAs in 1 x TE buffer (to make the final volume 20 μL). The zygotes were microinjected with ~10 pL of the RNPs solution using a microinjection pump (Eppendorf Femtojet) under an inverted fluorescence microscope (Nikon TE300, Japan). The *hv*+ group was followed by irradiation with ultraviolet light (excitation wavelengths of 365 nm, 10 mW·cm$^{-2}$) for 90 seconds and others were incubated in dark. After the injections were completed, the zygotes were transferred to 50 μL-drop of KSOM medium (Sigma-Aldrich) and cultured to blastocyst stage, in humidified 5% $CO_2$ at 37 °C. The embryos were harvested and placed into PCR reaction solution, and the target genes were amplified for sequencing analysis. Two rounds of continuous PCR were used for amplification, because only one amplification was likely to be insufficient for sequencing. The primer pairs were as follows: MSTN-FW2: 5'-CGGCACGCTGTCTCTTAGTCTC-3', MSTN-RV2: 5'-TGGGTACACACCTACCTTTGGAGTA-3'; MSTN-FW3: 5'-AAATATCCCTTAGCCCAGAGTTCTC-3', MSTN-RV3: 5'-GCAACACTGTCTTCACATCAATACT-3'.

**13. Supplementary tables and figures**

**Table S1: DNA and RNA sequences used in the study**

(Blue bold letters indicate the modification sites substituted with NPBOM, and the red lines indicate the NPBOM derived linkage upon reacting with 1,4-bis(azidomethyl)benzene, while the ochre red bold letters indicate the NPBOM modification sites reacted with benzyl azide)

| Names | Sequences (5’-3’) | calcd MW | found MW |
|---|---|---|---|
| GFP-crRNA | GGGCACGGGCAGCUUGCCGGGUUUUAGAGCUAUGCU | 11613.9 | 11613.6 |
| GFP-crRNA-a1 | GGGC**A**CGGGCAGCUUGCCGGGUUUU**A**GAGCUAUGCU | 12023.7 | 12023.3 |
| GFP-crRNA-c1 | GGGC**A**CGGGCAGCUUGCCGGGUUUU**A**GAGCUAUGCU | 12207.7 | 12209.3 |
| GFP-crRNA-a2 | GGGCACGGGC**A**GCUUGCCGGGUUUU**A**GAGCUAUGCU | 12023.7 | 12020.9 |
| GFP-crRNA-c2 | GGGCACGGGC**A**GCUUGCCGGGUUUU**A**GAGCUAUGCU | 12207.7 | 12217.0 |
| GFP-crRNA-a1’ | GGGC**A**CGGGCAGCUUGCCGGGUUUU**A**GAGCUAUGCU<br>**A** = 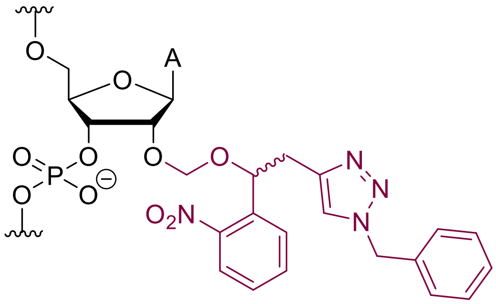 | | |
| GFP-crRNA-a2’ | GGGCACGGGC**A**GCUUGCCGGGUUUU**A**GAGCUAUGCU | | |
| VEGFA-crRNA | UCAGAUGUGACAAGCCGAGGGUUUUAGAGCUAUGCU | 11590.9 | 11594.3 |
| VEGFA-crRNA-a | UCAGAUGUG**A**CAAGCCGAGGGUUUU**A**GAGCUAUGCU | 12000.7 | 12001.4 |
| VEGFA-crRNA-c | UCAGAUGUG**A**CAAGCCGAGGGUUUU**A**GAGCUAUGCU | 12184.8 | 12210.3 |
| GFP-cpf1-sgRNA | AAUUUCUACUCUUGUAGAUCGUCGCCGUCCAGCUCG<br>ACCAGGA | 13649.1 | 13653.2 |
| GFP-cpf1-sgRNA-a1 | **A**AUUUCUACUCUUGUAGAUCGUCGCCGUCCAGCUCG<br>ACCAGG**A** | | |
| GFP-Cpf1-sgRNA-c1 | **A**AUUUCUACUCUUGUAGAUCGUCGCCGUCCAGCUCG<br>ACCAGG**A** | 14238.8 | 14244.3 |
| GFP-Cpf1-sgRNA-a2 | **A**AUUUCUACUCUUGUAGA**U**CGUCGCCGUCCAGCUCG<br>ACCAGGA | 14054.1 | 14059.8 |

| | | | |
|---|---|---|---|
| GFP-Cpf1-sgRNA-c2 | **A**AUUUCUACUCUUGUAGA**U**CGUCGCCGUCCAGCUCG<br>ACCAGGA | 14238.8 | 14210.3 |
| GFP-Cpf1-sgRNA-a3 | AAUUUCUACUCUUGUAGAU**C**GUCGCCGUCCAGCUCG<br>ACCAGG**A** | 14054.1 | 14059.8 |
| GFP-Cpf1-sgRNA-c3 | AAUUUCUACUCUUGUAGAU**C**GUCGCCGUCCAGCUCG<br>ACCAGG**A** | 14238.8 | 14216.0 |
| *MSTN*-sgRNA | AAAGACGGUACAAGGUAUACGUUUUAGAGCUAGAAAUAGCAA<br>GUUAAAAUAAGGCUAGUCCGUUAUCAACUUGAAAAAGUGGCA<br>CCGAGUCGGUGCUUUU | 32283.2 | 32296.4 |
| *MSTN*-sgRNA-a1 | **A**AAGACGGUACAAGGUAUACGUUUUAGAGCUAGAAAUAGCAA<br>GUUAAAAUAAGGCUAGUCCGUUAUCAACUUGAAAAAGUGGCA<br>CCGAGUCGGUGCUUU**U** | 32693.0 | 32703.7 |
| *MSTN*-sgRNA-c1 | **A**AAGACGGUACAAGGUAUACGUUUUAGAGCUAGAAAUAGCAA<br>GUUAAAAUAAGGCUAGUCCGUUAUCAACUUGAAAAAGUGGCA<br>CCGAGUCGGUGCUUU**U** | | |
| *MSTN*-sgRNA-a2 | AAAGACGGUACAAGG**U**AUACGUUUUAGAGCUAGAAAUAGCAA<br>GU**U**AAAAUAAGGCUAGUCCGUUAUCAACUUGAAAAAG**U**GGCA<br>CCGAGUCGGUGCUUUU | 32897.9 | 32905.8 |
| *MSTN*-sgRNA-c2 | AAAGACGGUACAAGG**U**AUACGUUUUAGAGCUAGAAAUAGCAA<br>GU**U**AAAAUAAGGCUAGUCCGUUAUCAACUUGAAAAAG**U**GGCA<br>CCGAGUCGGUGCUUUU | | |
| FAM-ssDNA1 | FAM-GGTGGTCACGAGGGTGGGCCAGGGCACGGGCAGCTTGCCGGTGGTGCAGATGAACTTCA | | |
| ssDNA2 | TGAAGTTCATCTGCACCACCGGCAAGCTGCCCGTGCCCTGGCCCACCCTCGTGACCACC | | |
| EGFP-FW1 | TGGTGAGCAAGGGCGAGGAG | | |
| EGFP-RV1 | GTCCTCGATGTTGTGGCGGAT | | |
| EGFP-FW2 | CGGTTTGACTCACGGGGATT | | |
| EGFP-RV2 | CTCGATGTTGTGGCGGATCT | | |
| VEGFA-FW | CAGGGCTGGGGCTGTTCTCATAC | | |
| VEGFA-RV | GTATGTGGGTGGGTGTGTCTACAG | | |

| | |
|---|---|
| MSTN-FW1 | TCATTTTTCATAAAAATGATCT |
| MSTN-RV1 | ATAAGCACAGGAAACTGGTAGT |
| MSTN-FW2 | CGGCACGCTGTCTCTTAGTCTC |
| MSTN-RV2 | TGGGTACACACCTACCTTTGGAGTA |
| MSTN-FW3 | AAATATCCCTTAGCCCAGAGTTCTC |
| MSTN-RV3 | GCAACACTGTCTTCACATCAATACT |

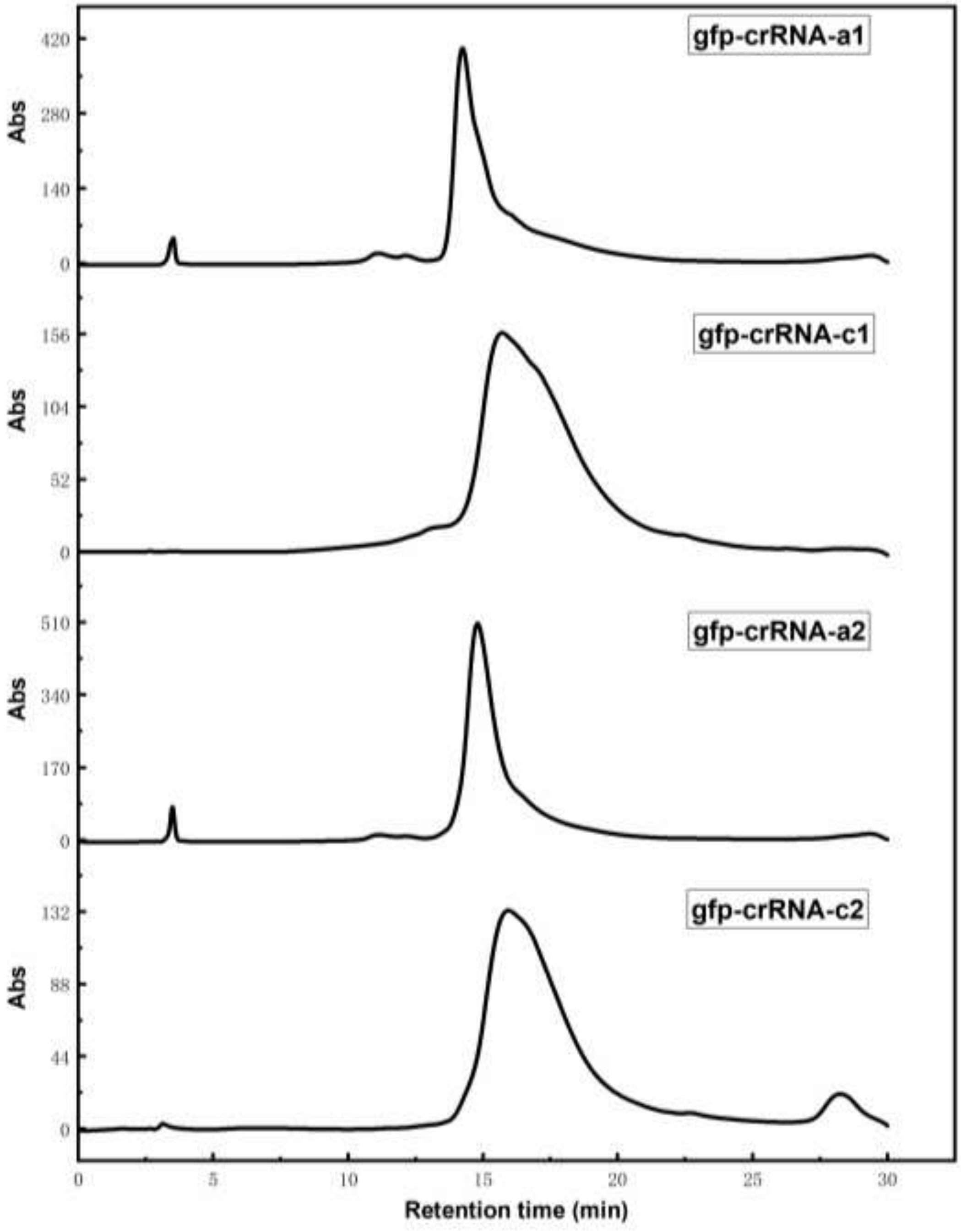

**Figure S6**. Monitoring of *GFP*-crRNA circularization reaction by RP-HPLC (PLRP-S column, 250 mm × 4.6 mm, 100 Å, 8 μm; A = 0.1 M triethylammonium acetate pH 7, B = methanol, 15 % to 85 % methanol in 25 minutes, flow 1 mL·min$^{-1}$). The retention time of circular crRNAs was longer than linear crRNAs.

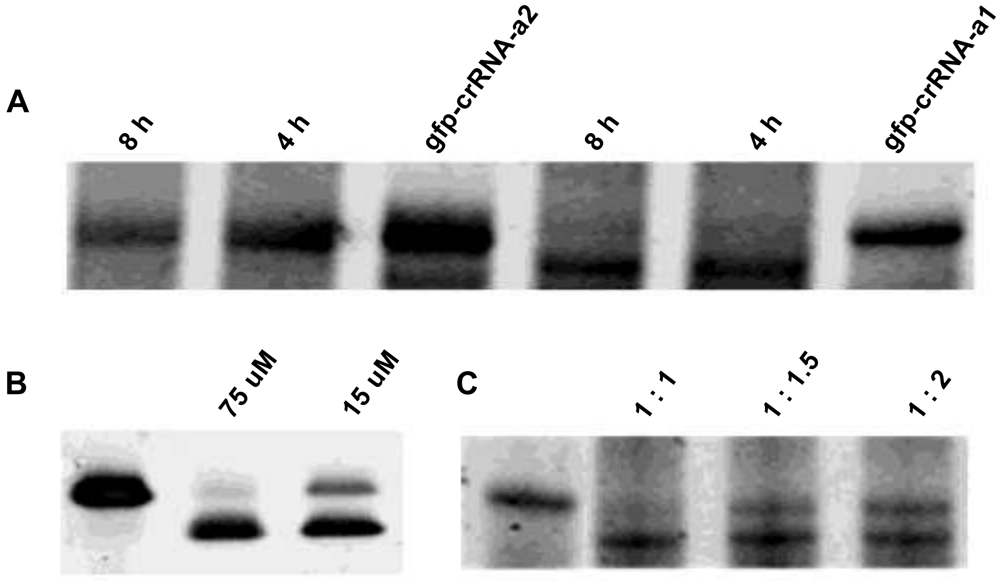


**Figure S7**. Optimization of circularization conditions. A) Reaction results of circularization of the linear modified gRNAs at A) different reaction times, 4 h or 8 h respectively, B) different reaction concentrations, C) different ratios of reactants (gRNA: bis(azidomethyl)benzene).

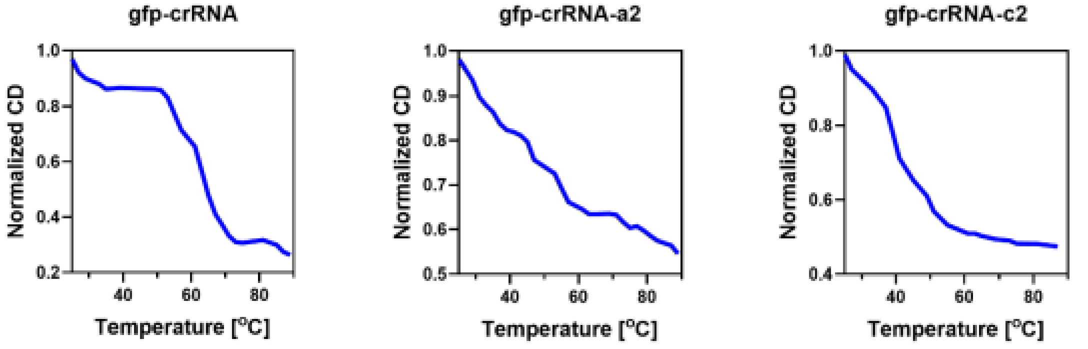


**Figure S8**. Normalized CD detection of *GFP*-crRNA, *GFP*-crRNA-**a2**, *GFP*-crRNA-**c2**. The melting temperature is 64 °C, 55 °C, 42 °C, respectively. Modified and unmodified crRNA/complementary DNA were mixed in 1×PBS to the final concentration of 5 μM. Oligonucleotide solutions were hybridized by first heating at 65 °C for 15 min and then slowly cooled down to room temperature. The maximum absorbance of CD spectrum was recorded by Jasco J-815 along with the increasing temperature from 25 to 90 °C. The heating rate was 1 °C·min$^{-1}$.

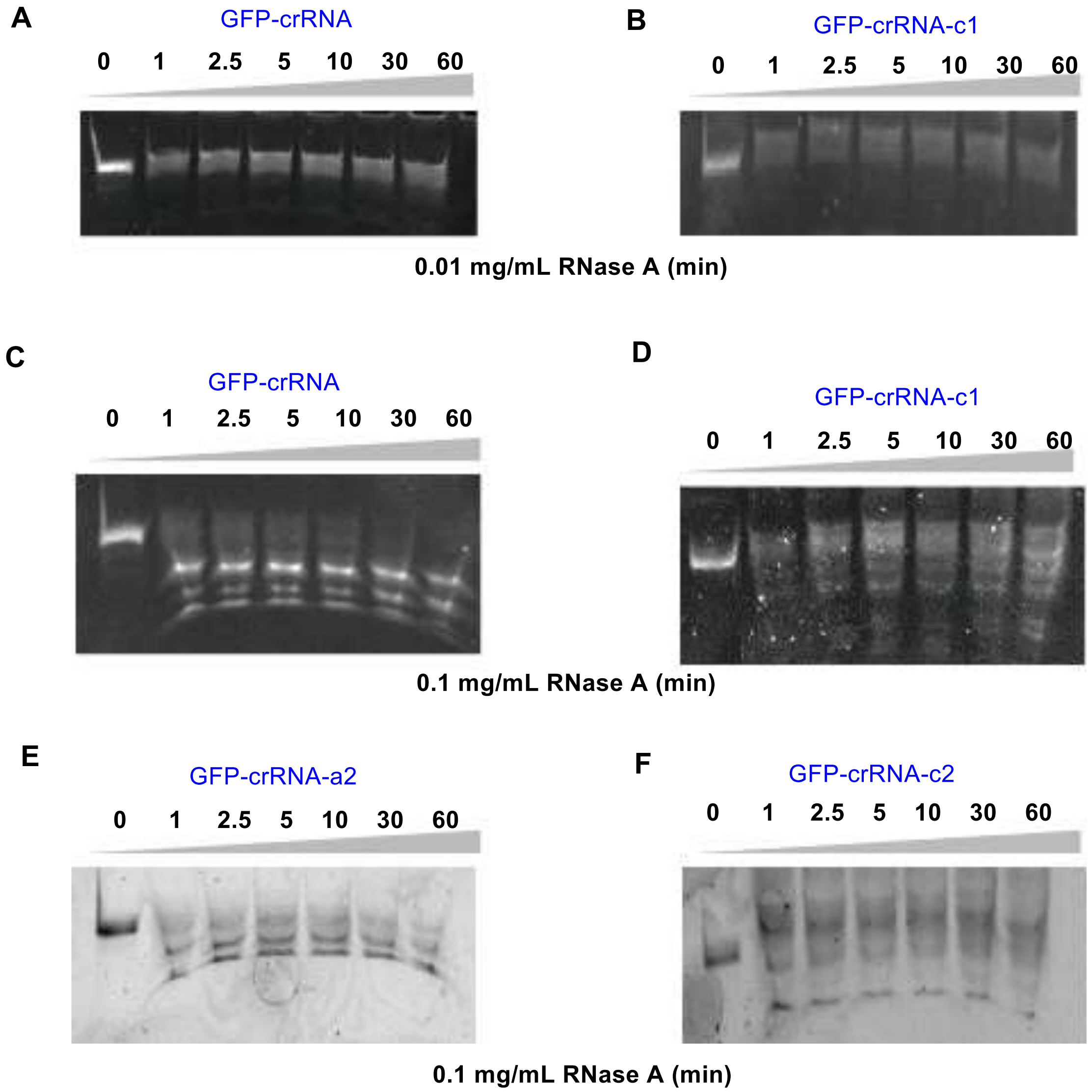


**Figure S9**. Comparison of stability between different crRNAs under RNase A. A) *GFP*-crRNA in 0.01 mg·L$^{-1}$ RNase A, B) *GFP*-crRNA-**c1** in 0.01 mg·L$^{-1}$ RNase A, C) *GFP*-crRNA in 0.1 mg·L$^{-1}$ RNase A, D) *GFP*-crRNA-**c1** in 0.1 mg·L$^{-1}$ RNase A, E) *GFP*-crRNA-**a2** in 0.1 mg·L$^{-1}$ RNase A, F) *GFP*-crRNA-**c2** in 0.1 mg·L$^{-1}$ RNase A. The unmodified/ circular modified crRNA (10 μM, 10 μL) was incubated in the presence of RNase A in different concentrations (0.01 mg·L$^{-1}$ or 0.1 mg·L$^{-1}$) at 37 °C for 0, 1, 2.5, 5, 10, 30, 60 minutes, respectively.

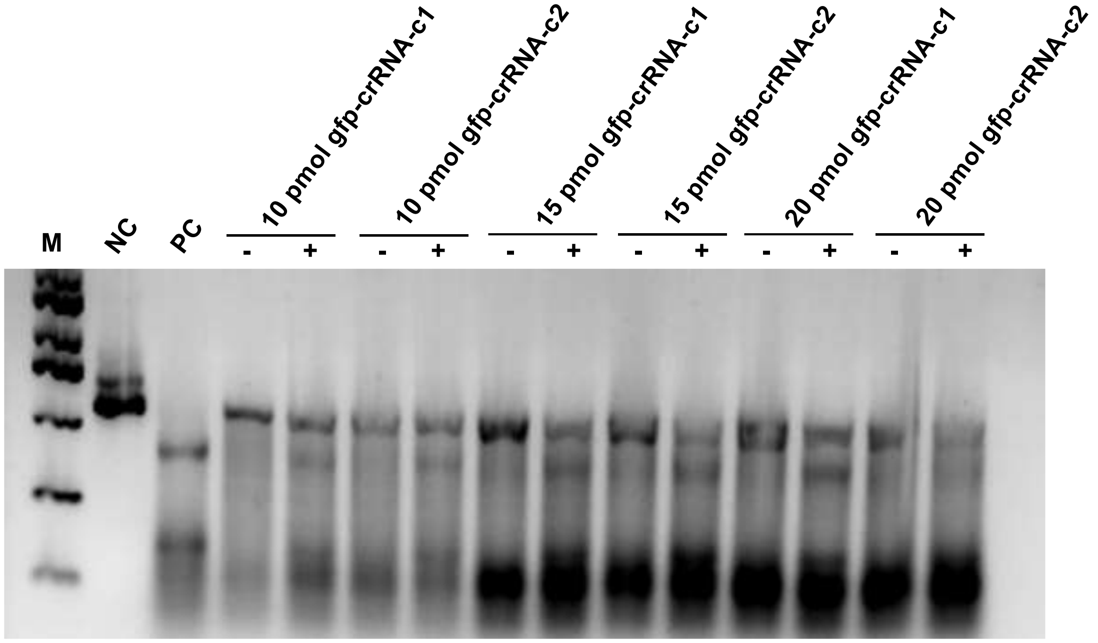


**Figure S10**. *In vitro* DNA cleavage assay at different crRNA's concentrations. Negative control (NC) without crRNA; positive control (PC) transfected with unmodified *GFP*-crRNA:tracrRNA duplex. "+", "-" were with or without UV (365 nm, 20 mW·cm$^{-2}$). The concentrations of tracrRNA and Cas9 were also increased according to the ratio of 1:1:1. Increasing the amount of circular crRNA had no obvious background cleavage before light exposure, which basically did not affect the light regulation ability of gene editing.

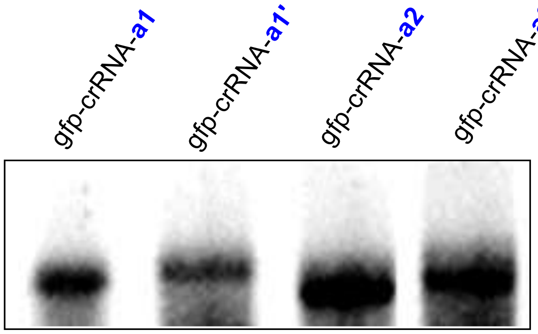


**Figure S11.** PAGE analysis of the linear *GFP*-crRNA-a and two linear and di-modified crRNAs *GFP*-crRNA-a'.

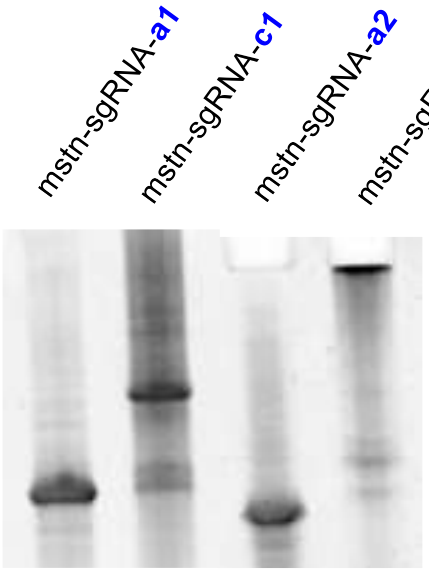


**Figure S12.** PAGE analysis of the linear *MSTN*-sgRNA-a and circular crRNAs *MSTN*-sgRNA-c.

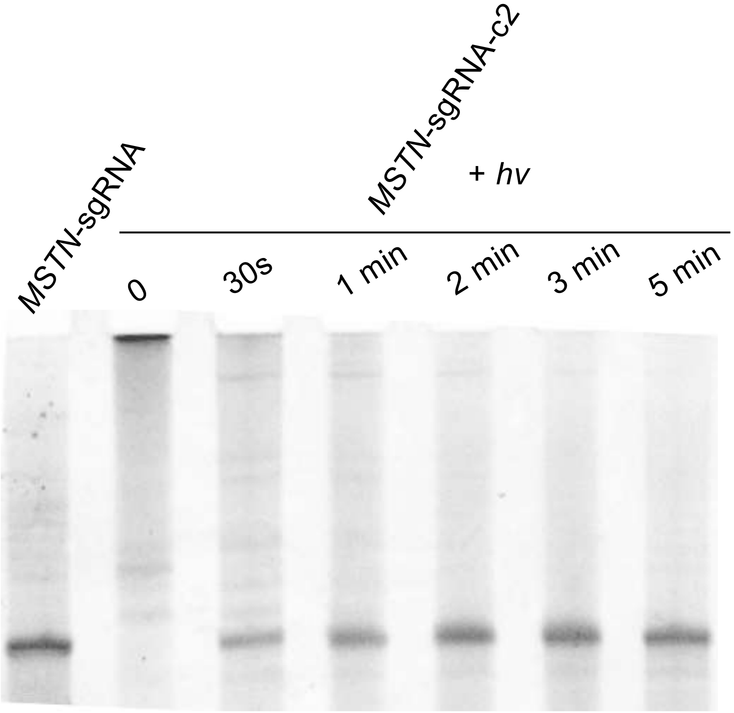


**Figure S13.** Comparison of native MSTN-sgRNA and light-mediated uncaging reaction of *MSTN*-sgRNA-c2 monitored by gel images at various time points. A sample of the circular RNA (30 μL, 3 μg) was irradiated with UV light (365 nm, 20 mW·cm$^{-2}$) for different durations (0 second, 30 seconds, 1 minute, 2 minutes, 3 minutes and 5 minutes, respectively) and then analyzed by 15% denatured PAGE gel (170 V, 45 minutes).

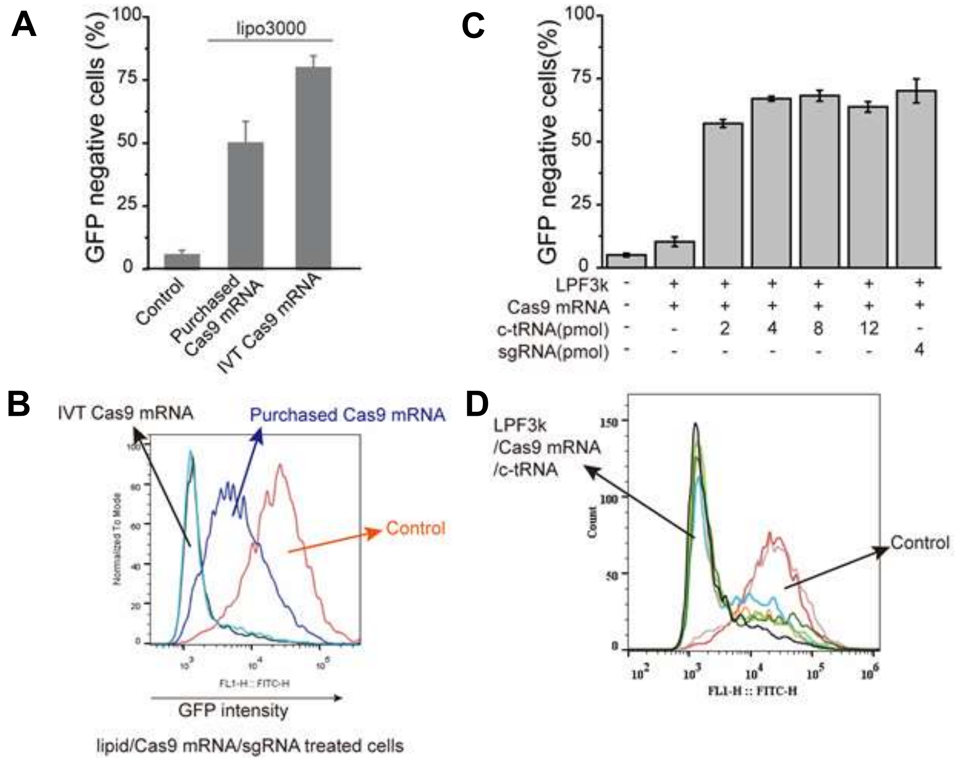


**Figure S14**. In cell CRISPR-Cas9 experiment with Cas9-mRNA delivery. A) Comparison of gene editing effects between cas9-mRNAs from different sources. IVT Cas9 mRNA was produced using the mMESSAGE mMACHINE® T7 Ultra Kit (Catalog Number AM1345, ThermoFisher Scientific, USA); Purchased Cas9 mRNA was purchased from APExBIO. B) Green fluorescence intensity of cells with Cas9-mRNAs and sgRNA delivery. After delivery of both purchased Cas9 mRNA and IVT-Cas9 mRNA, the fluorescence intensity of cells was significantly reduced compared to the control group, indicating gene editing. C) Effect of crRNA:tracrRNA dose on GFP knockdown when Cas9 mRNA dose was held constant at 100 ng. c-tRNA referred to the duplex of crRNA and tracrRNA; sgRNA referred to the single guide RNA. D) Green fluorescence intensity of cells with crRNA:tracrRNA duplex and IVT Cas9-mRNA co-delivery.

A

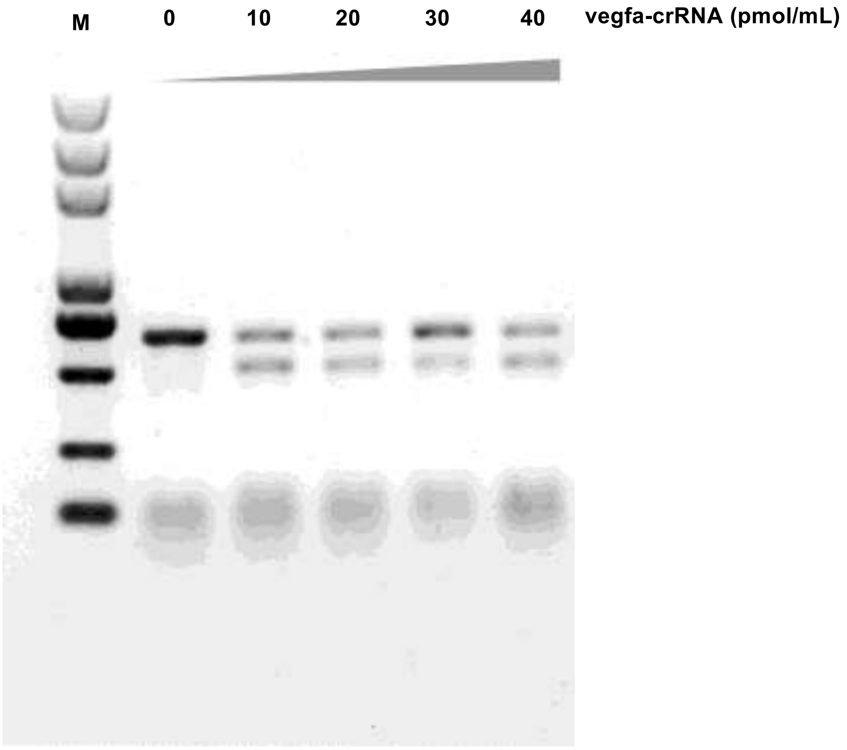


B

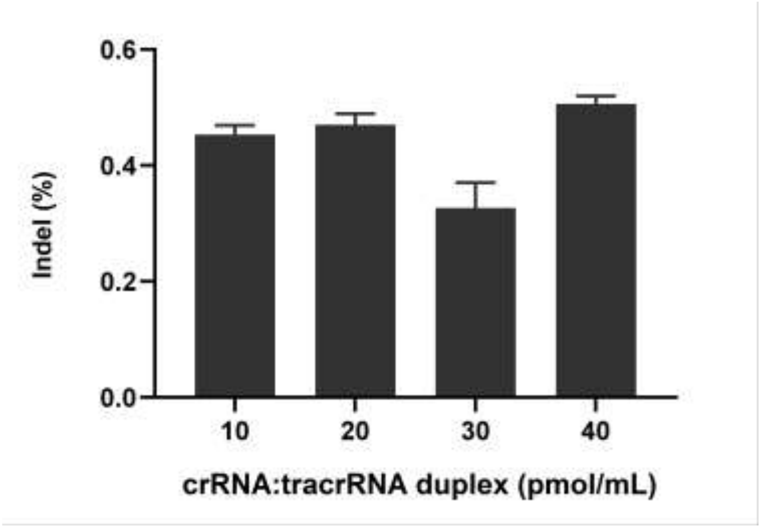


**Figure S15**. Indel mutation efficiency measured by T7E1 assay. Different doses of crRNA:tracrRNA duplex were delivered to 293T-Cas9 cells, indicating 10 pmol·mL$^{-1}$ of crRNA:tracrRNA duplex was enough to gene editing.

A

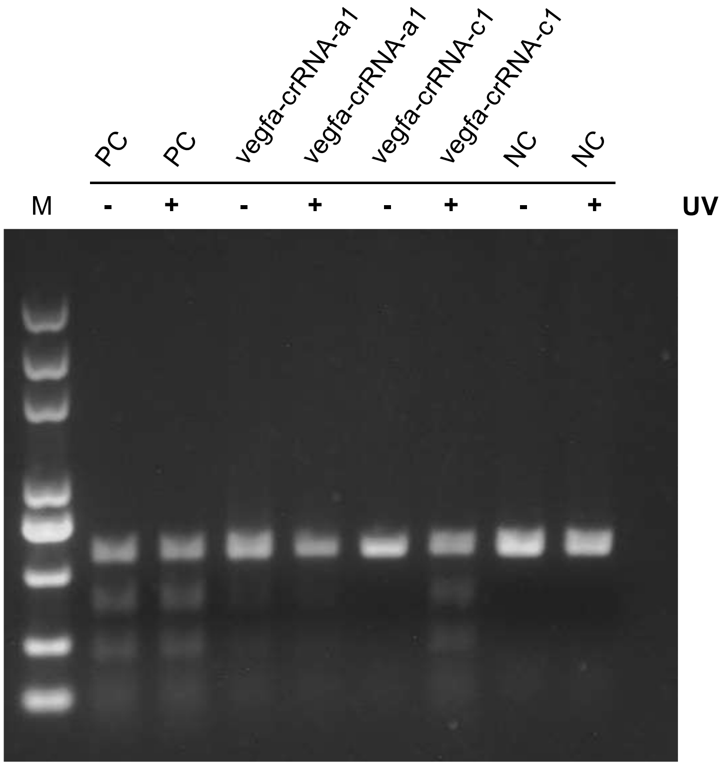


B

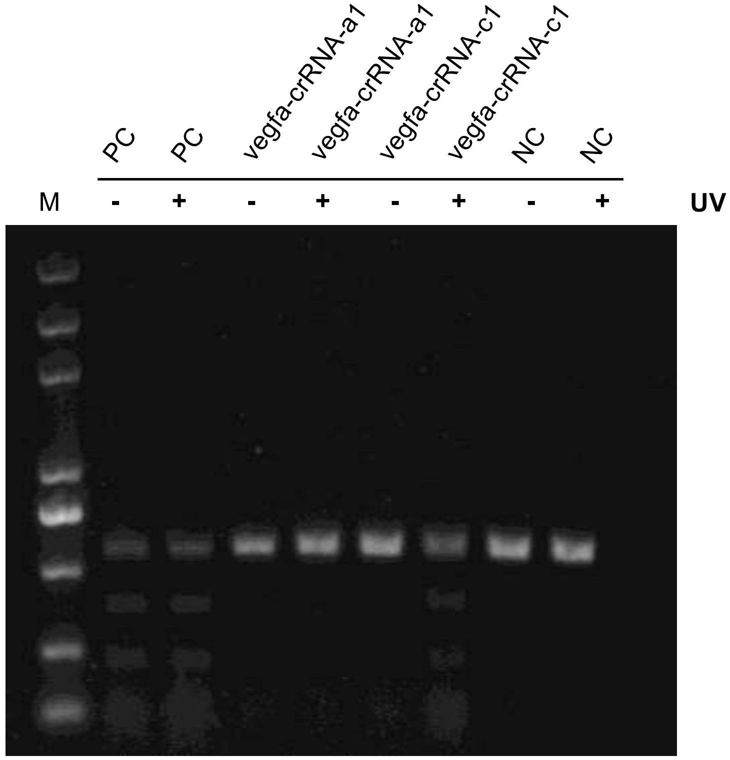


**Figure S16**. Indel mutation efficiency measured by T7E1 assay. Negative control (NC) without crRNA; positive control (PC) transfected with unmodified vegfa-crRNA:tracrRNA duplex. “+”, “-” were with or without UV (365 nm, 10 mW·cm$^{-1}$).

**A**

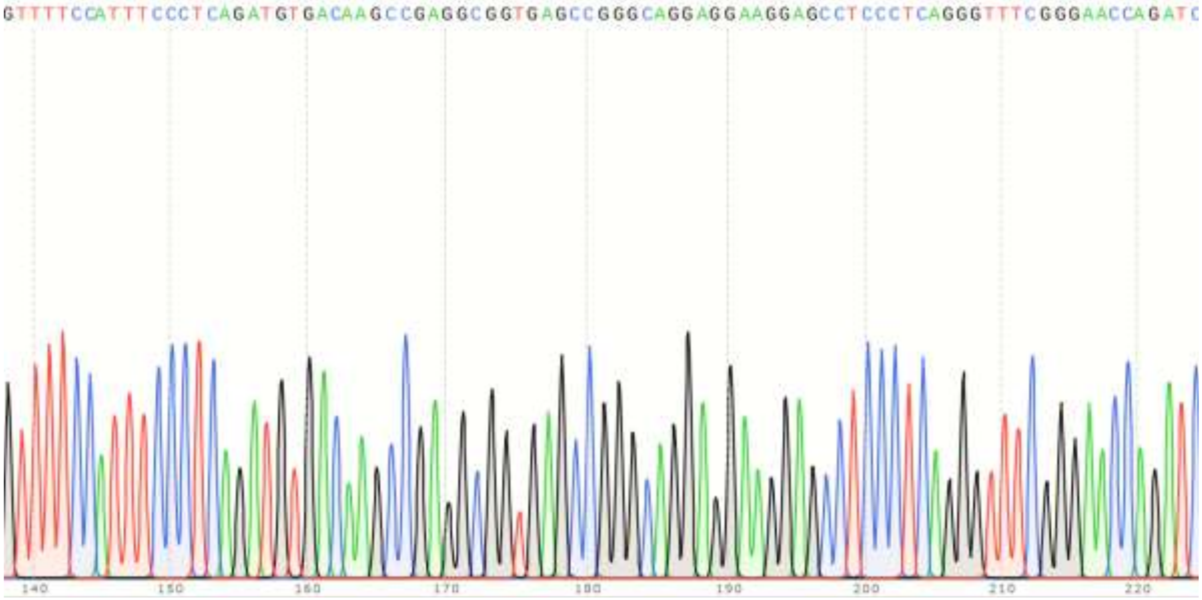


**B**

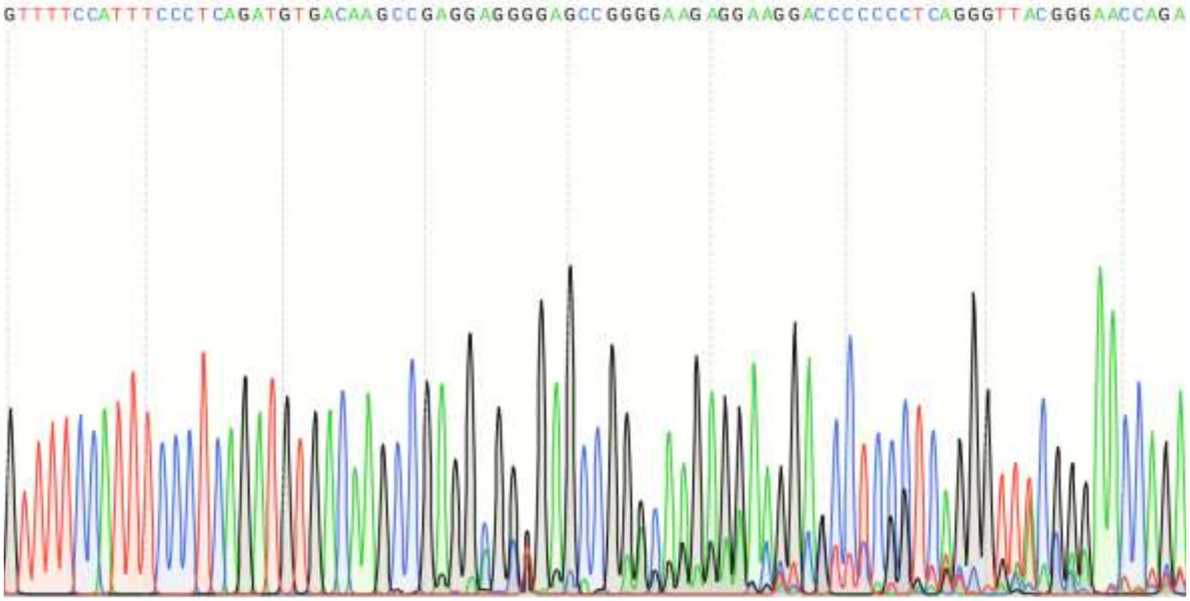


**Figure S17**. Sanger sequencing of indel mutations in VEGFA fragment under different conditions. A) Sequences of VEGFA gene without delivered *VEFGA*-crRNA with UV light (365 nm, 10 mW·cm$^{-2}$). B) Sequences of VEGFA gene with delivered *VEGFA*-crRNA with UV light (365 nm, 10 mW·cm$^{-2}$).

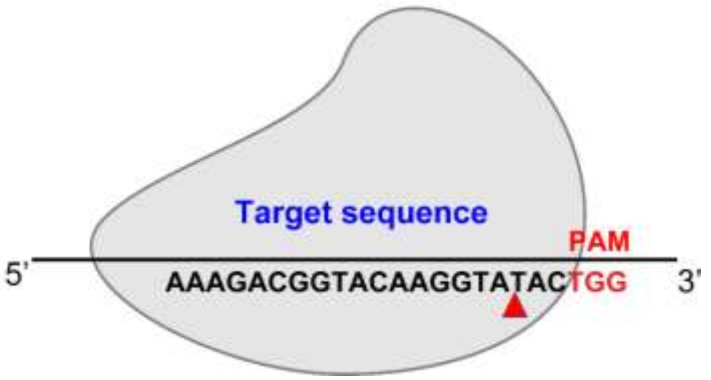


| | |
|---|---|
| Wild sequence | AACCCATGAAAGACGGTACAAGGTATACTGGAATCCGATCTCTGAAA |
| Typical types of mutated sequence | AACCCATGAAAGACGGTACAAGG- - - - - - - - - - AATCCGATCTCTGAAA |
| | AACCCATGAAAGACGGTAC - - - - - - - - - - TGGAATCCGATCTCTGAAA |
| | AACCCATGAAAGACGGTACAAGGTATTACTGGAATCCGATCTCTGAAA |
| | AACCCATGAAAGACGGTACAAGGTAT - - - - - - - - - CCGATCTCTGAAA |
| | AACCCATGAAAGACGGTACAAGG- - - - - - - - - - AATCCGATCTCTGAAA |
| | AACCCATGAAAGACGGTACAAGGTAT - - - - - - - - - CCGATCTCTGAAA |
| | AACCCATGAAAAACGGTACAAGGTA - - CTGGAATCCGATCTCTGAAA |
| | AACCCATGAAAGACGGTACAAGGTATT--GGAAACCCATCTCTGAAA |
| | AACCCATGAAAGACGGTACAAGG--TACTGGAATCCCATCTCTGAAA |
| | AACCCATGAAAGACGGTACAAGGAATTACTGGAATCCGATCTCCGAAA |
| | AACCCATGAAAAAAGGGACAAGGGAA--TGGAATCCCATCTCTGAAA |
| | AACCCATGAAAGACGGTACAAGGTA - - CTGGAATCCGATCTCTGAAA |
| | AACCCCTGGAAGACGGGACAAGG- ATACTGGAATCCCAACTCTGAAA |
| | AACCCATG AAAGACGGTACAAGGTA- - CTGGAATCCCATCTCTGAAA |
| | AACCCATGAAAGACGGTACAAGGTATTGTGGAATCCCATCTCTGAAA |

**Figure S18**. Sequences of MSTN gene with co-delivered unmodified or circular sgRNAs and Cas9 protein induced mutations. The wild-type MSTN gene sequences were shown on the top. Sequences in red were the target sites of guide RNA followed by PAM sequence; Dashes indicated the deletion of nucleotides, and Blue indicated insertion of nucleotides. Same types of mutation were not given.

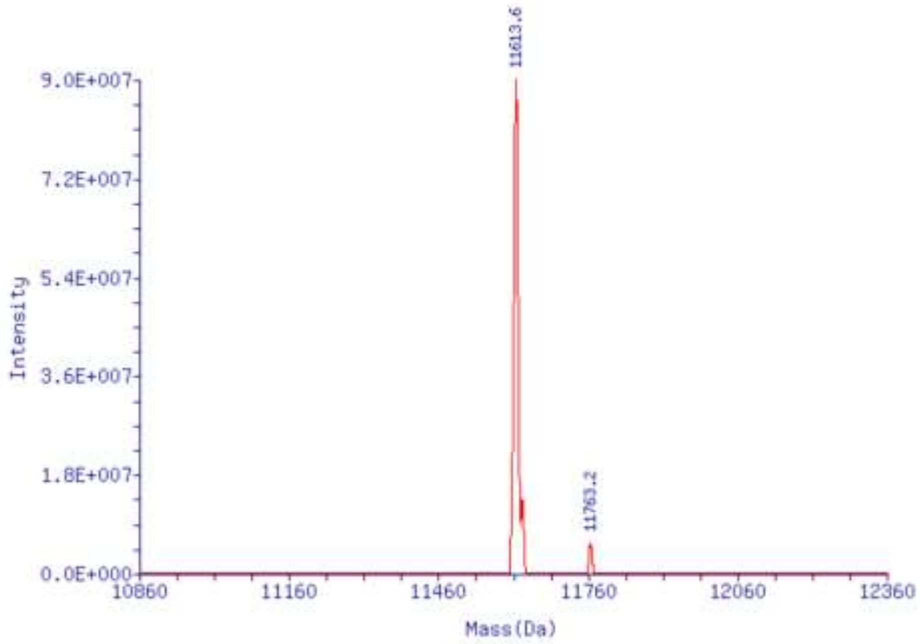


**Figure S19**. ESI Mass Spectrum of *GFP*-crRNA

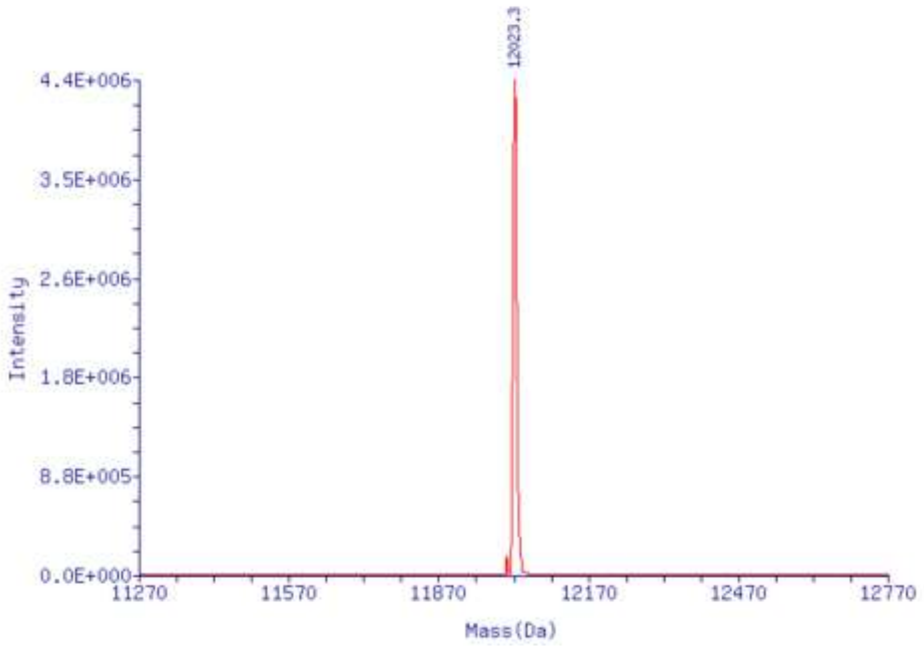


**Figure S20**. ESI Mass Spectrum of *GFP*-crRNA-a1

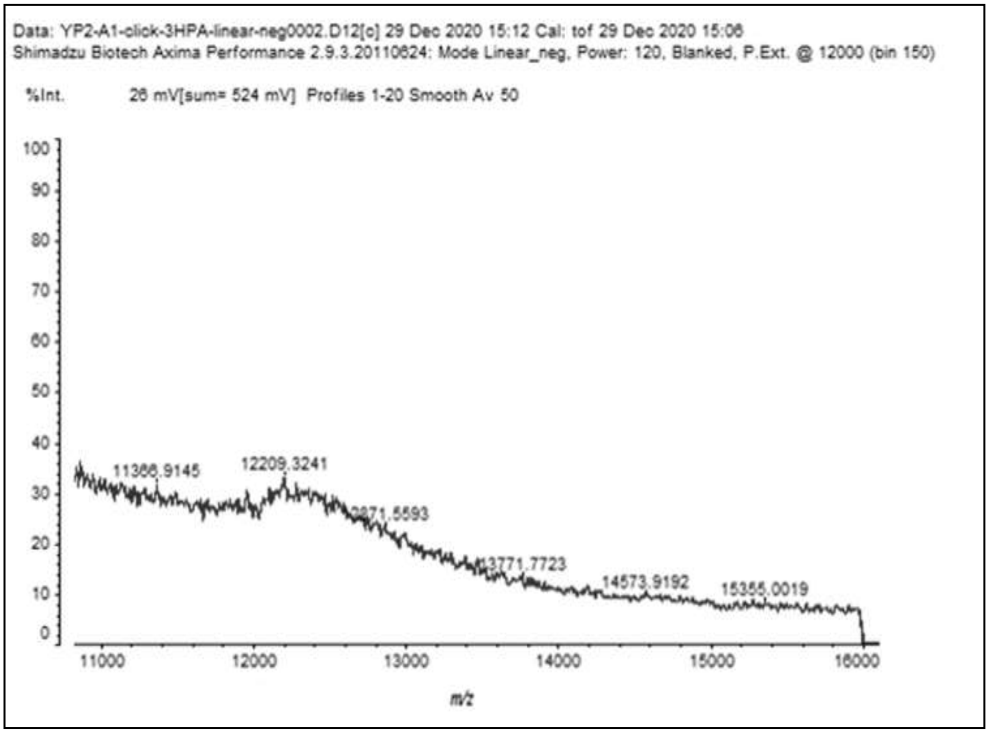


**Figure S21**. MALDI-TOF of *GFP*-crRNA-**c1**

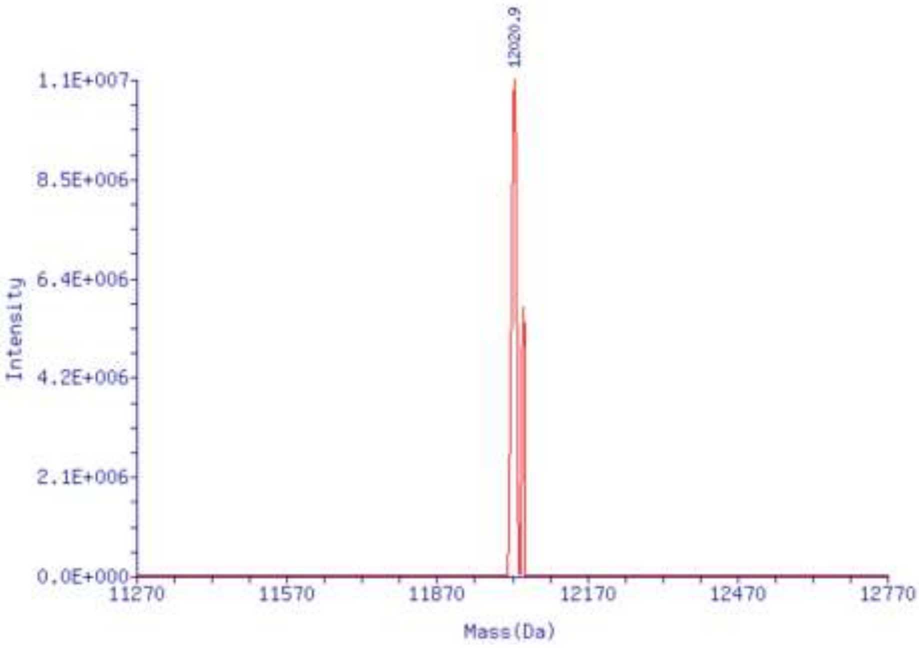


**Figure S22**. ESI Mass Spectrum of *GFP*-crRNA-a2

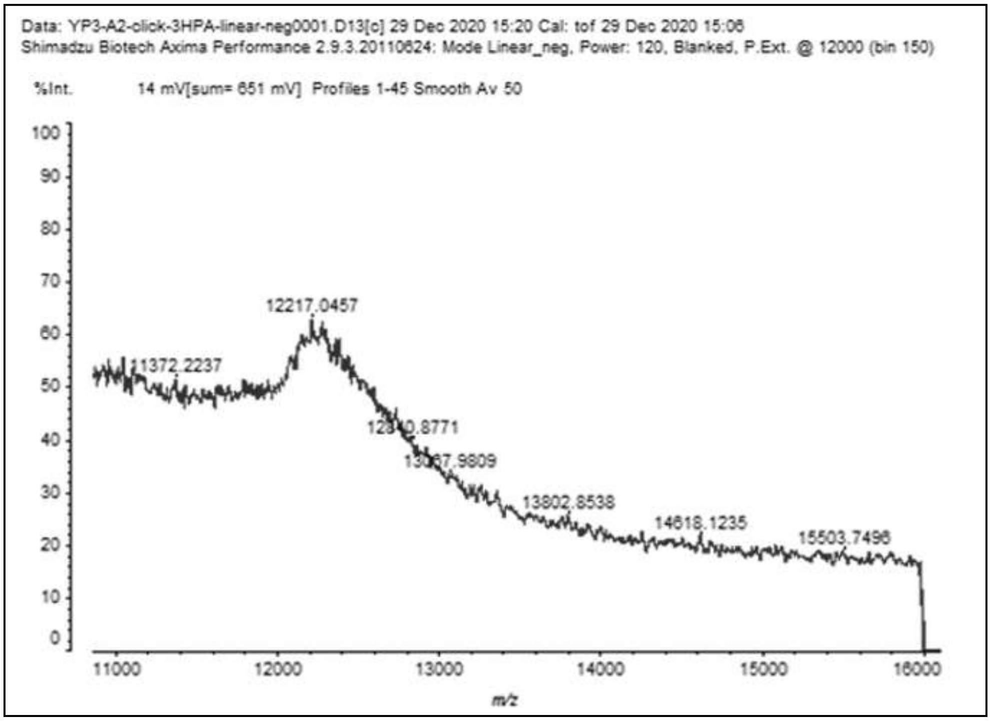


**Figure S23**. MALDI-TOF of *GFP*-crRNA-**c2**

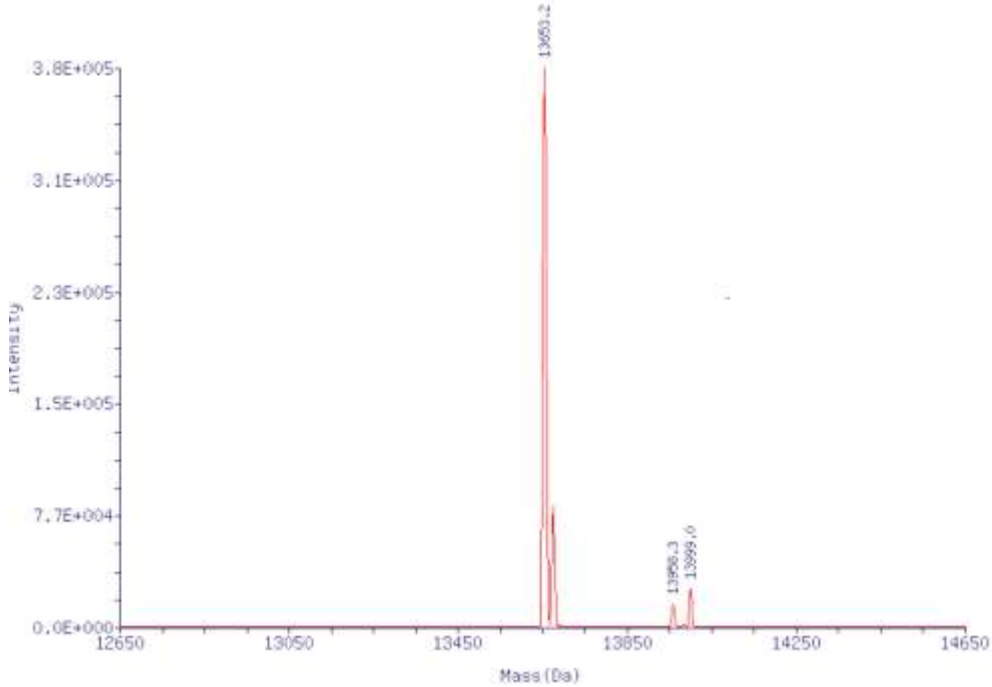


**Figure S24**. ESI Mass Spectrum of *GFP*-cpf1-**sgRNA**

**[TBF]**

**Figure S25**. MALDI-TOF of *GFP*-Cpf1-sgRNA-**a1**

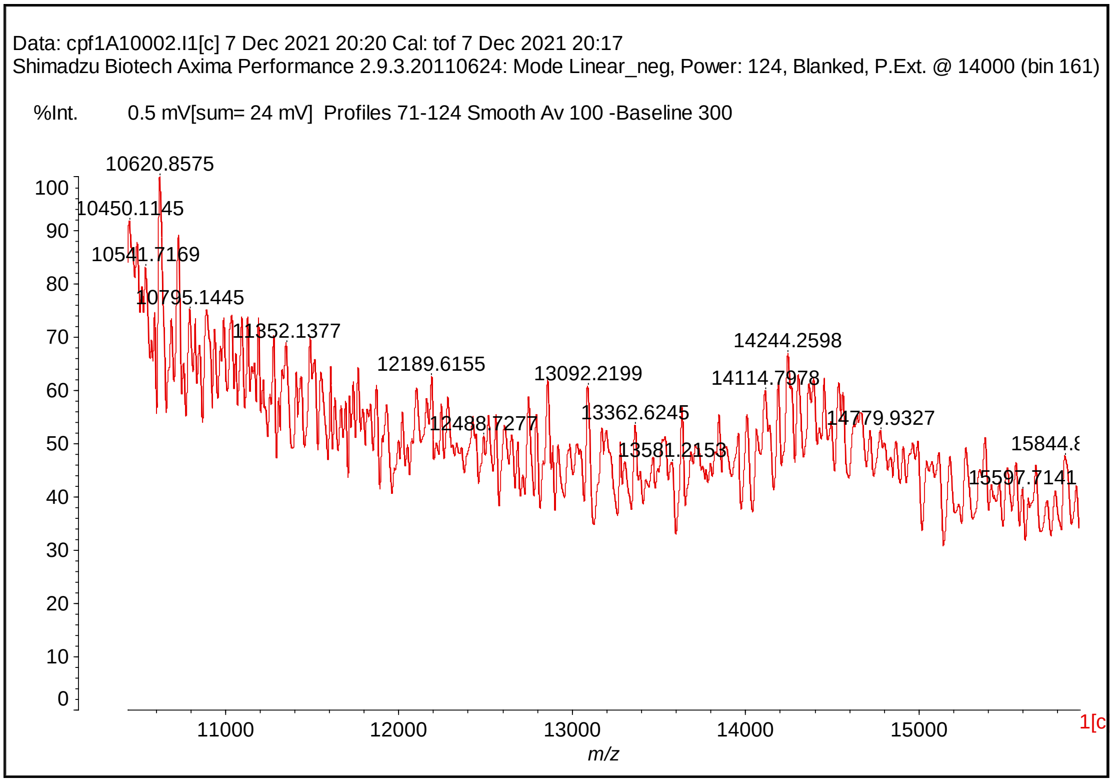


**Figure S26**. MALDI-TOF of *GFP*-Cpf1-sgRNA-**c1**

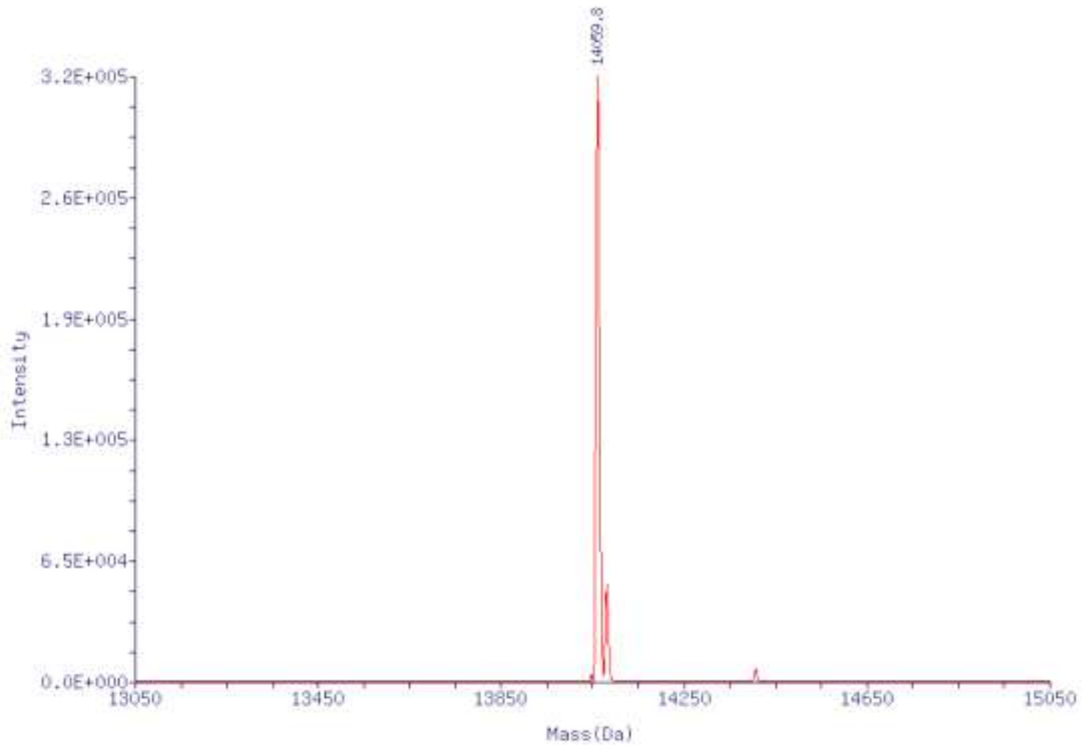


**Figure S27**. ESI Mass Spectrum of *GFP*-cpf1-sgRNA**-a2**

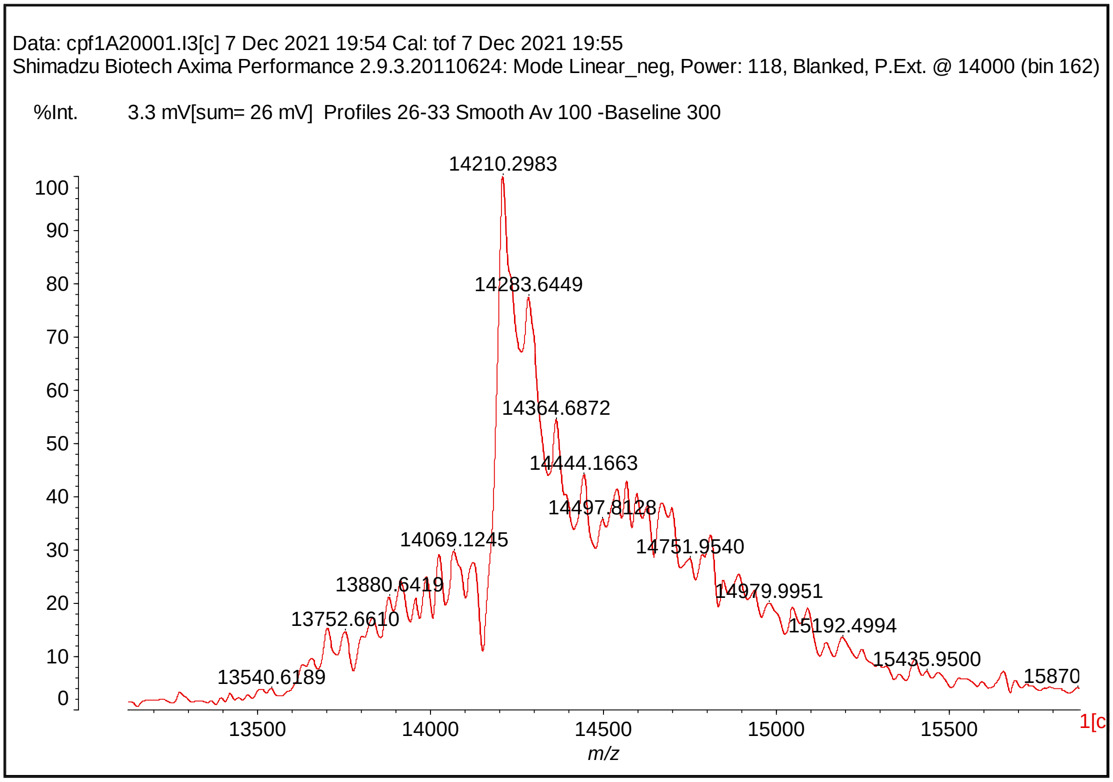


**Figure S28**. MALDI-TOF of *GFP*-Cpf1-sgRNA-**c2**

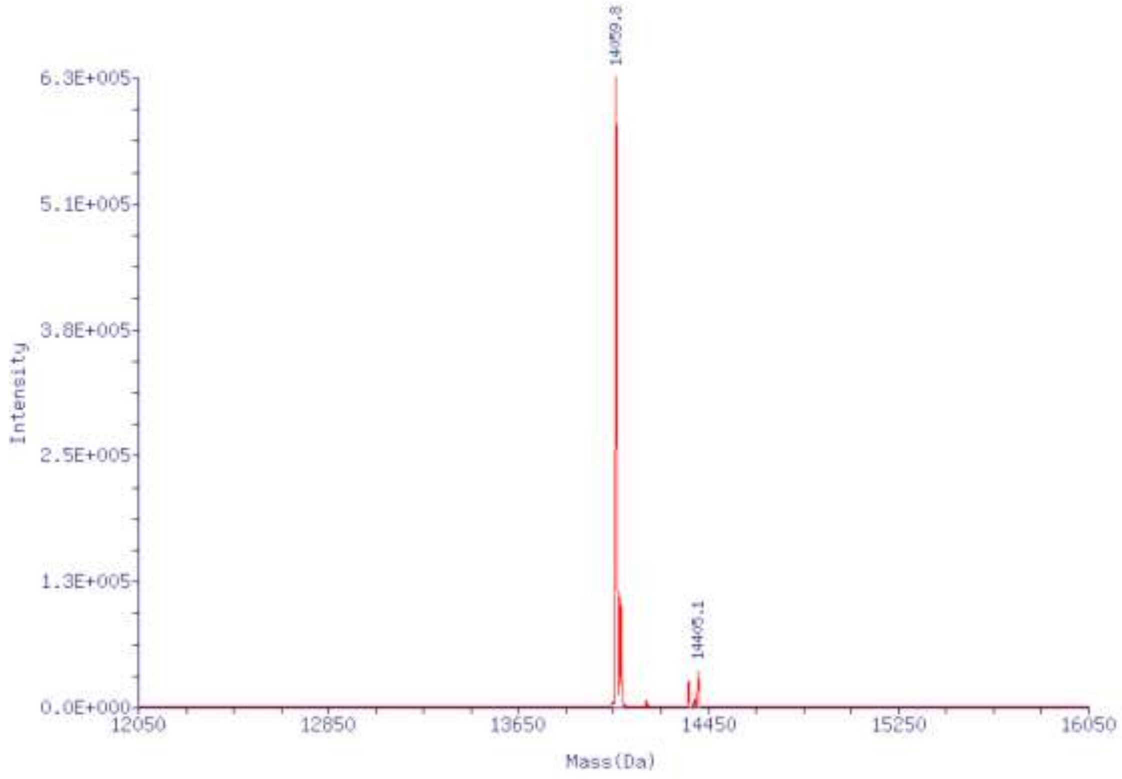


**Figure S29**. ESI Mass Spectrum of *GFP*-cpf1-sgRNA**-a3**

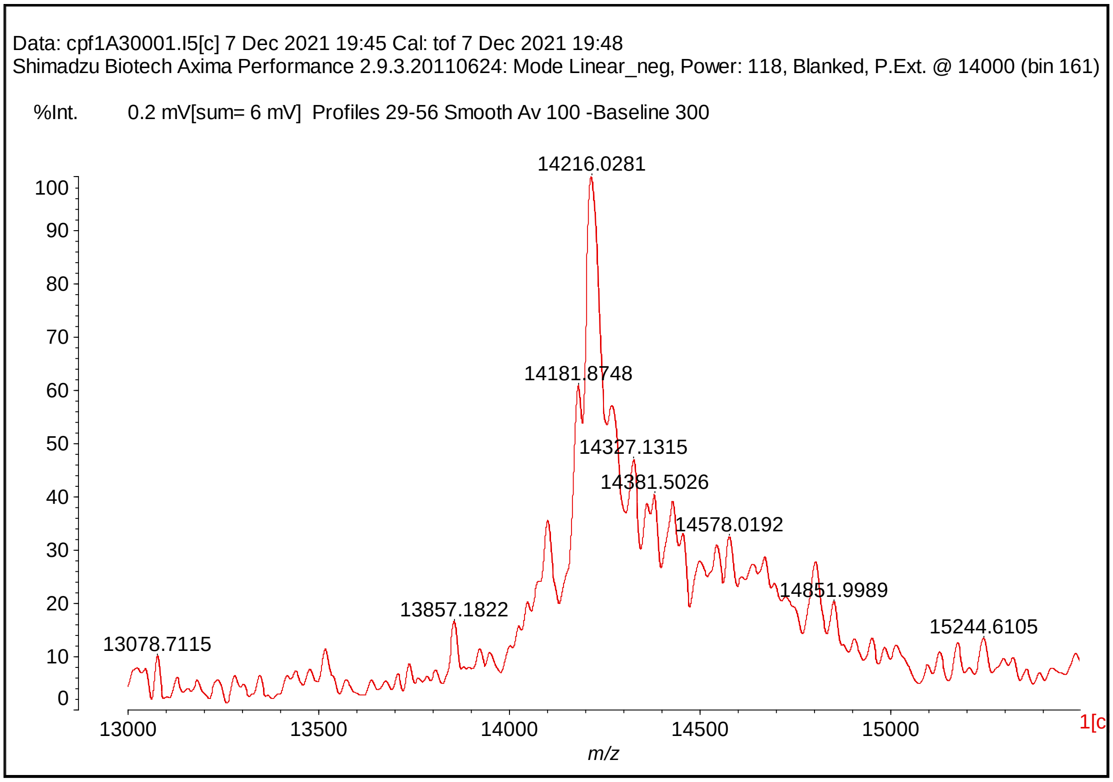


**Figure S30**. MALDI-TOF of *GFP*-Cpf1-sgRNA-**c3**

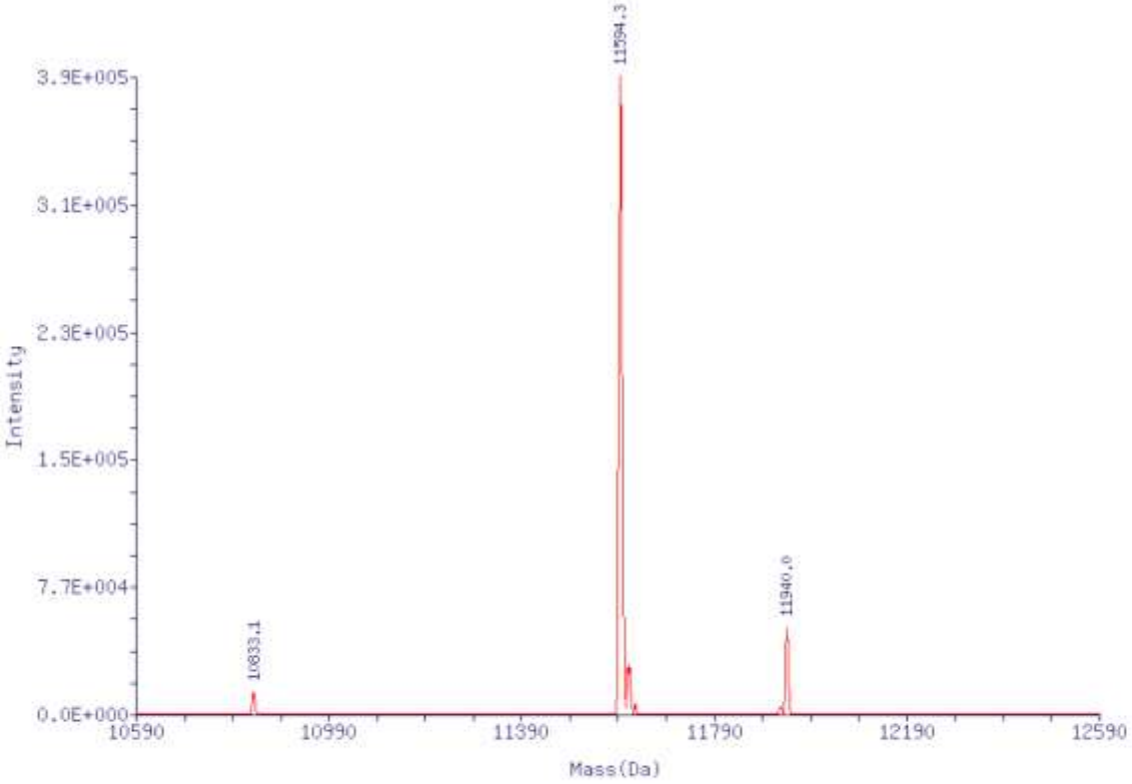


**Figure S31**. ESI Mass Spectrum of *VEGFA*-crRNA

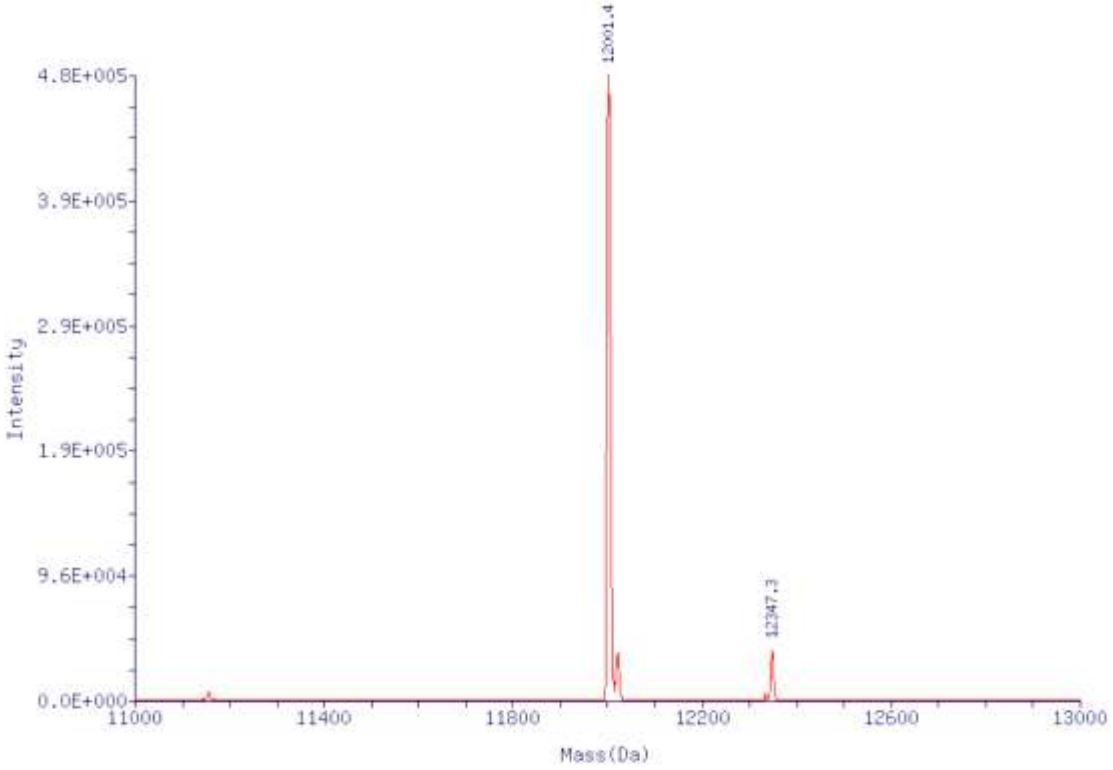


**Figure S32**. ESI Mass Spectrum of *VEGFA*-crRNA-**a**

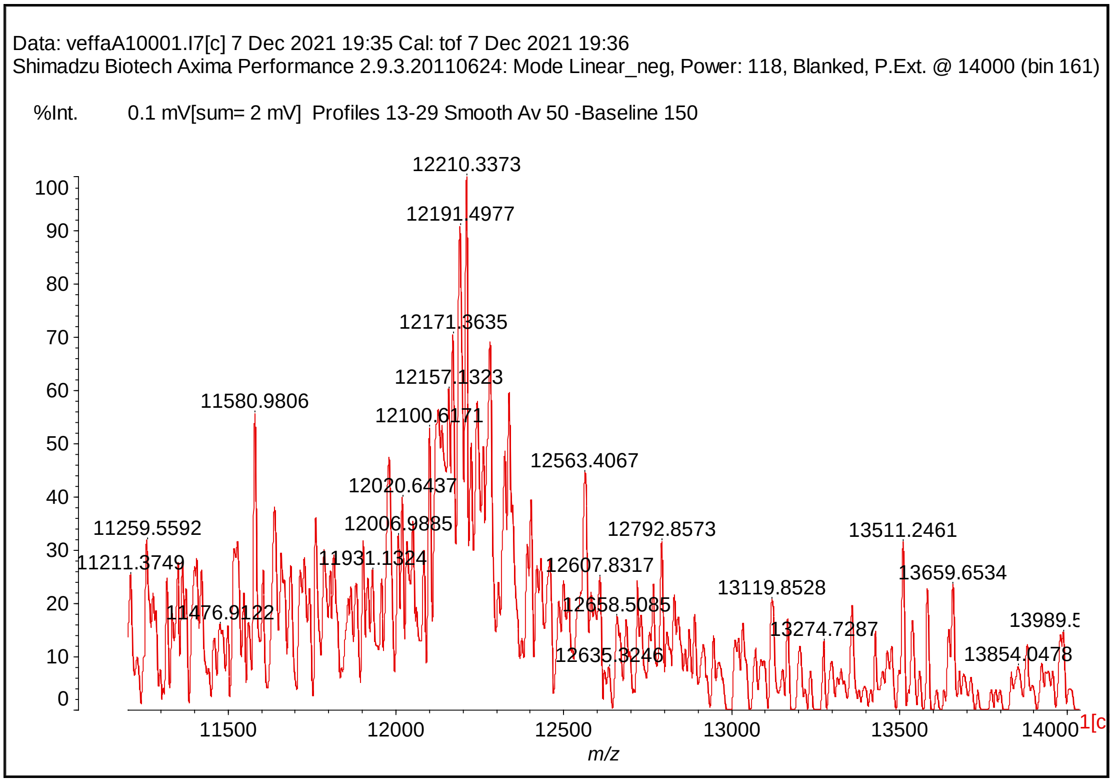


**Figure S33**. MALDI-TOF of *VEGFA*-crRNA-**c**

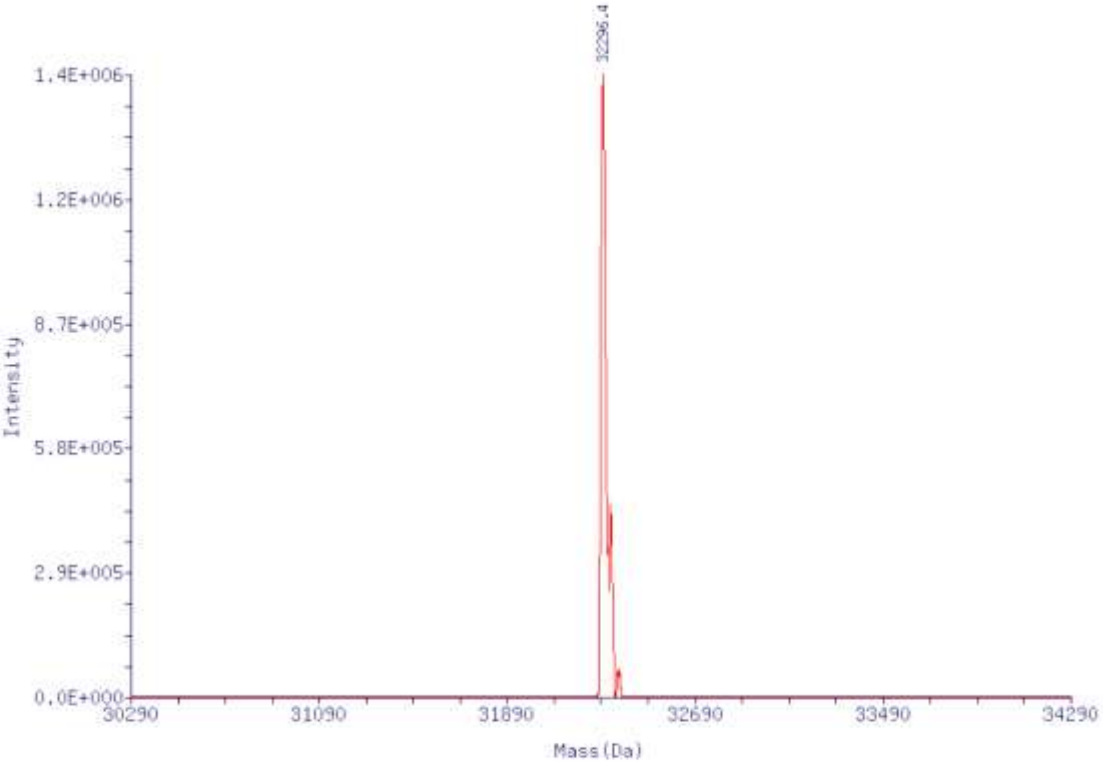


**Figure S34**. ESI Mass Spectrum of *MSTN*-sgRNA

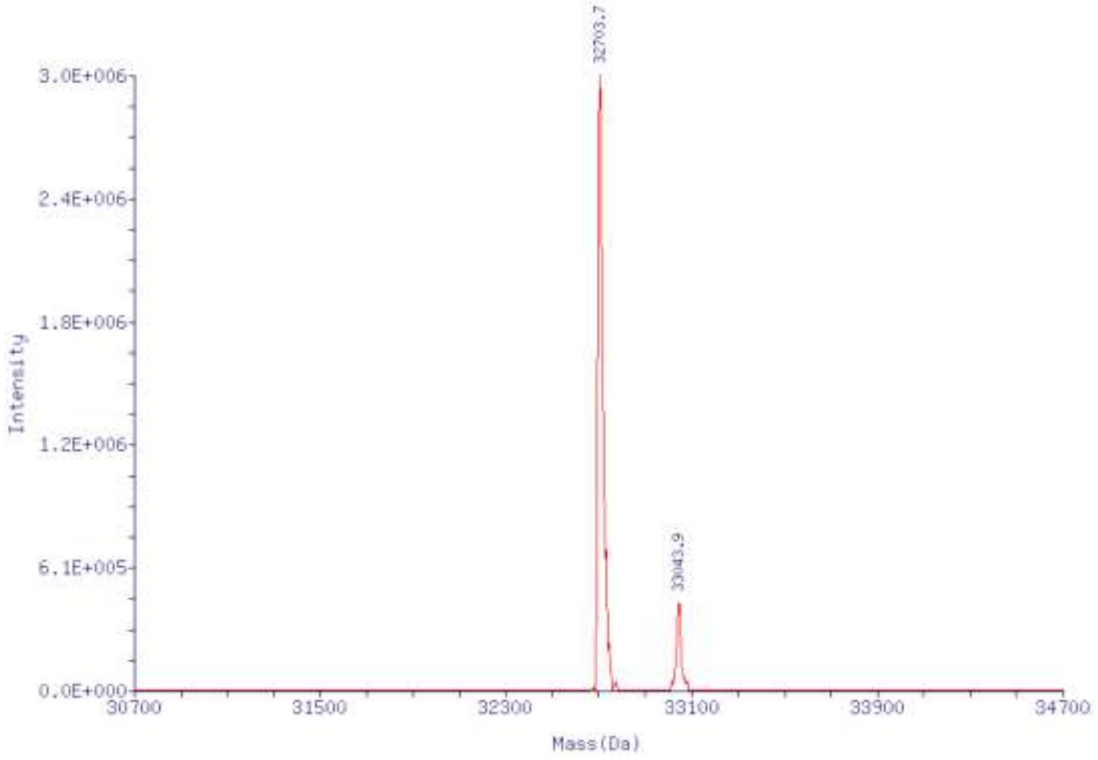


**Figure S35**. ESI Mass Spectrum of *MSTN*-sgRNA-**a1**

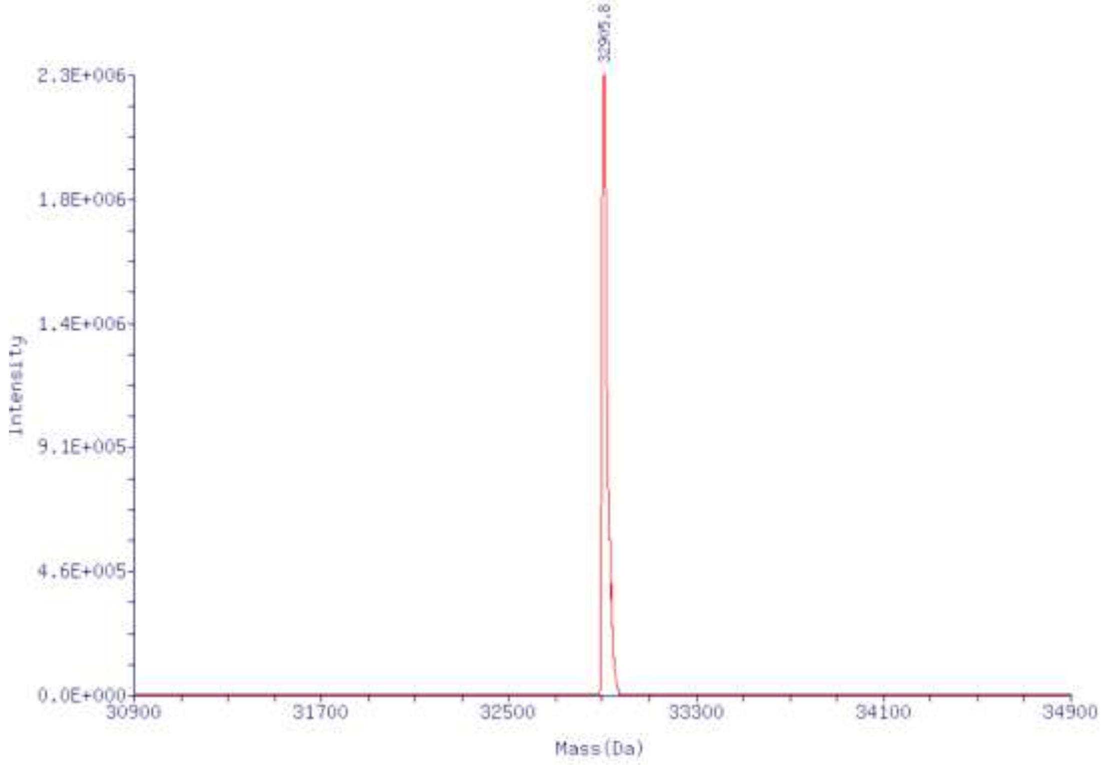


**Figure S36**. ESI Mass Spectrum of *MSTN*-sgRNA-**a2**

#### 14. NMR and mass spectra

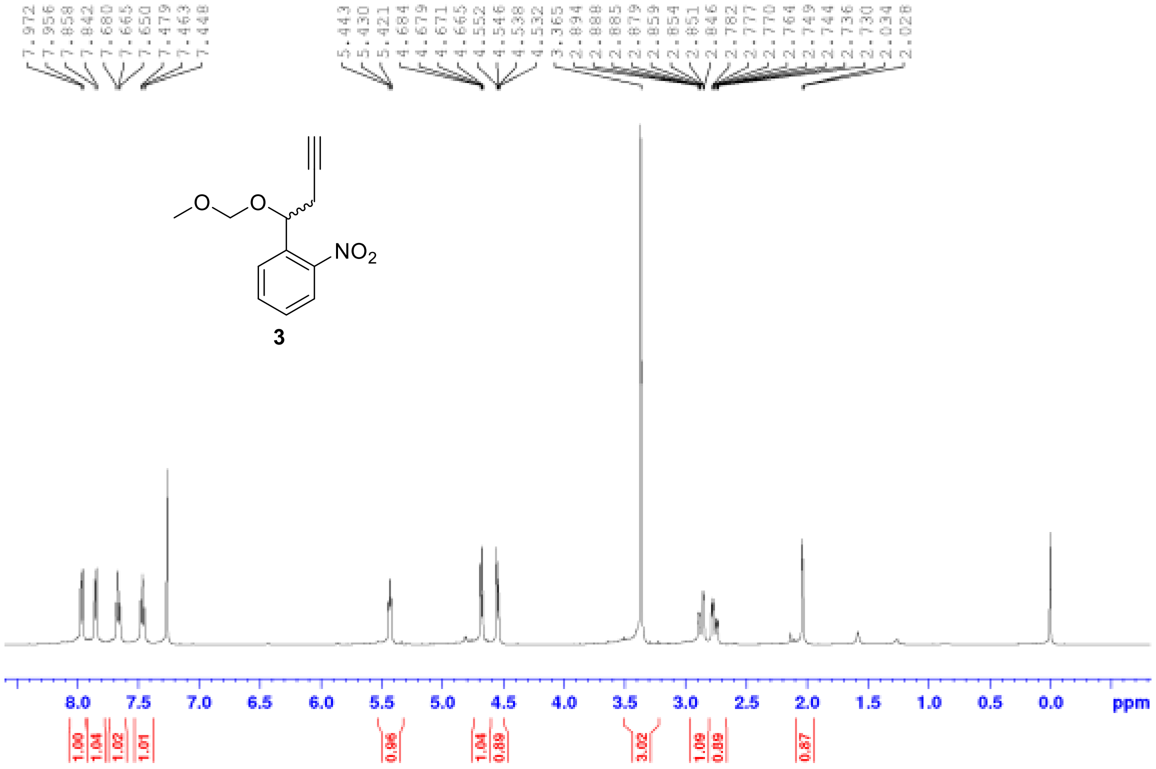

[1]H NMR of compound **3**

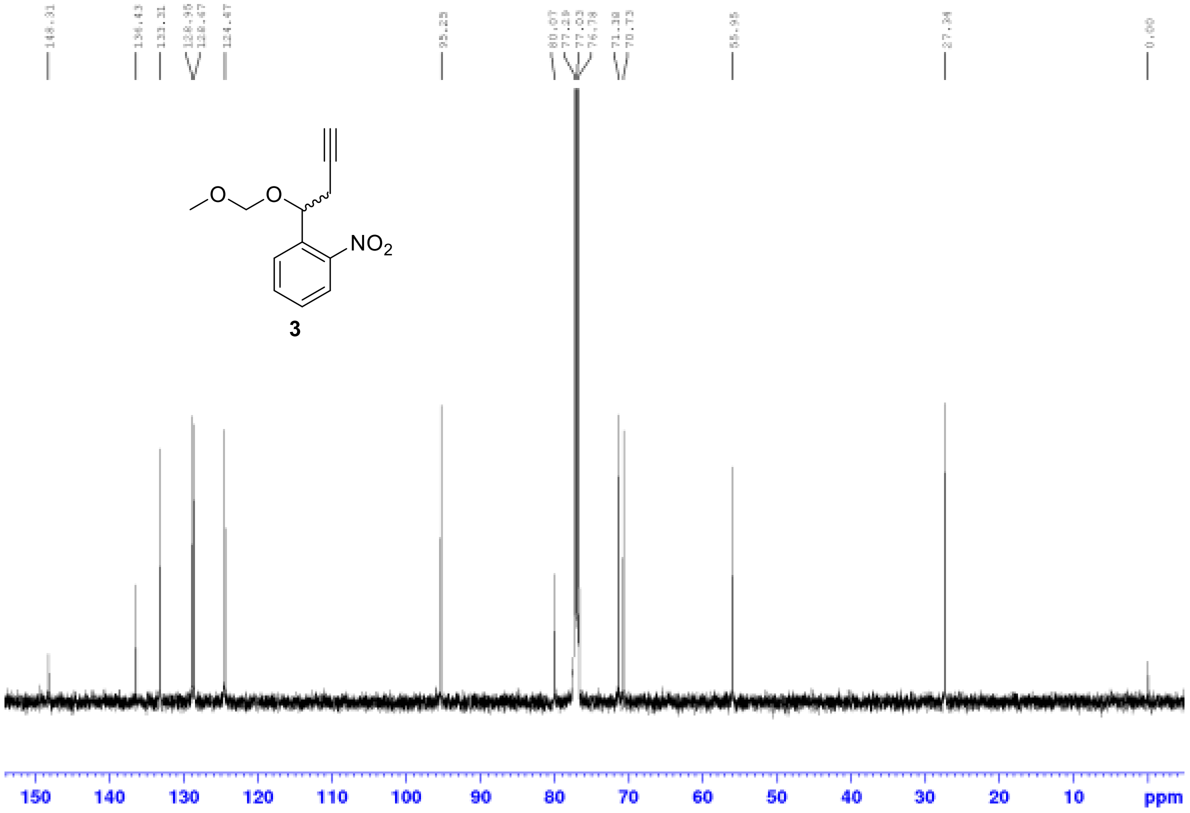

[13]C NMR of compound **3**

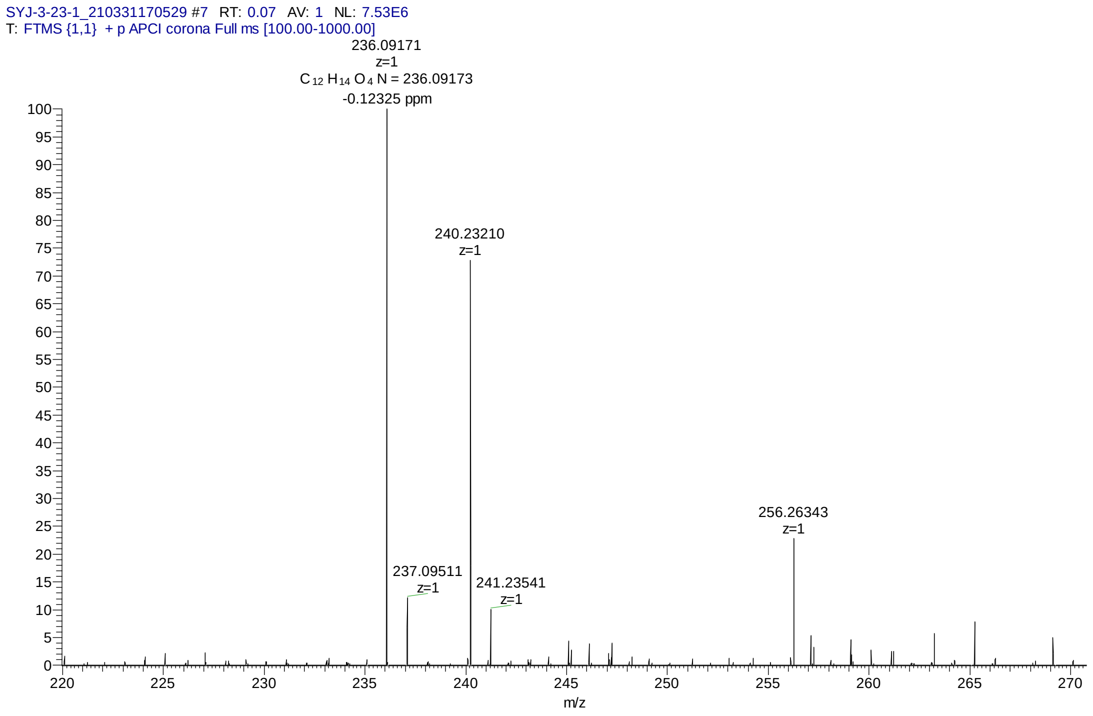


MS spectrum of compound **3**

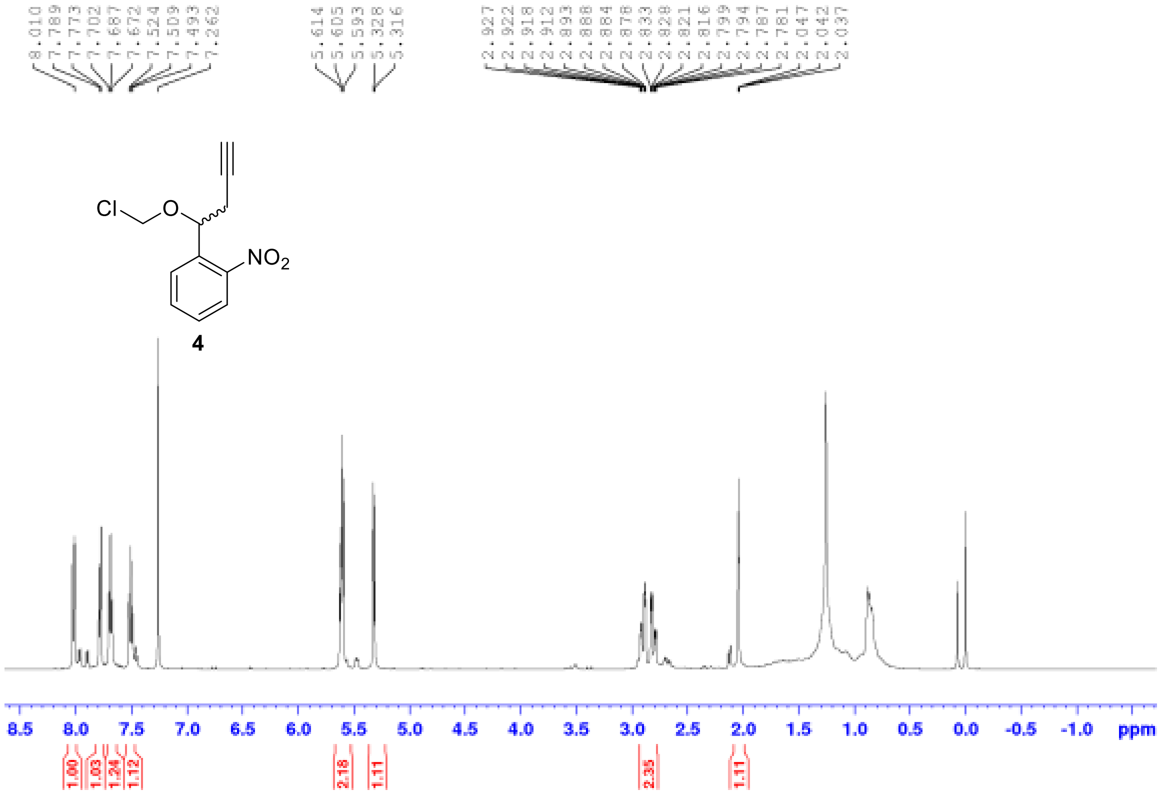


[1]H NMR of compound **4**

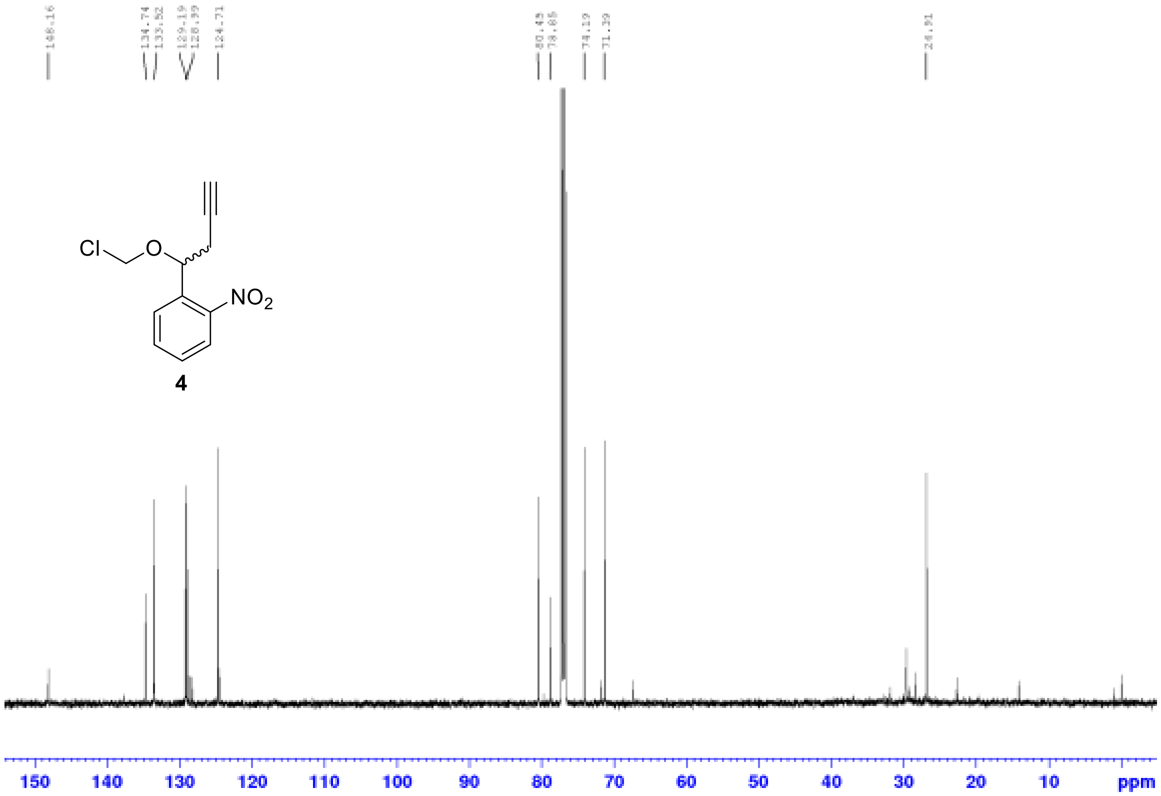

$^{13}$C NMR of compound **4**

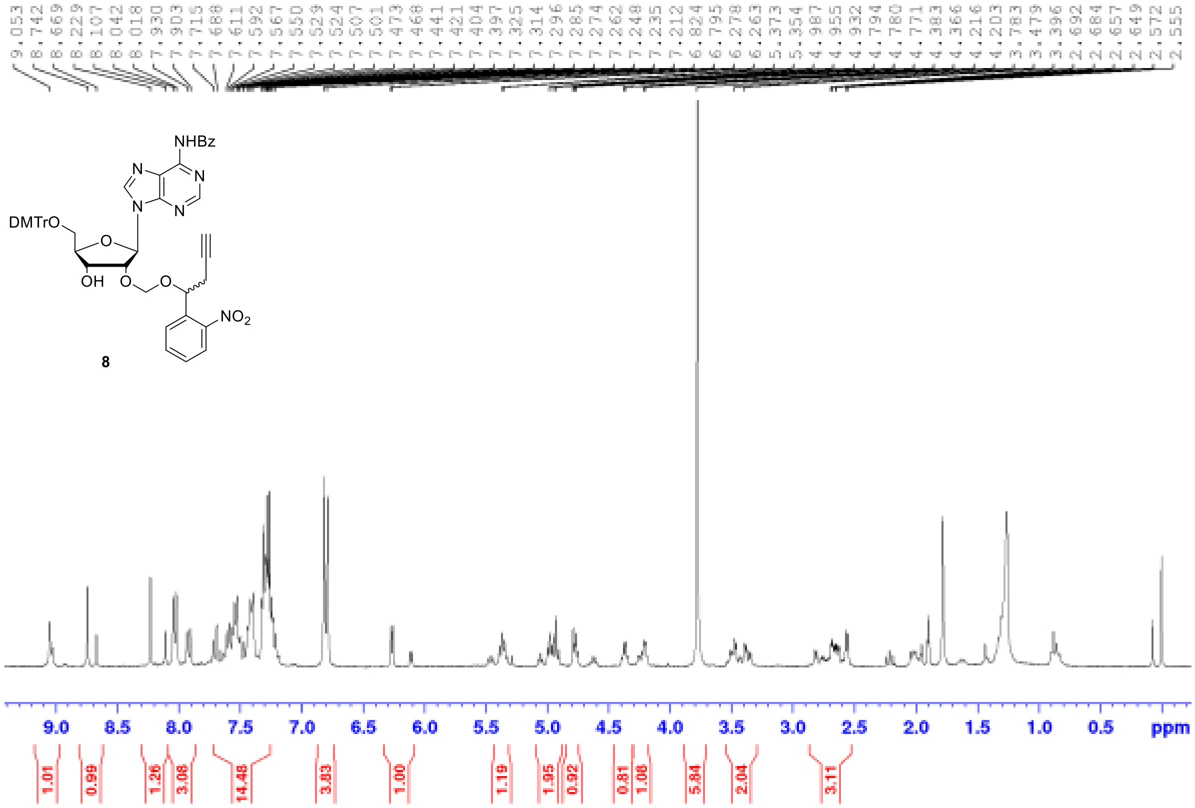

$^{1}$H NMR of compound **8**

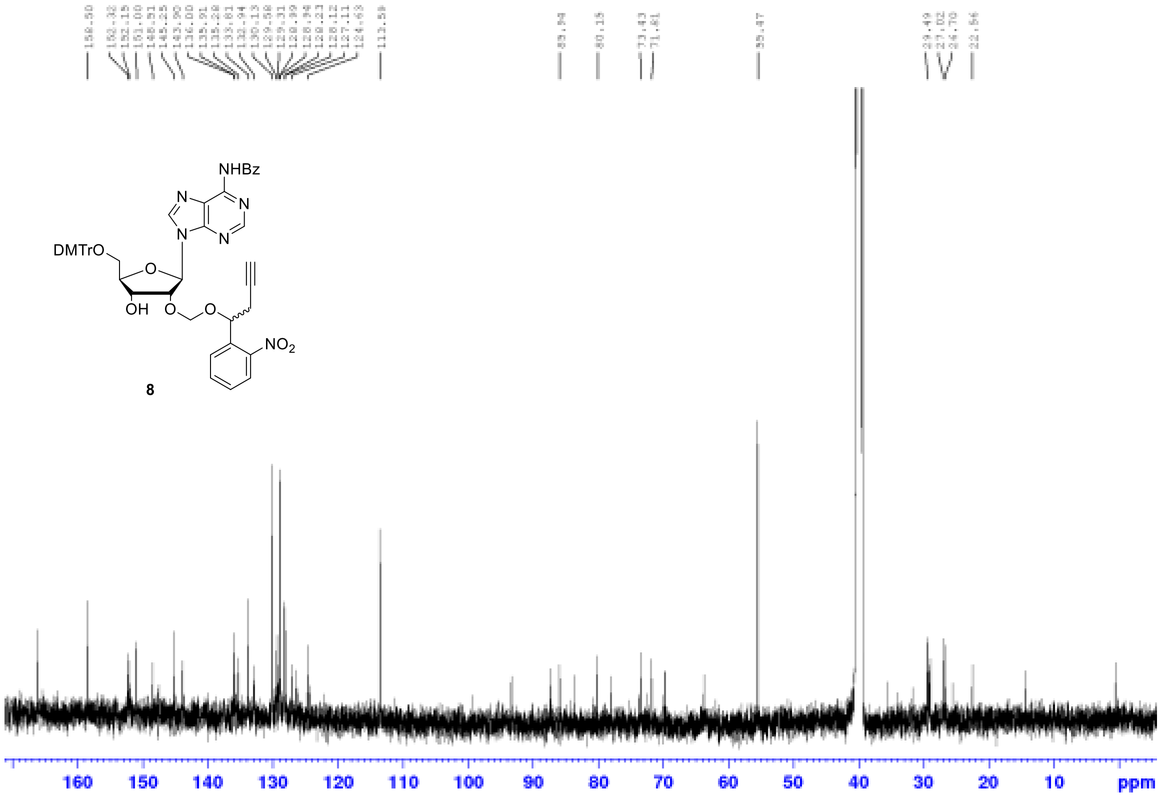


$^{13}$C NMR of compound **8**

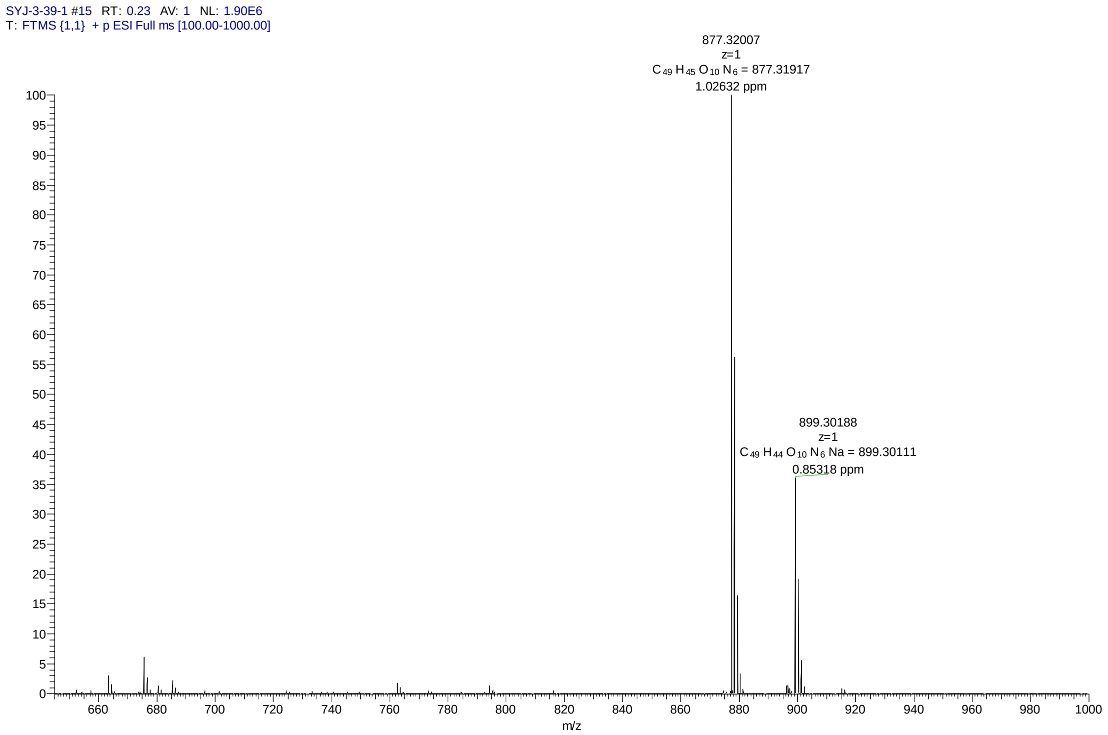


MS spectrum of compound **8**

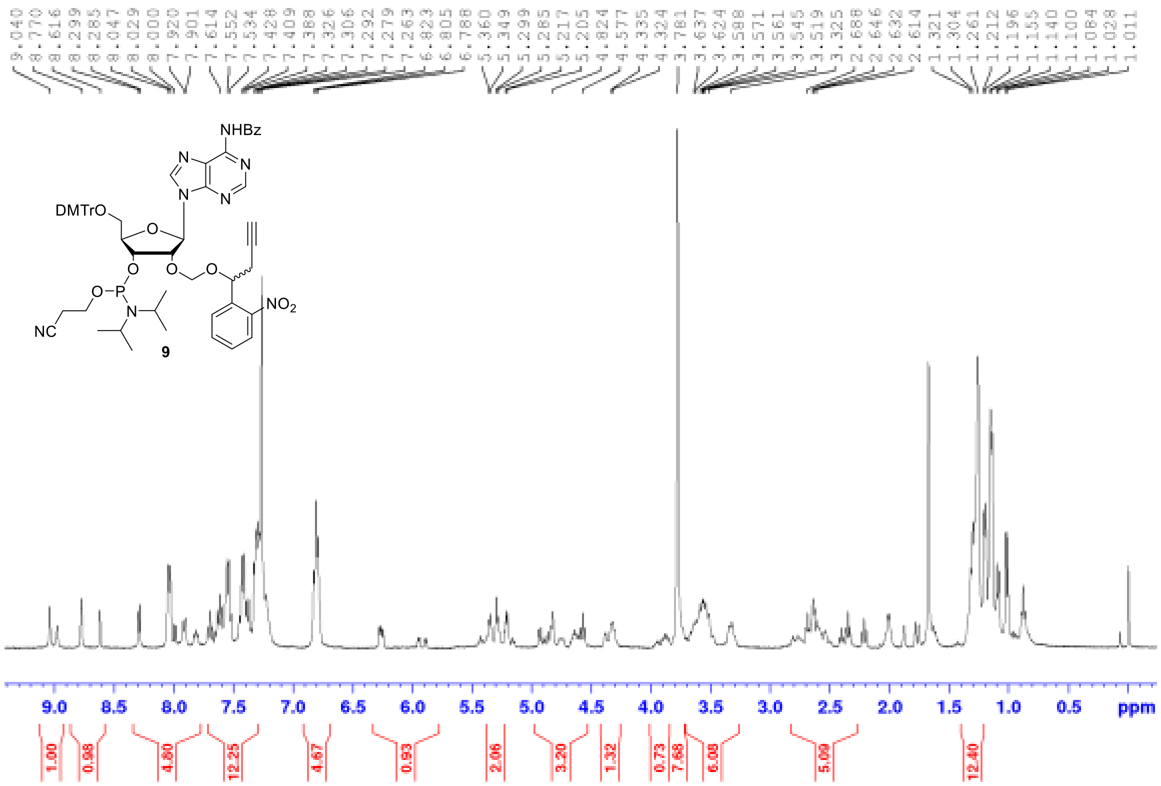

$^1$H NMR of compound **9**

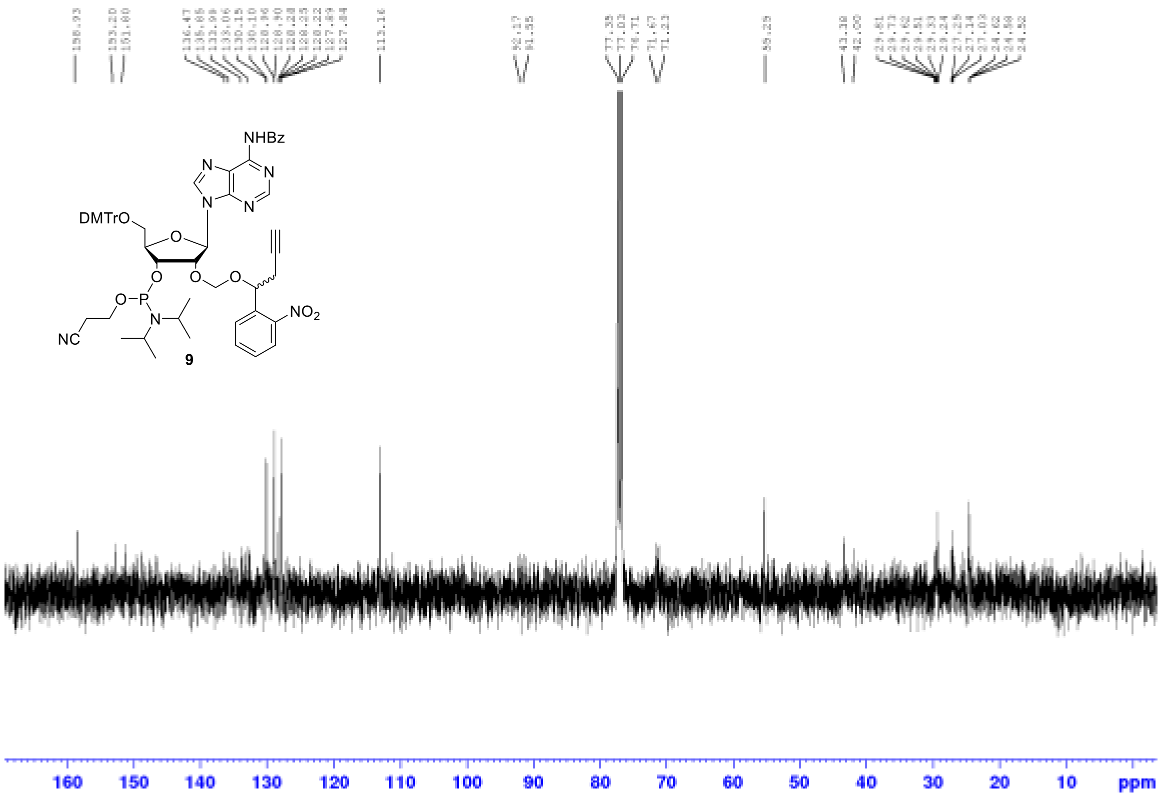

$^{13}$C NMR of compound **9**

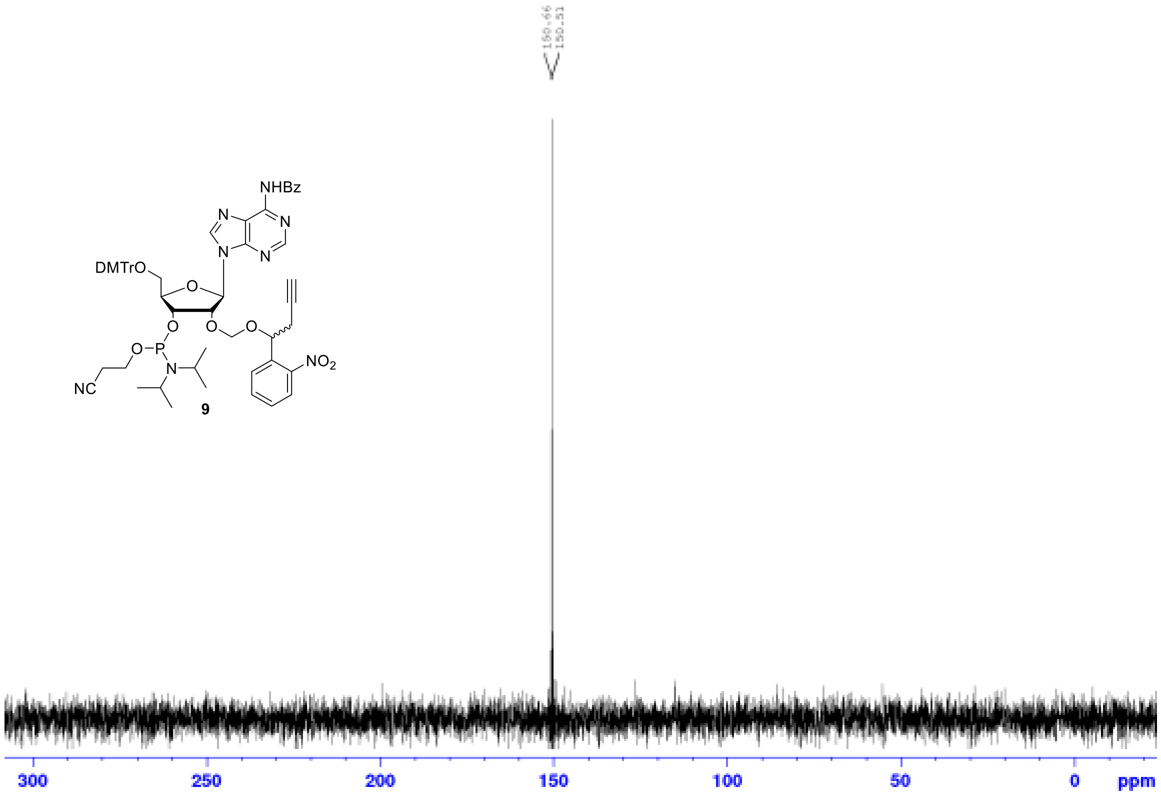


$^{31}$P NMR of compound **9**

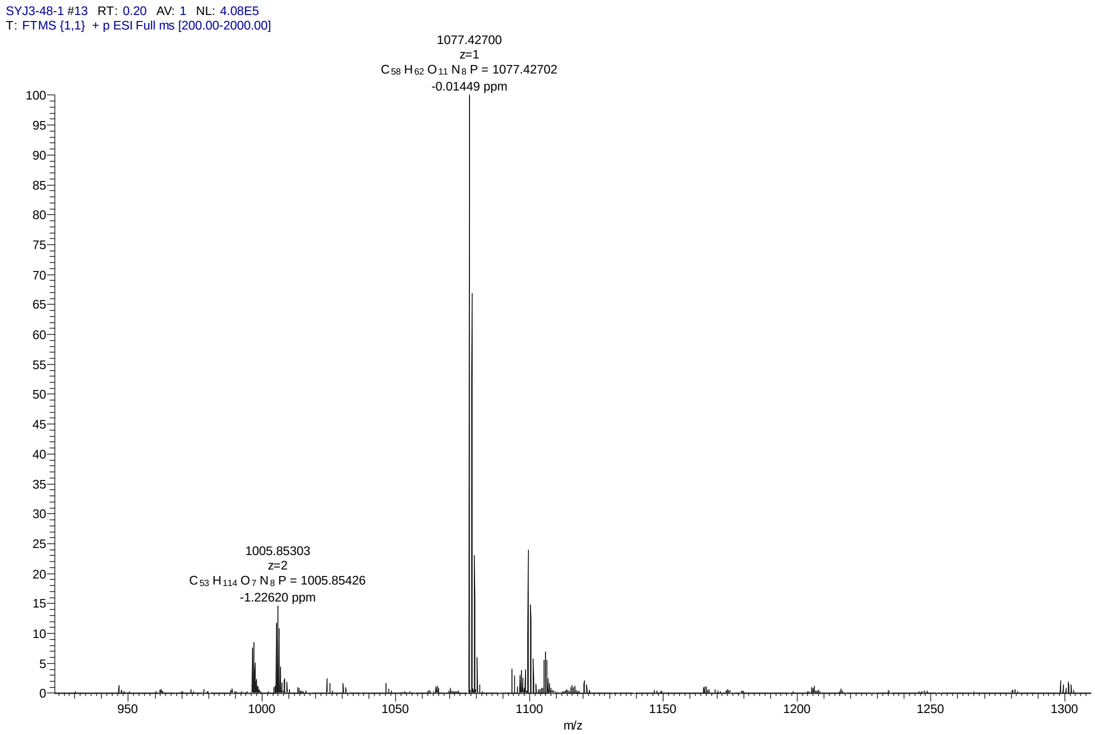


MS spectrum of compound **9**

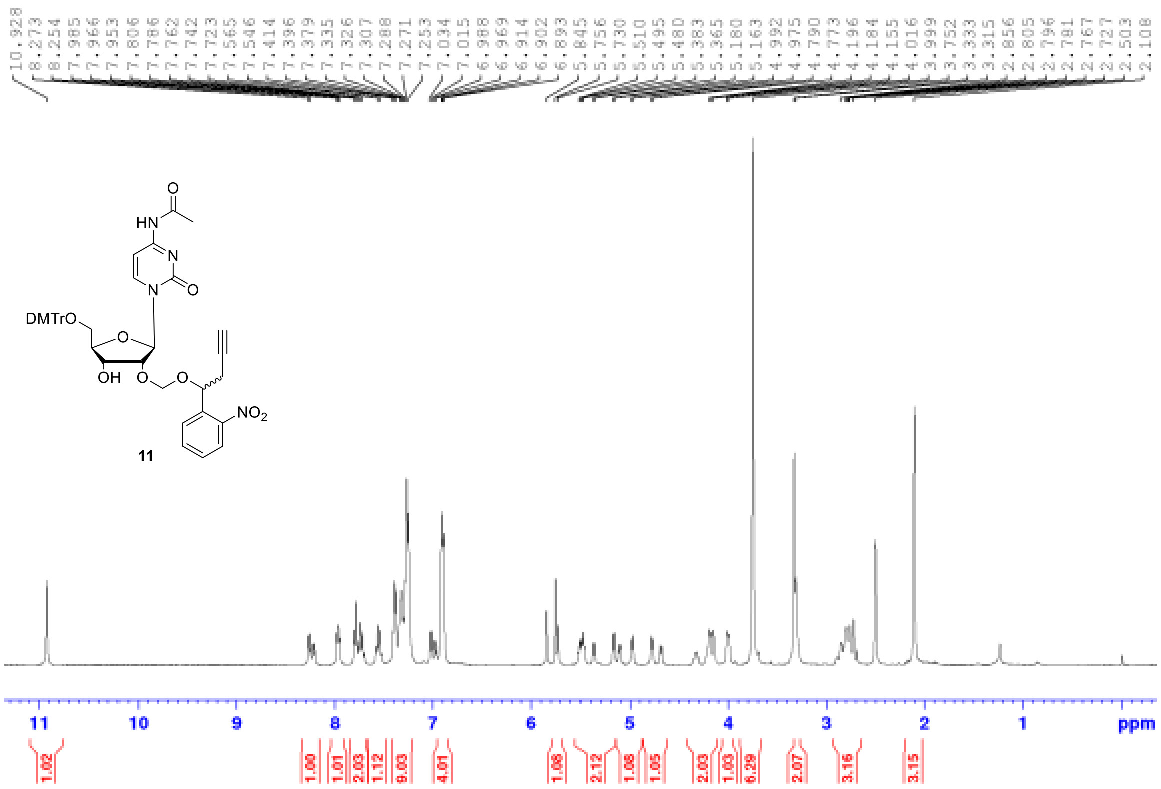

$^1$H NMR of compound **11**

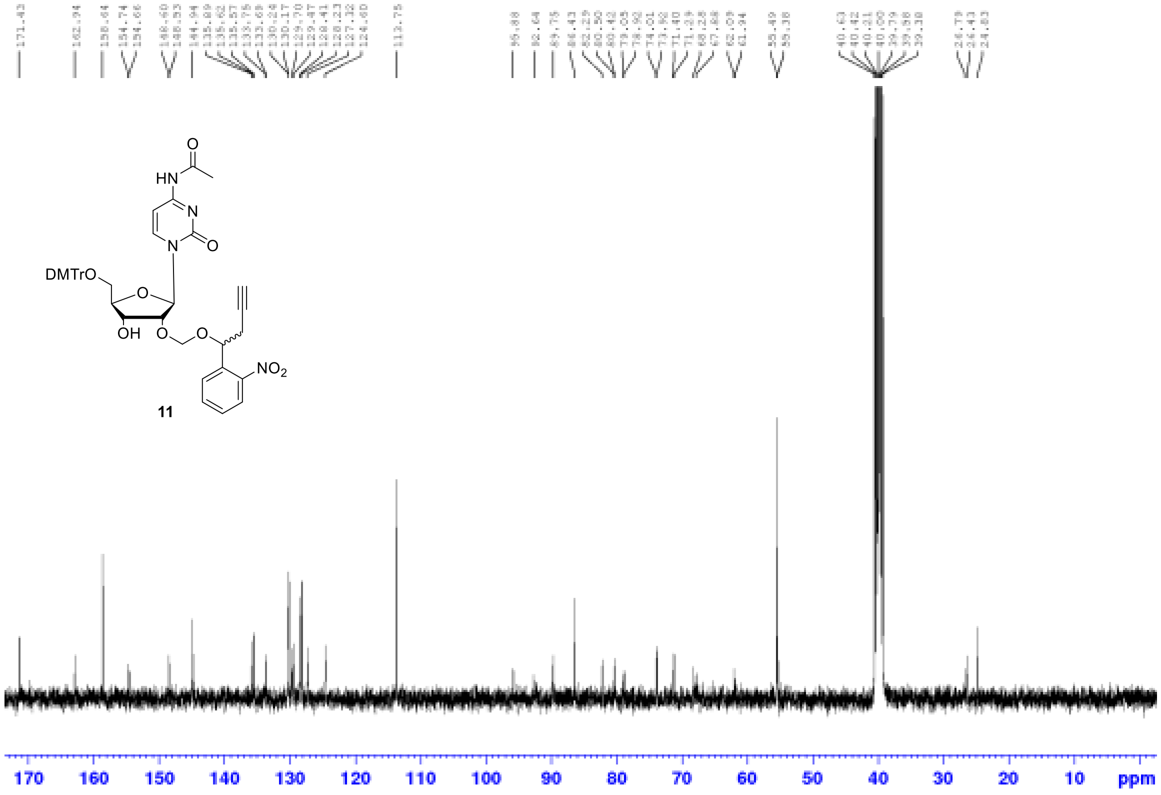

$^{13}$C NMR of compound **11**

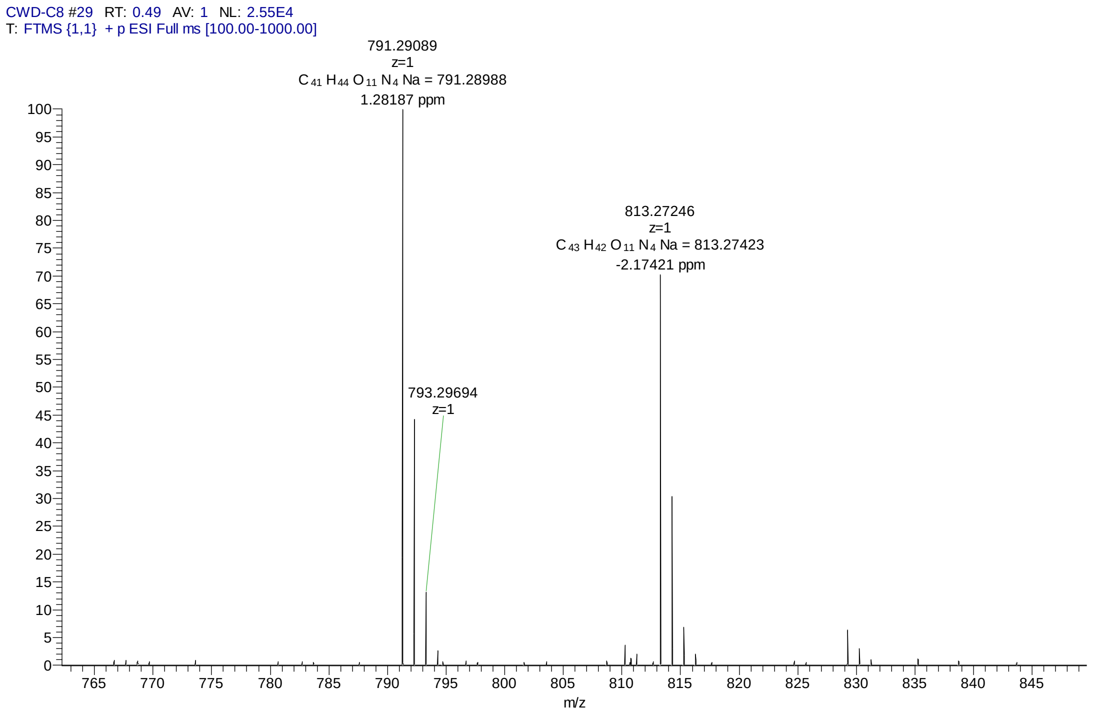


MS spectrum of compound **11**

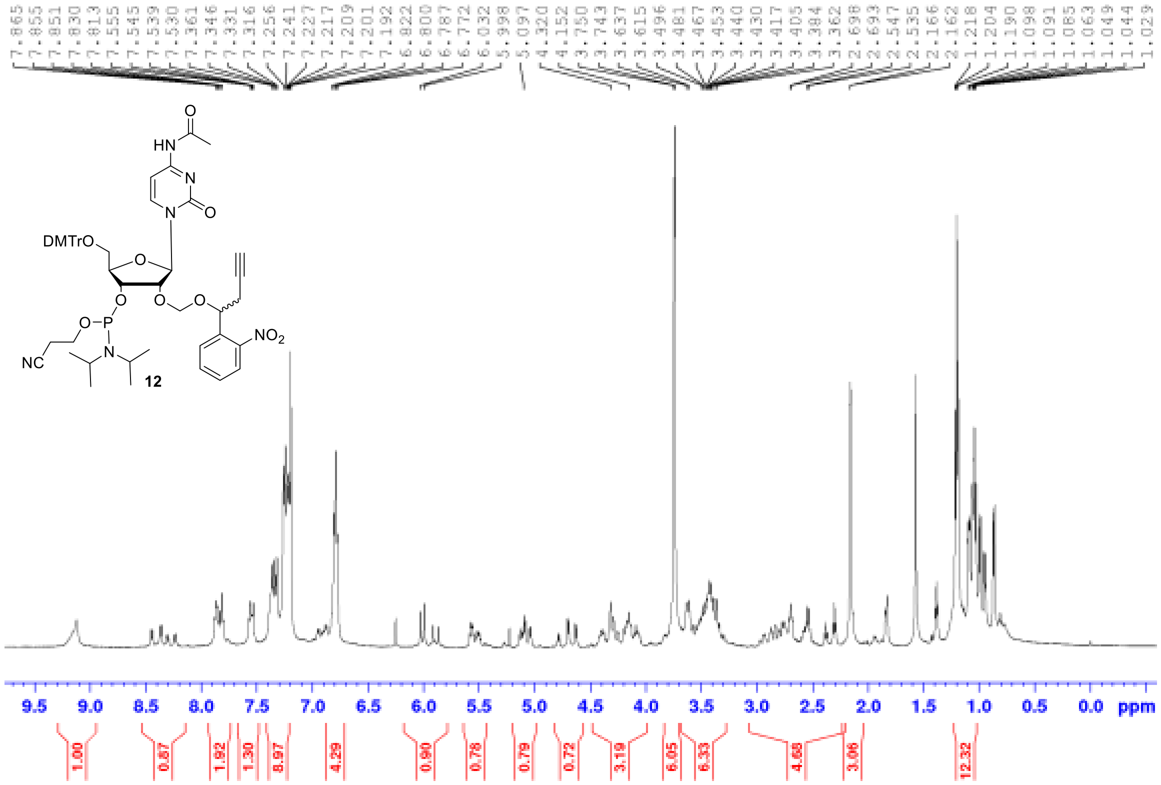


$^1$H NMR of compound **12**

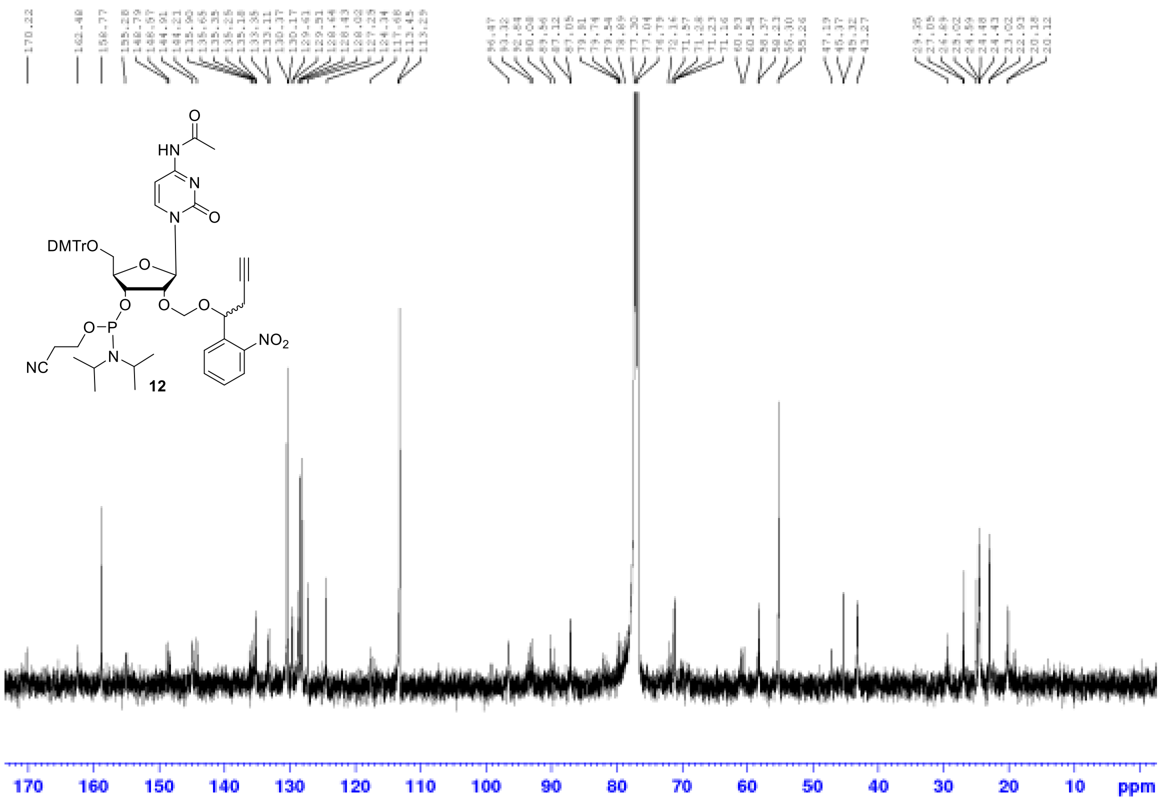

$^{13}C$ NMR of compound **12**

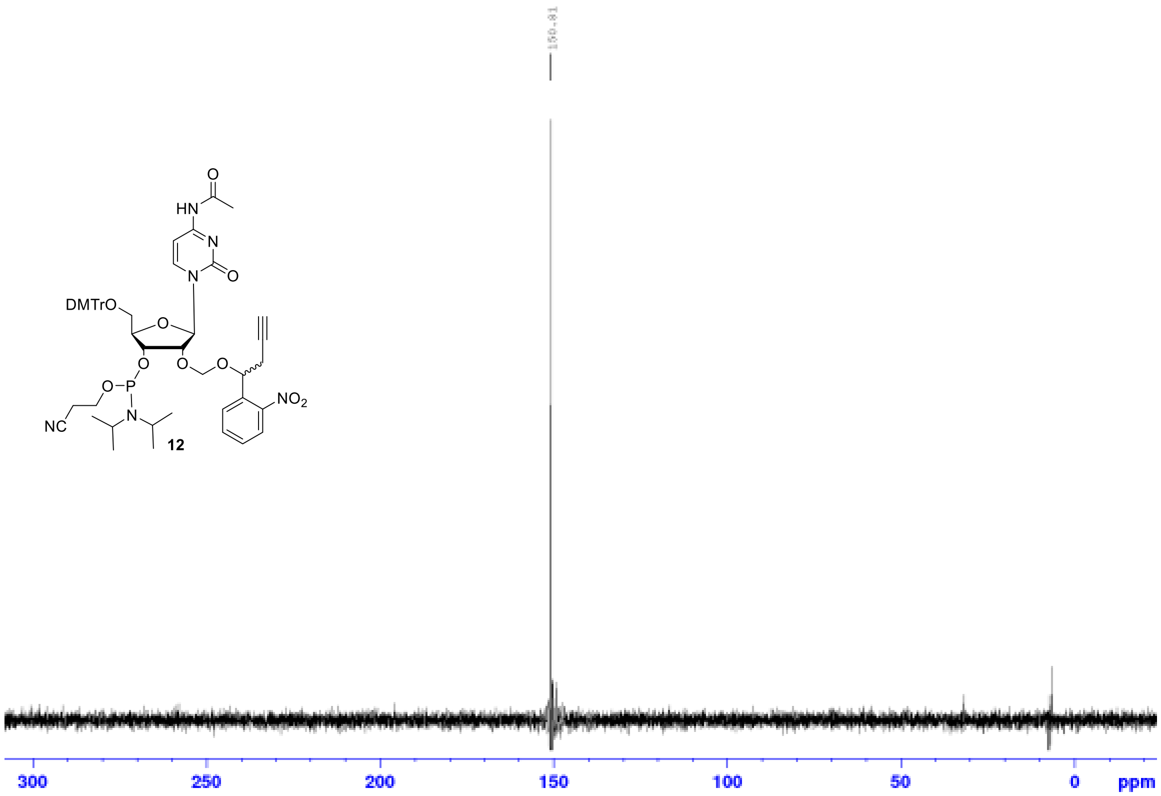

$^{31}P$ NMR of compound **12**

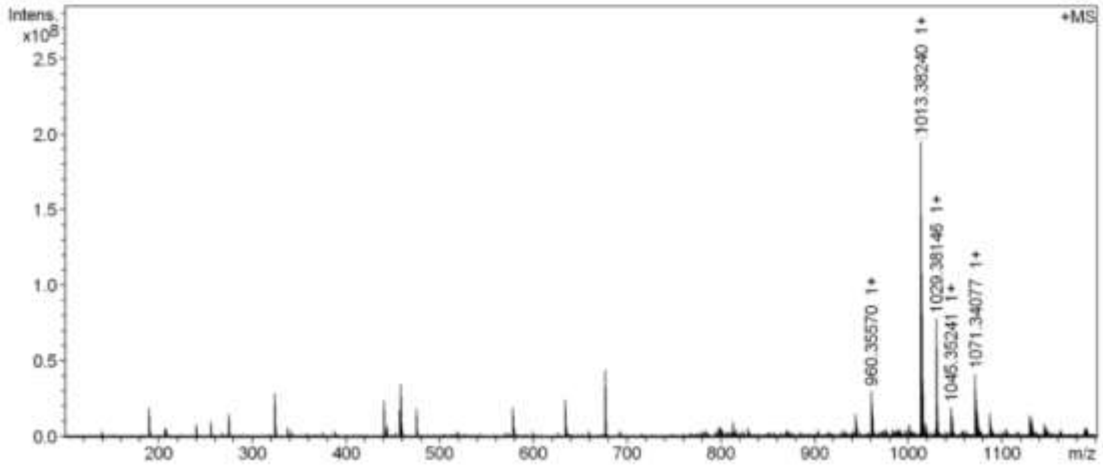


MS spectrum of compound **12**

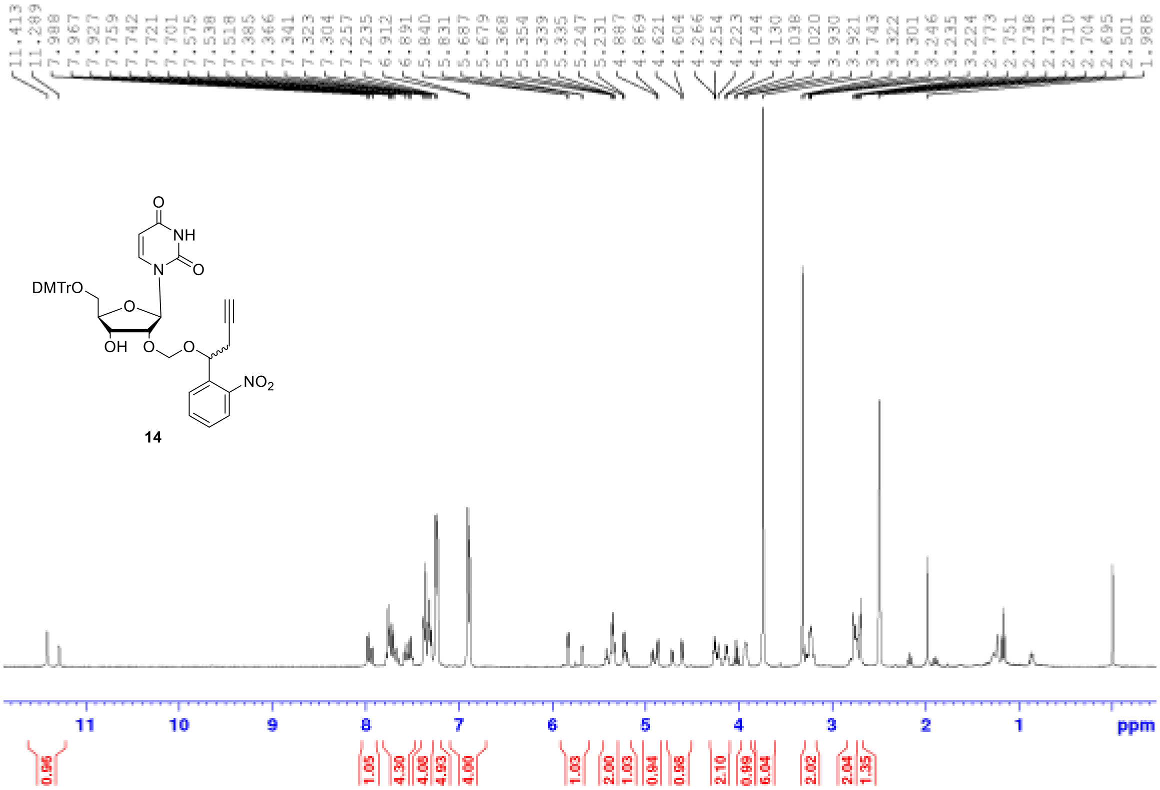


$^1$H NMR of compound **14**

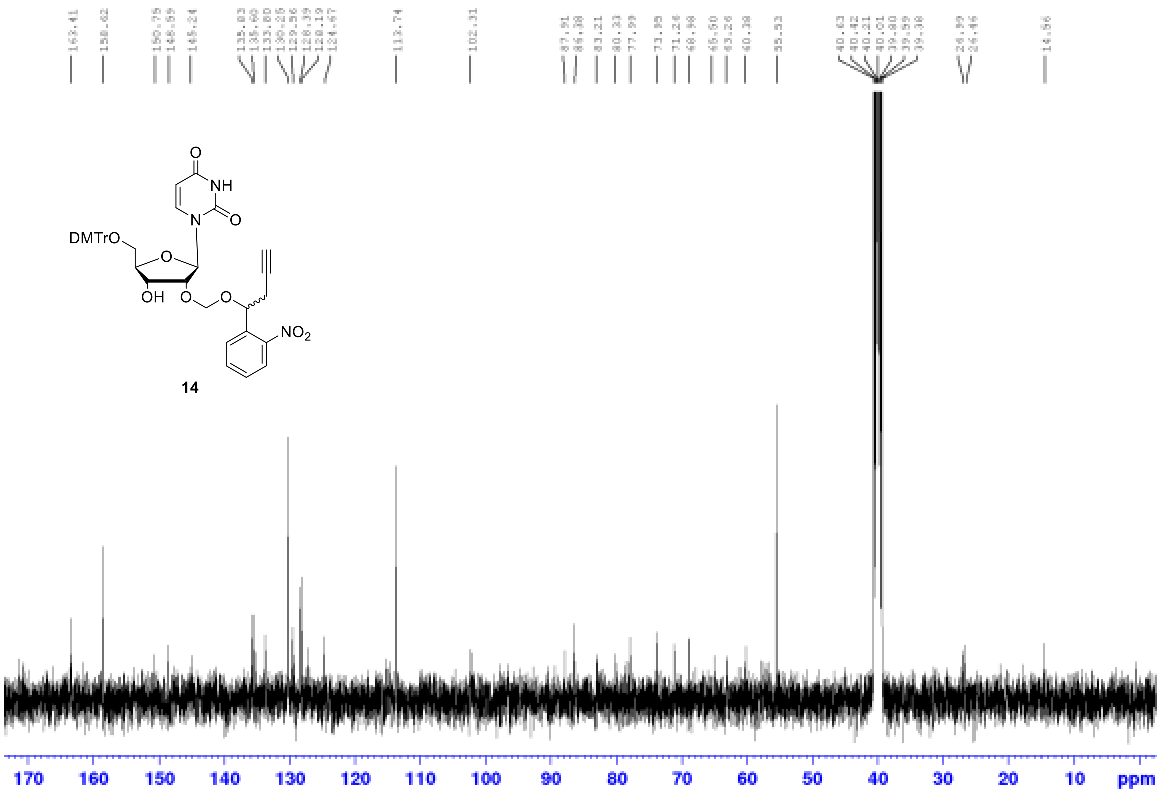

$^{13}C$ NMR of compound **14**

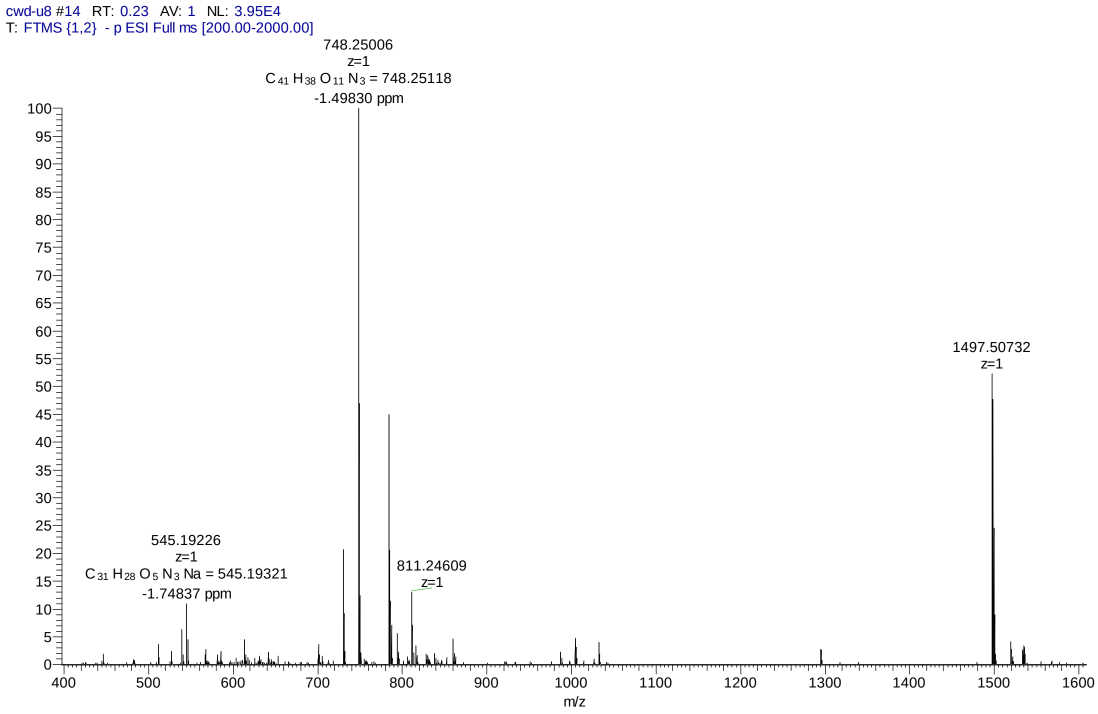


MS spectrum of compound **14**

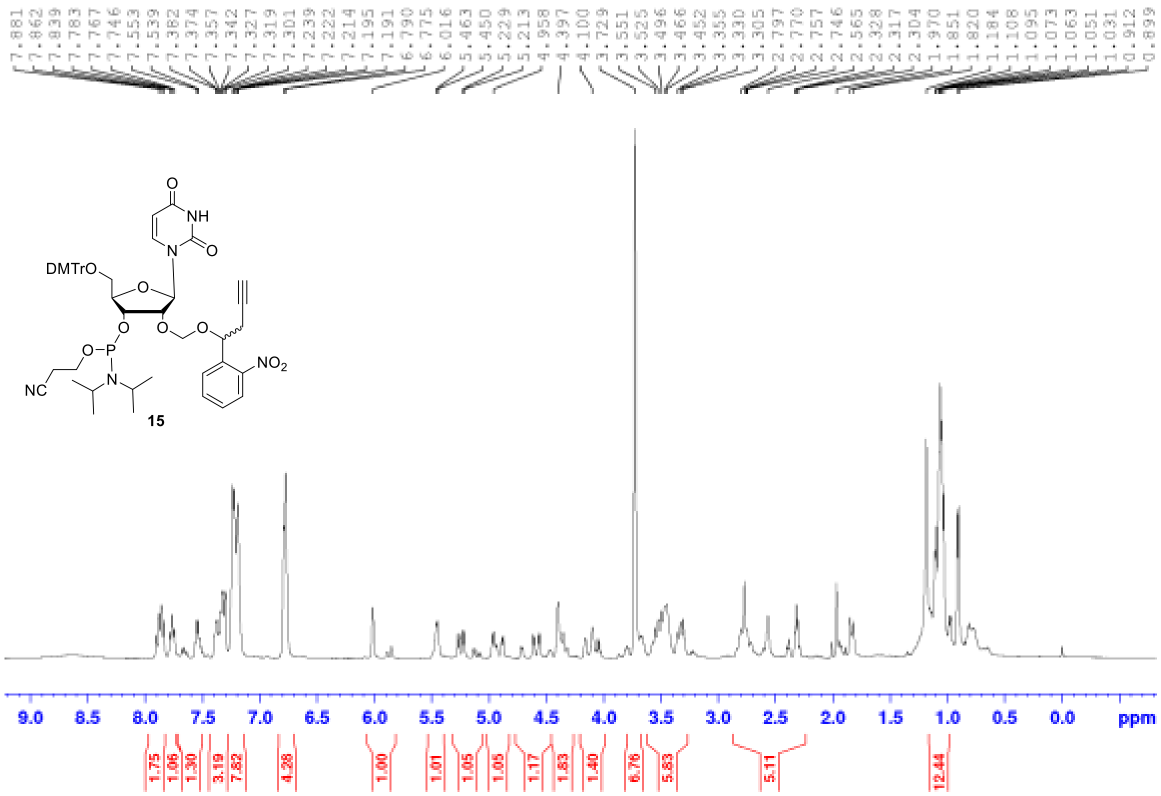


$^{1}$H NMR of compound **15**

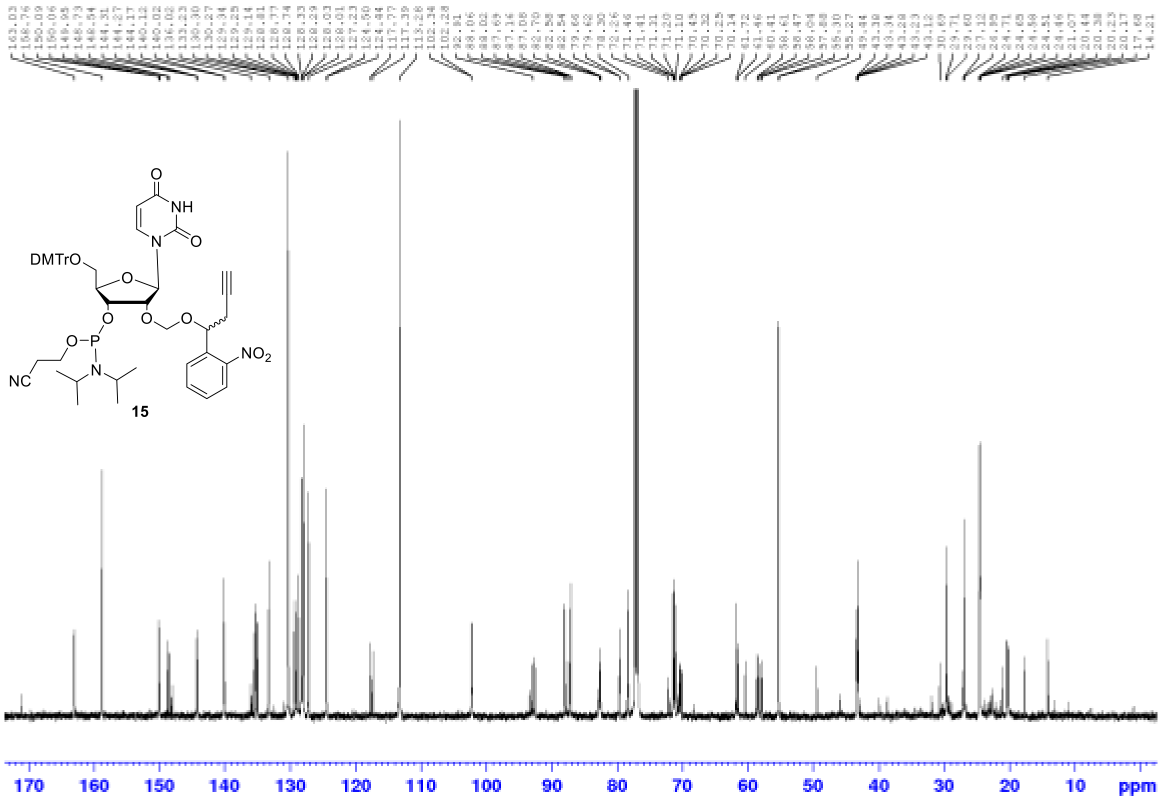


$^{13}$C NMR of compound **15**

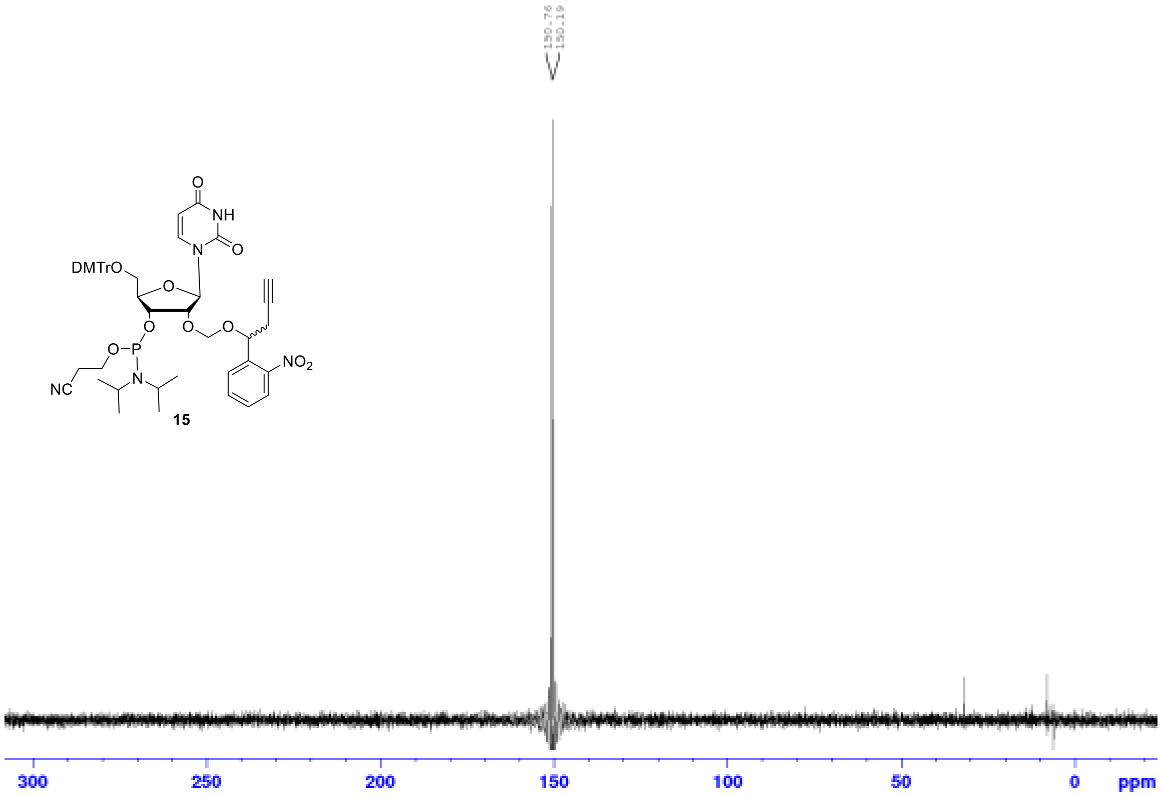


$^{31}$P NMR of compound **15**

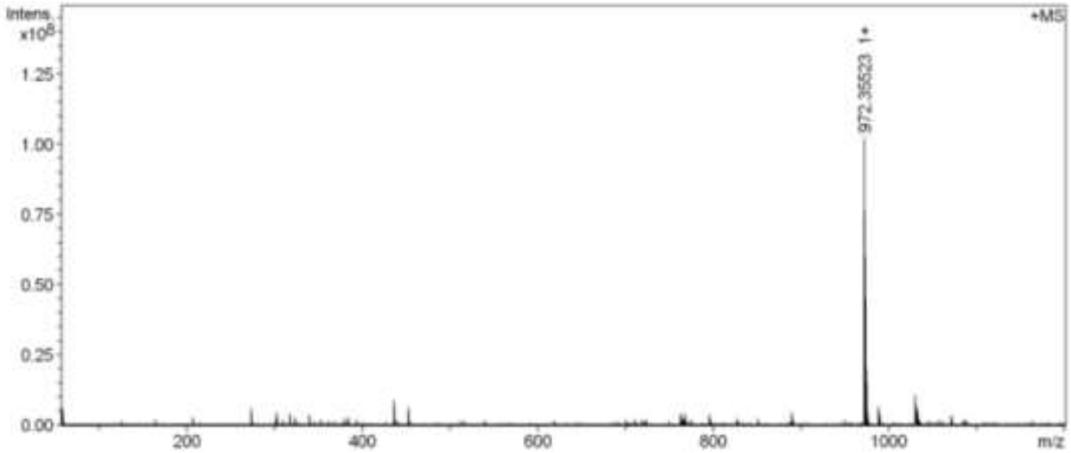


MS spectrum of compound **15**
